# MitoAtlas: a Domain-Resolved Spatial Map of the Human Mitochondrial Proteome

**DOI:** 10.64898/2026.04.03.716438

**Authors:** Jiwoong Kang, Sanghee Shin, Chulhwan Kwak, Song-Yi Lee, Minkyo Jung, Hanseul Lee, Myeong-Gyun Kang, Jiho Sim, Seung-Jae V. Lee, Ji Young Mun, Jong-Seo Kim, Hyun-Woo Rhee

## Abstract

Mitochondrial function depends on the precise sub-organellar localization and topological orientation of its proteome. However, mapping this architecture with domain-level precision across the double-membrane system has remained challenging. Here, we present MitoAtlas, a spatial systems biology resource built on SR-PL (super-resolution proximity labeling) that resolves the sub-mitochondrial architecture of 861 proteins at domain-level resolution. By deploying 42 spatially restricted APEX2 and BioID/BioID2 baits across mitochondrial compartments and cytosol in HEK293T-REx cells, we mapped 13,440 unique proximity-labeled sites. Integrating high-density spatial data with a machine-learning classification pipeline and AlphaFold structural modeling, we resolved membrane topologies for 196 transmembrane proteins and assigned sub-mitochondrial localizations to 160 previously unannotated mitochondrial proteins absent from MitoCarta3.0, including MINPP1 as an intermembrane space resident. Coupling bait-specific proximity labeling signatures with AlphaFold structure prediction and metabolite docking identified 47 protein–protein interactions, including the LETM1–Citrate Synthase complex, and 155 protein–metabolite interactions. MitoAtlas is available at mitoatlas.org as a domain-resolution spatial map of mitochondrial architecture and metabolism.

## INTRODUCTION

The mitochondrion is defined by its sophisticated double-membrane architecture, which partitions the organelle into distinct physiological environments: the outer mitochondrial membrane (OMM), the intermembrane space (IMS), the inner mitochondrial membrane (IMM), and the matrix. This compartmentalization is not merely structural; it is the foundation for specialized biological processes. For example, LYR domains facilitate respiratory complex assembly in the matrix, while Bcl-2 homology (BH) domains at the OMM mediate apoptotic signaling.^1,3,4,8^ Consequently, mapping the mitochondrial proteome with domain-level precision is essential for deciphering the spatial logic of mitochondrial metabolism and regulation.

Despite extensive efforts to catalog mitochondrial proteins, our current spatial understanding remains incomplete.^11^ MitoCarta3.0, which catalogs over 1,100 mitochondrial proteins, still assigns 90 of them to ambiguous “membrane” or “unknown” categories without sub-compartment resolution. Furthermore, over 400 proteins lack precise topological classifications, leaving their functional orientation across the IMM or OMM unresolved. Traditional biochemical methods for topology mapping often struggle with high-throughput scaling and are susceptible to methodological artifacts, leading to long-standing topological errors for critical proteins such as RTN4IP1^9^ and EXD2.^5,6^ Similarly, the topology of LETM1 remains a subject of active debate, complicating our understanding of its role in ion transport and mitochondrial morphology.^7^ Such errors fundamentally hinder the modeling of protein-protein interaction networks and the spatial distribution of functional domains across the double-membrane system. Addressing these challenges requires a spatial systems biology approach: one that integrates proximity labeling data from multiple spatially defined baits within a unified computational framework, rather than relying on binary comparisons between individual compartments.

Proximity labeling (PL) coupled with mass spectrometry has emerged as a powerful tool for mapping sub-mitochondrial architectures in live cells.^2–16^ However, conventional PL workflows often suffer from high false-positive rates due to the detection of non-specific biotinylation or background contaminants. To overcome these limitations, we previously developed super-resolution proximity labeling (SR-PL).^10,17,18^ By focusing exclusively on biotinylated peptides and their specific modification sites, SR-PL provides high spatial accuracy. While our initial efforts with a small number of APEX2 baits validated the precision of this approach, a comprehensive, organelle-wide map required significantly greater proteome coverage. Recent advances in AI-driven protein structure prediction, particularly AlphaFold, now offer the potential to cross-validate and refine such spatial assignments at domain-level resolution by integrating high-confidence structural models with proximity labeling data.

Here, we describe an expanded SR-PL workflow integrating both APEX2 and BioID enzymes across 42 mitochondrial baits and characterize the sequence selectivity and steric constraints underlying each labeling chemistry. A two-stage classification pipeline assigns sub-mitochondrial localization and membrane topology from multi-bait intensity profiles. This framework reveals a substantial cohort of mitochondrial proteins absent from existing databases, whose identities we validate through imaging, biochemistry, and functional assays. Systematic topology mapping across both mitochondrial membranes uncovers unexpected structural features, including a novel β-barrel architecture. Integration of site-resolved labeling data with AlphaFold-Multimer and metabolite docking further enables prediction of protein–protein and protein–metabolite interactions. All spatial annotations, topology maps, and structural predictions are accessible through MitoAtlas.org.

Altogether, MitoAtlas constitutes the most detailed sub-mitochondrial proteome map to date. Unlike existing databases that classify proteins into broad compartmental categories, it provides domain-level spatial resolution through systems-level integration of multi-bait proximity labeling, enabling membrane topology determination and functional domain mapping inaccessible to fractionation-based approaches. Across 861 proteins and 13,440 uniquely labeled sites, we assigned precise sub-mitochondrial localizations to 160 previously unannotated proteins and proposed 47 protein–protein and 155 protein–metabolite interactions, including the LETM1–Citrate Synthase complex.

## RESULTS

### Multiplexed Baits and Orthogonal Labeling Chemistries Dramatically Expand Mitochondrial Proteome Coverage

To achieve comprehensive organellar coverage, we expanded our proximity labeling (SR-PL) toolkit, building upon the foundational framework established in our previous work.^13,14,18^ We utilized two enzymatic systems with distinct chemistries: APEX2^19^, which generates biotin-phenoxyl radicals that label tyrosine residues primarily through contact-dependent labeling within ∼10 nm, with low-level diffusive labeling extending further^20,21^, and BioID/BioID2, which produces biotin-AMP esters that modify lysine residues (radius ∼10 nm)^22^ (**Figure 1A**).

**Figure 1.**
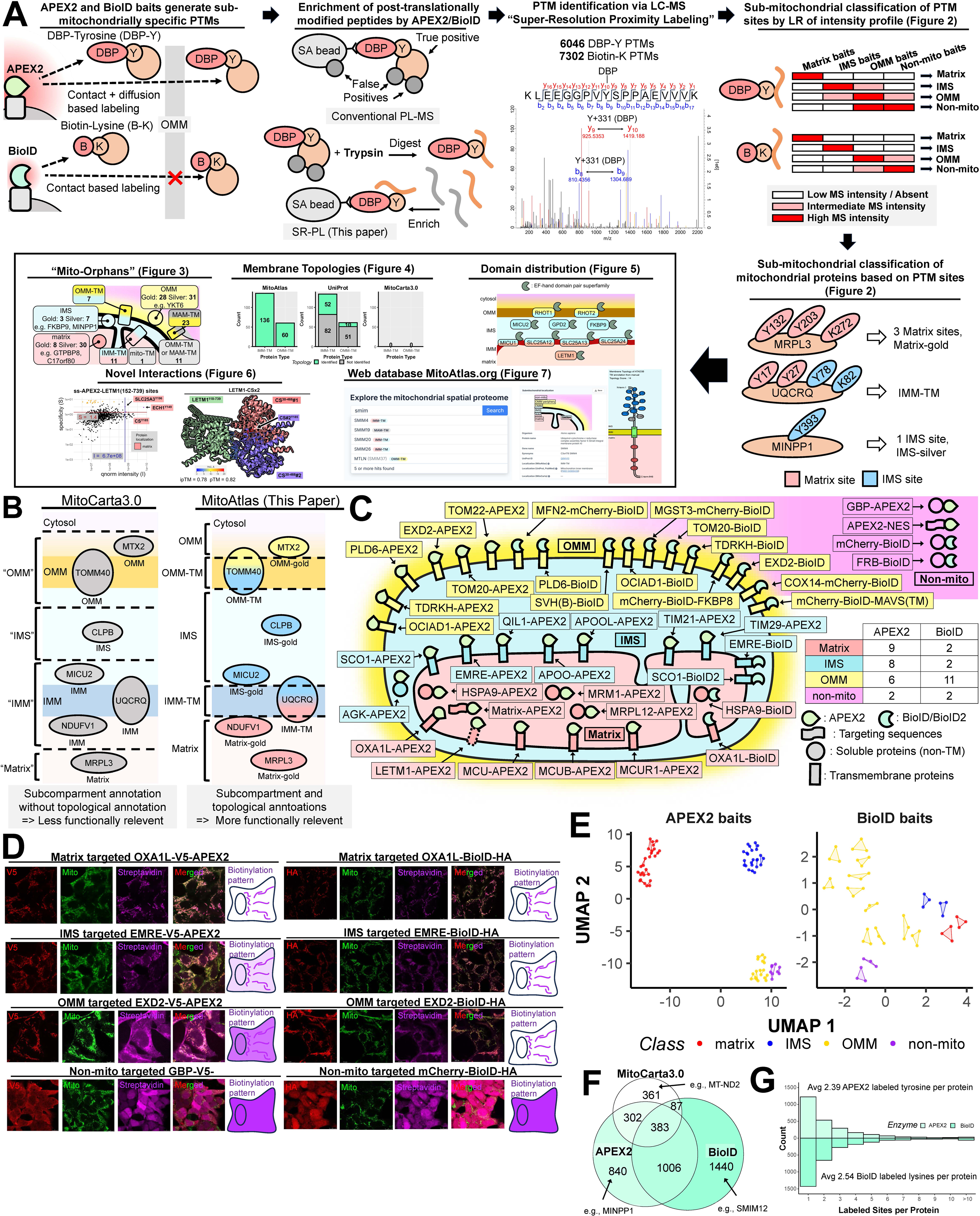
Construction of MitoAtlas via Multiplexed SR-PL Mapping (see also Figure S1 and Table S1). (A) Overview of the MitoAtlas workflow. APEX2 generates biotin-phenoxyl radicals that label tyrosine residues through contact-dependent labeling (radius ∼10 nm), while BioID/BioID2 produces biotin-AMP esters that modify lysine residues (radius ∼10 nm). Proximity-labeled peptides are enriched by streptavidin pulldown and identified via LC-MS/MS as desthiobiotin-phenol (DBP)-modified tyrosines or biotin-modified lysines (Super-Resolution Proximity Labeling, SR-PL). Site-specific MS intensity profiles across matrix, IMS, OMM, and non-mitochondrial baits are then classified by logistic regression (LR) to assign sub-mitochondrial localizations, enabling downstream analyses of membrane topologies (Figure 3), domain distributions (Figure 5), mito-orphan characterization (Figure 4), protein–protein/metabolite interactions (Figure 6), and the MitoAtlas web database (Figure 7). All data are accessible through the MitoAtlas web database (mitoatlas.org). Representative examples show site-level classification for MRPL3 (Matrix-gold), UQCRQ (IMM-TM), and MINPP1 (IMS-silver). (B) Comparison of sub-mitochondrial annotation resolution between MitoCarta3.0 and MitoAtlas. MitoCarta3.0 assigns proteins to broad categories (e.g., “OMM,” “IMS,” “IMM,” “Matrix”) without topological detail, whereas MitoAtlas provides refined classifications including membrane-spanning orientation (e.g., OMM-TM) and confidence tiers (Gold/Silver), yielding more functionally informative annotations. (C) Sub-mitochondrial distribution of the 42-bait library across the four mitochondrial compartments. APEX2 fusions (filled circles) were preferentially deployed to the matrix (9 baits) and IMS (8 baits) to leverage high labeling efficiency in luminal compartments, while BioID/BioID2 fusions (open circles) were primarily targeted to the OMM (11 baits) for confined, neighborhood-specific labeling. Soluble proteins are depicted as circles and transmembrane proteins as rectangles. Colored arrowheads indicate targeting sequences. (D) Validation of bait localization and enzymatic activity. Confocal immunofluorescence microscopy images show colocalization of representative APEX2 (V5-tagged) and BioID/BioID2 (HA-tagged) baits with a mitochondrial marker, and streptavidin staining confirms biotinylation activity. Biotinylation pattern images show the spatial distribution of labeling for matrix-targeted (OXA1L), IMS-targeted (EMRE), OMM-targeted (EXD2), and non-mitochondrial (GBP, mCherry-BioID) baits. (E) UMAP analysis of biological replicates from APEX2 baits (left) and BioID baits (right). Each bait is represented by colored dots: matrix (red), IMS (blue), OMM (gold), and non-mitochondrial (purple). Shaded polygons connect biological replicates of the same bait. (F) Venn diagram showing the overlap between APEX2- and BioID-identified proteins and MitoCarta3.0. APEX2 and BioID together identified 861 mitochondrial proteins, with 383 detected by both enzyme systems. Examples of uniquely identified proteins are shown (e.g., MINPP1 by APEX2 only; SMIM12 by BioID only). (G) Distribution of the number of labeled sites per protein for APEX2 (DBP-modified tyrosines; average 2.39 sites per protein) and BioID (biotin-modified lysines; average 2.54 sites per protein).

We generated 42 stable Flp-In 293T-REx (hereafter HEK293T-REx) cell lines expressing 38 sub-mitochondrial and four non-mitochondrial baits. We strategically biased APEX2 toward the matrix and IMS (17 baits) to leverage its high labeling efficiency in dense compartments. Notably, we previously observed that when APEX2 is fused to OMM transmembrane (TM) proteins such as EXD2, the resulting biotinylation pattern is highly diffusive and cytosolic, raising concerns about the reliability of APEX2 for OMM proteomic analysis.^6^ To circumvent this limitation, BioID was primarily deployed to the OMM (11 baits), where its shorter labeling radius produces more confined, neighborhood-specific labeling that faithfully captures the local OMM protein environment (**Figure 1C**).^6,20,22^ Bait localization was confirmed via confocal microscopy and transmission electron microscopy (TEM) combined with 3,3’-diaminobenzidine (DAB) polymerization, while enzymatic activity was validated by streptavidin-conjugated horseradish peroxidase (Streptavidin-HRP) western blotting (**Figures 1D, S1A, S1B, S1C, and S1D**).

LC-MS/MS analysis of 129 biological replicates following SR-PL yielded 13,440 unique proximity-labeled sites (6,046 desthiobiotin-phenol (DBP)-labeled Tyr; 7,394 biotin-labeled Lys) across 4,058 proteins (**Figure 1A**). This is a 19-fold increase compared to our prior SR-PL study.^18^ UMAP analysis of biological replicates confirmed that samples clustered by sub-mitochondrial compartment for both APEX2 and BioID baits, demonstrating the reproducibility and spatial specificity of the labeling approach (**Figure 1E**). Together, APEX2 and BioID detected 772 MitoCarta3.0 proteins, with 383 identified by both enzyme systems (**Figure 1F**). On average, APEX2 labeled 2.39 tyrosine residues per protein, while BioID labeled 2.54 lysine residues per protein (**Figure 1G**).

Critically, the identification of multiple labeled sites within the same protein not only increases the confidence of sub-mitochondrial localization assignments but also provides the spatial resolution necessary to determine the membrane topology of transmembrane proteins; that is, which functional domains are exposed to the matrix, IMS, or cytosol across the lipid bilayer. This site-level resolution represents a fundamental advance over existing resources such as MitoCarta3.0, which assigns proteins to broad sub-mitochondrial categories (e.g., "OMM," "IMM," "IMS," "Matrix") without topological detail. By contrast, MitoAtlas provides refined classifications including membrane-spanning orientation and confidence tiers (Gold/Silver), yielding more functionally informative annotations (**Figure 1B**).

The value of our multiplexed approach is further evidenced by the expanded proteome coverage achieved using multiple baits within the same sub-organellar region. For instance, combining data from mitochondrial targeting sequence (MTS)-APEX2, HSPA9-APEX2, and MRM1-APEX2 revealed a vast array of distinct modified peptides, significantly increasing the depth of the matrix proteome (**Figure S1E**). When the same bait proteins were used with both systems (e.g., TOM20, EXD2, and TDRKH), BioID generated a higher proportion of non-overlapping modified peptides compared to APEX2 (**Figure S1E**). This is consistent with the distinct labeling mechanisms of the two enzymes: APEX2 operates through both contact-dependent and diffusive labeling, capturing a broader spatial neighborhood, whereas BioID labels exclusively through a contact-dependent mechanism, detecting only direct interaction partners.^20^ This dual-enzyme strategy thus provides both the breadth required for organelle-wide mapping and the specificity needed to define local protein environments.

We observed that APEX2 and BioID display distinct sequence preferences and steric limitations, distinct from previously reported N-hydroxysuccinimide ester modified sites (Figure S1F).^23^ Sequence motif analysis revealed that glycine (Gly) is frequently adjacent (±1) to APEX2-modified tyrosines, likely because its small side chain minimizes steric interference with the reactive phenol radical. The sequence logo also revealed that electron-rich amino acids capable of radical quenching, including Cys, Trp, Met, Arg, Phe, Leu, Tyr, and His, were negatively enriched at positions flanking the labeled tyrosine (Figure S1F). Among these, cysteine exhibited the strongest depletion, consistent with the high reactivity of sulfur-containing and aromatic amino acid side chains toward electrophilic radicals (k = 10⁹–10¹⁰ M⁻ ¹s⁻ ¹, compared with ∼10⁷ M⁻ ¹s⁻ ¹ for aliphatic residues such as glycine).^24^ These flanking residues likely intercept the short-lived phenoxyl radical before it forms a stable covalent adduct at the target tyrosine, providing mechanistic insight into the sequence selectivity of APEX2 labeling. In contrast, BioID modification sites favored branched amino acids (Val, Leu, Ile) at the +1 position, consistent with the substrate preference of the ancestral E. coli BirA enzyme, supporting its contact-dependent labeling mechanism.^20,25^

To further explore the physical constraints of labeling, we analyzed the correlation between solvent-accessible surface area (SASA) and peptide detection using AlphaFold2 structures. APEX2 showed an optimal probe radius near that of water (1.1–1.2 Å), suggesting it can probe deep into small protein clefts. In contrast, BioID exhibited a much larger optimal radius (11.5 Å), indicating that the bulkiness of the biotin-AMP molecule restricts labeling to more exposed surface residues (**Figures S1G and S1H**). These distinct labeling constraints have direct consequences for MitoAtlas: APEX2’s small effective probe radius enables it to reach buried residues critical for resolving transmembrane topology (**Figure 3**), while BioID’s contact-dependent mechanism, confirmed here by the retained BirA substrate preference, underlies the identification of specific bait–prey interactions (**Figure 6**).

**Figure 2.**
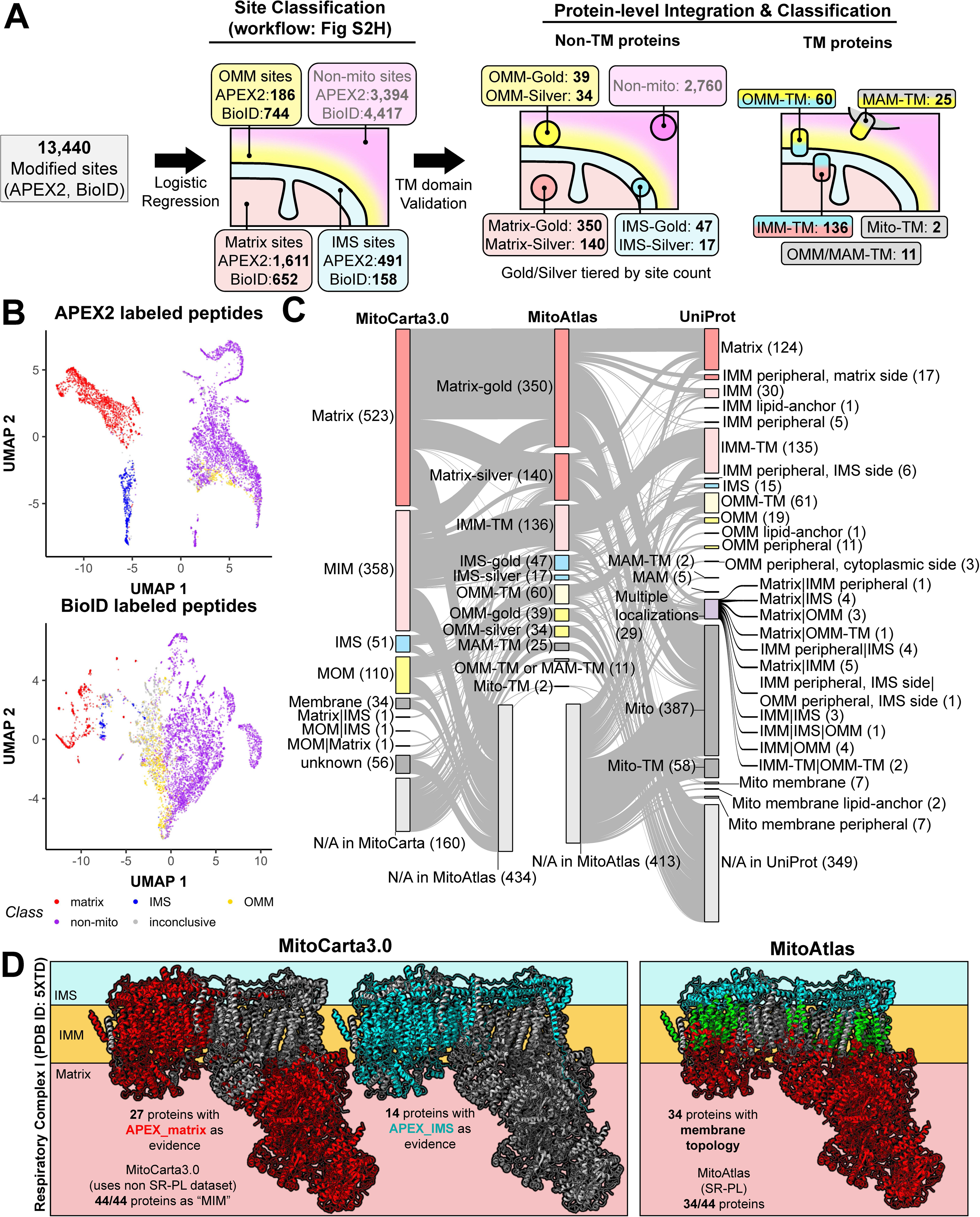
**A Two-Step Classification Pipeline for Sub-mitochondrial Proteome Mapping (see also Figure S2 and Tables S2 and S3).** (A) Schematic overview of the two-step classification pipeline. In Step 1 (Site Classification), 13,440 modified sites from APEX2 and BioID experiments are classified into four sub-mitochondrial compartments (Matrix, IMS, OMM, and Non-mitochondrial) by logistic regression (workflow detailed in Figure S2H). In Step 2 (Protein-level Integration and Classification), site-level assignments are aggregated per protein. Non-TM proteins are assigned gold or silver confidence tiers based on the number of supporting sites, while TM proteins undergo additional topology validation to resolve membrane orientation (IMM-TM, OMM-TM, MAM-TM). (B) UMAP visualization of APEX2-labeled (top) and BioID-labeled (bottom) peptides classified by the site-level logistic regression model. Each point represents a single modified site, colored by its predicted sub-mitochondrial compartment (Matrix, red; IMS, blue; OMM, yellow; Non-mitochondrial, purple; Inconclusive, gray). Distinct clustering of compartment-specific sites demonstrates the discriminative power of the multi-bait labeling profiles. (C) Sankey diagrams illustrating the flow of protein annotations across MitoCarta3.0, MitoAtlas, and UniProt. Flows highlight how MitoAtlas resolves previously ambiguous categories (e.g., “MIM,” “Membrane,” “Unknown”) from MitoCarta3.0 into precise sub-mitochondrial localizations with topological detail, and how these assignments compare with UniProt annotations. Notably, MitoAtlas classifies 160 proteins not annotated in MitoCarta3.0 and provides topology information for proteins listed only broadly as “Mitochondrion” in UniProt. (D) Comparative spatial mapping of respiratory Complex I subunits. MitoCarta3.0 (left) assigns all 44 subunits broadly as “MIM” using non-SR-PL data. MitoAtlas (right) resolves 34 of 44 subunits with membrane topology information, distinguishing Matrix-facing, IMS-facing, and IMM-TM components consistent with high-resolution cryo-EM structural data.

**Figure 3.**
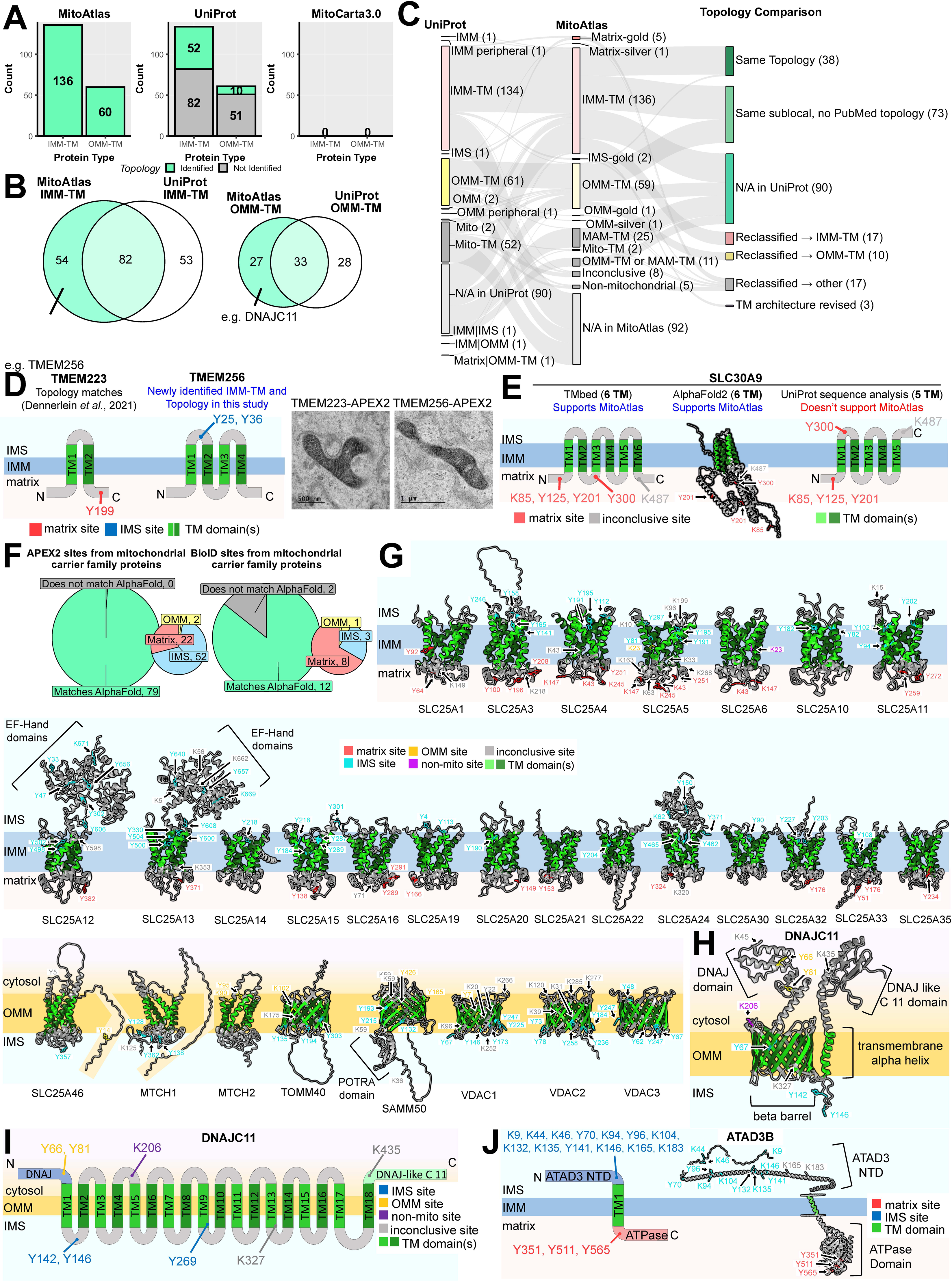
**High-Resolution Membrane Topology and Domain Architecture of the Mitochondrial Proteome (see also Figure S3 and Tables S3 and S4).** (A) Comparison of the number of IMM-TM and OMM-TM proteins with identified topologies across databases. MitoAtlas (left) resolves the membrane topologies of all 136 IMM-TM and 60 OMM-TM proteins detected. UniProt (middle) contains curated topological annotations for a subset of entries. MitoCarta3.0 (right) lacks topological orientation data (0/0). Other databases without topology information (Human Cell Map, MitoCoP) are also indicated. (B) Venn diagrams illustrating the overlap and novel identification of IMM-TM (left) and OMM-TM (right) proteins in MitoAtlas relative to UniProt. (C) Sankey plot depicting the consistency and discrepancies in membrane topology assignments between MitoAtlas and UniProt experimental records. (D) Validation of IMM-TM topologies. (Left): Proposed membrane topologies of TMEM223 and TMEM256 at the IMM based on SR-PL data. (Right): Validation of C-terminal orientation via APEX2-enhanced electron microscopy (APEX-EM) imaging. (E) Comparative analysis of SLC30A9 TM domain count and orientation as determined by MitoAtlas integrated with TMbed (left), MitoAtlas integrated with AlphaFold2 (AF2) structures (middle), and current UniProt annotations (right). (F) Structural validation of mitochondrial carriers. Concordance between SR-PL-derived labeling sites and AF2-predicted TM domain positions for the mitochondrial carrier protein family (SLC25). (G) Topology of multi-pass and β-barrel proteins. Proposed membrane topologies for 24 SLC25 family proteins and 5 OMM β-barrel proteins identified in MitoAtlas. (H) AF2-predicted structure of DNAJC11 highlighting the 17-stranded β-barrel TM domain and corresponding SR-PL labeling sites. (I) Schematic representation of the DNAJC11 domain organization and sub-mitochondrial localization based on MitoAtlas SR-PL data. (J) Domain architecture and membrane topology of ATAD3B. (Left) Schematic of ATAD3B domain organization and sub-mitochondrial localization based on MitoAtlas SR-PL data, with the N-terminal domain facing the IMS and the C-terminal ATPase domain in the matrix. (Right) Membranome-adjusted AlphaFold2 structure^64^ with SR-PL site annotations mapped onto the 3D model, confirming the IMM-TM topology.

### A Two-Step Classification Pipeline for Sub-mitochondrial Proteome Mapping

The high-density labeling profiles generated by our 42-bait library provided a unique fingerprint for each proximity-labeled site. We observed that MS intensities for sites within proteins such as MRPL22 (Matrix), MICU2 (IMS), FKBP8 (OMM), and ACTB (non-mitochondrial) exhibited compartment-specific enrichment patterns across the bait panel, suggesting that these signatures could be used for systematic spatial classification (**Figure S2A**).

To build a classification framework, we developed a two-step computational pipeline (**Figure 2A**). In Step 1 (Site Classification), we evaluated six supervised learning models and selected Logistic Regression (LR) for its accuracy, interpretability, and tunable probability thresholds (**Figure S2B**). Separate models were trained on APEX2 and BioID datasets, with probability thresholds optimized to maintain an overall FDR below 10% (**Figures S2C and S2D**). UMAP visualization of the classified sites confirmed distinct compartment-specific clustering for both APEX2- and BioID-labeled peptides (**Figure 2B**). By integrating data from all 42 baits simultaneously, MitoAtlas outperforms conventional binary enrichment analysis based on a single bait–control comparison. We benchmarked this on the most challenging classification task: discriminating OMM from non-mitochondrial sites, where OMM proteins face the cytosol and share the labeling environment with cytosolic background. MitoAtlas reduced classification error by 43% for APEX2 (AUC = 0.923 vs. 0.866 for TOM20-APEX2 vs. APEX2-NES) and by 58% for BioID (AUC = 0.976 vs. 0.943 for TOM20-BioID vs. mCherry-BioID) (**Figure S2E**).

In Step 2 (Protein Classification), site-level P1 outputs were aggregated into protein-level features and classified by a second LR model (**Figures 2A** **and S2H**). For non-TM proteins, we implemented a tiered “Gold” and “Silver” hierarchy reflecting the depth of evidence for each assignment. For Matrix and IMS proteins, those with a single labeled site were designated Silver and those with two or more sites as Gold. Recognizing the higher risk of cytosolic contamination at the outer membrane, we applied more stringent filters for OMM proteins: a minimum of two sites for Silver and three or more for Gold. This pipeline successfully categorized 627 non-TM proteins, including Matrix-Gold/Silver (490), IMS-Gold/Silver (64), and OMM-Gold/Silver (73), while filtering out 2,760 non-mitochondrial proteins (**Figures 2A, S2F, and S2H**). At the selected P2 thresholds, protein-level FDR remained below 5% for Matrix, IMS, and non-mitochondrial compartments; the higher FDR for OMM (13%) reflects the inherent difficulty of discriminating OMM from cytosolic proteins, which motivated the more stringent Gold/Silver site-count requirements described above (**Figure S2G**).

An additional ∼1,700 labeled sites did not meet the multi-bait consensus threshold for confident compartment assignment. These inconclusive sites were predominantly BioID-derived lysine modifications (1,323 of 7,394 BioID sites, 17.9%, versus 364 of 6,046 APEX2 sites, 6.0%). Among these inconclusive sites, 96% (APEX2) and 90% (BioID) were detected by at least one mitochondrial bait, indicating genuine mitochondria-proximal labeling events rather than cytosolic background (**Figure S2K**). Within this group, 37% of BioID cases were detected by only a single bait, suggesting that BioID retains a contact-dependent component of its ancestral BirA mechanism, preferentially capturing direct protein–protein interactions. We retained all inconclusive sites with full probability scores in the MitoAtlas database, as they likely reflect contact-dependent bait–prey interactions and represent candidate interactors amenable to targeted validation (see **Figure 6**).

For transmembrane (TM) proteins, we integrated site-level P1 values with transmembrane domain predictions from four independent sources: UniProt curated annotations, DeepTMHMM (sequence-based deep learning), TMbed (protein language model embeddings), and TmDet 4.0 (AlphaFold2 structure-based detection)^26–30^ to assign membrane topology (**Figures 2A** **and S2H**). This workflow uses a topology scoring system that evaluates the concordance between site localizations and predicted TM domain boundaries, as illustrated for ATAD3B, an IMM-TM protein whose mixed Matrix and IMS site assignments are reconciled through integration with the DeepTMHMM-predicted topology and AlphaFold structure (**Figure S2I**). This pipeline successfully assigned membrane topology to 136 IMM-TM, 60 OMM-TM, and 25 MAM-TM proteins, with ambiguous cases classified into “Mito-TM” or “OMM-TM or MAM-TM” categories.

MitoAtlas resolves sub-mitochondrial localizations at a resolution inaccessible to existing databases.^29^ MitoCarta3.0 groups all 44 Complex I subunits as “MIM,” whereas MitoAtlas distinguishes matrix-facing and IMS-facing peripheral subunits from core transmembrane components, with topologies concordant with established cryo-EM structures (**Figure 2D**). More broadly, Sankey diagrams tracing annotation flow across MitoCarta3.0, MitoAtlas, and UniProt show how the pipeline resolves ambiguous categories into precise sub-mitochondrial addresses (**Figure 2C**). In total, MitoAtlas encompasses 861 mitochondrial proteins (matrix/IMM-TM/IMS/OMM-TM/OMM peripheral = 490/136/64/60/73), covering 79% of OXPHOS subunits,^31,32^ 91% of the mitochondrial ribosome,^33^ and 86% of the TOM complex^34^ (**Figure S2J**).

### High-Resolution Mapping of Mitochondrial Membrane Topology and Domain Architecture via SR-PL

Determining the membrane topology and spatial domain architecture of TM proteins is fundamental to understanding their biological functions. Yet current mitochondrial topology information in public databases remains far less resolved than our SR-PL dataset. Although MitoCarta3.0 annotates 469 proteins as localized to the inner (IMM) or outer (OMM) mitochondrial membranes, it provides no topological orientation data.^35^ Similarly, other mitochondrial resources such as MitoCOP (high-confidence mitochondrial proteome from subtractive proteomics) and MitCOM (mitochondrial complexome from native gel profiling in yeast) provide protein inventories and complex compositions but do not resolve membrane topology, as their underlying methodologies were not designed to determine the spatial orientation of individual domains.^2,15,36^

To our knowledge, UniProt is the only database that provides mitochondrial topological information through manual curation of experimental evidence.^29^ As of January 2026, UniProt contains 134 IMM-TM and 61 OMM-TM entries; however, only a fraction of these are supported by specific PubMed references for topology (52 IMM; 10 OMM), reflecting the historical limitations of high-throughput methods (**Figure 3A**). MitoAtlas, by contrast, resolves the membrane topologies of all 136 IMM-TM and 60 OMM-TM proteins in our dataset by leveraging the inherent spatial specificity of SR-PL. Relative to curated UniProt records, MitoAtlas identified 78 previously uncharacterized mitochondrial TM proteins (52 IMM-TM; 26 OMM-TM) (**Figure 3B**). Of the 196 TM proteins identified in MitoAtlas, 38 are in full agreement with curated experimental evidence from UniProt (**Figure 3C**). Beyond these validated proteins, MitoAtlas resolved the topology of newly identified proteins such as TMEM256. Site-specific labeling indicates a topology in which both the N- and C-termini reside within the mitochondrial matrix, an architecture validated by C-terminal APEX2-EM imaging (**Figure 3D**).

However, our SR-PL data reveal several topological conflicts with existing UniProt annotations and sequence-based predictions. For instance, UniProt reports only 5 TM domains for SLC30A9 and SFXN2, an architecture incompatible with our labeling patterns. We instead propose a 6-TM domain model for both SLC30A9 (**Figure 3E**) and SFXN2 (**Figure S3A**), a configuration independently supported by AlphaFold structural modeling. Our data also reassigns TMEM11; previously characterized as an IMM protein with two UniProt-predicted TM domains ^37^, our results indicate it is an OMM-TM protein with a 3-TM architecture, consistent with AlphaFold predictions (**Figure S3B**). Similarly, whereas protease protection assays previously suggested the TMEM126A N-terminus faces the matrix ^38^, our SR-PL results indicate the N-terminus is oriented toward the IMS (**Figure S3C**). These discrepancies underscore the limitations of conventional methodologies. Approaches such as protease protection or antibody-based assays are often conducted under non-physiological conditions or rely on the specificity of reagents that may lead to erroneous assignments. In contrast, our SR-PL strategy conducts proximity labeling in situ within intact living cells, providing a high-resolution, physiologically relevant depiction of the mitochondrial membrane proteome.

Our SR-PL findings demonstrate high concordance with AlphaFold2 (AF2) models: across 29 IMM-TM carrier proteins and 8 multi-pass OMM-TM proteins, 97% of MitoAtlas labeling sites (112/115) matched the expected compartment assignment based on AF2 structures (**Figures 3F-G**). MitoAtlas also provides experimental support for AF2 predictions of novel proteins. For example, UniProt classifies DNAJC11 as a peripheral OMM protein, but MitoAtlas identified both IMS and OMM sites, characterizing it as an OMM-TM protein consistent with an AF2-predicted 17-stranded β-barrel structure harboring an internal pore (**Figures 3H-I, S4E**). Because β-barrel proteins, including VDAC1–3, SAMM50, and TOMM40, are a rare class restricted to the OMM for transport and signaling, our data suggest DNAJC11 is a novel member of this family, with its N-terminal DnaJ domain exposed to the cytosol. Similarly, this integrative approach allowed us to resolve the disputed topology of LETM1.^7^ While initially presumed to be an IMM-TM protein, combining our spatial labeling patterns with AF2 models indicates that LETM1, along with LETMD1^39^ and FOXRED1, is a peripheral matrix-resident protein (**Figure S3F**).

Beyond individual proteins, MitoAtlas enables the mapping of sub-mitochondrial functional domains. For the m-AAA protease AFG3L2^40^, our data (IMS: Y179, Y211; Matrix: K488–Y756) confirm that the M41 FtsH domain localizes to the IMS while the AAA+ ATPase and lid domains reside in the matrix (**Figure 3J**). This domain-resolution mapping, available for all 861 proteins in our dataset, enables molecular-level analysis of mitochondrial function.

The traditional approach of fusing a single proximity labeling enzyme to a target protein can result in mislocalization artifacts. For instance, COX14 was previously assigned as an OMM protein in our single-bait studies^18,41^, but multi-bait MitoAtlas SR-PL profiles and AlphaFold-Multimer predictions support its reclassification as an IMM-TM protein (**Figure S3G**). This misclassification likely arose because C-terminal fusion of proximity labeling enzymes prevented this IMM-TM protein from translocating past the OMM, effectively trapping it on the outer membrane. By integrating data from 42 multiplexed spatial baits and cross-validating with AlphaFold structural models, MitoAtlas successfully identifies and corrects such fusion-induced artifacts, demonstrating how a systems-level, multi-bait framework resolves ambiguities that single-enzyme approaches cannot.

### Expanding the Mitochondrial Map: Characterization of 160 Mito-Orphan Proteins

Through SR-PL, MitoAtlas identified 160 mitochondrial proteins previously unannotated in MitoCarta3.0,^35^ which we termed “Mito-Orphans” (**Figure 4A**). These mito-orphans are distributed across all mitochondrial compartments: 38 matrix, 10 IMS, 11 IMM-TM, 66 OMM (including 7 OMM-TM), 23 MAM-TM, and 12 non-definitive transmembrane proteins (**Figure 4B**).

**Figure 4.**
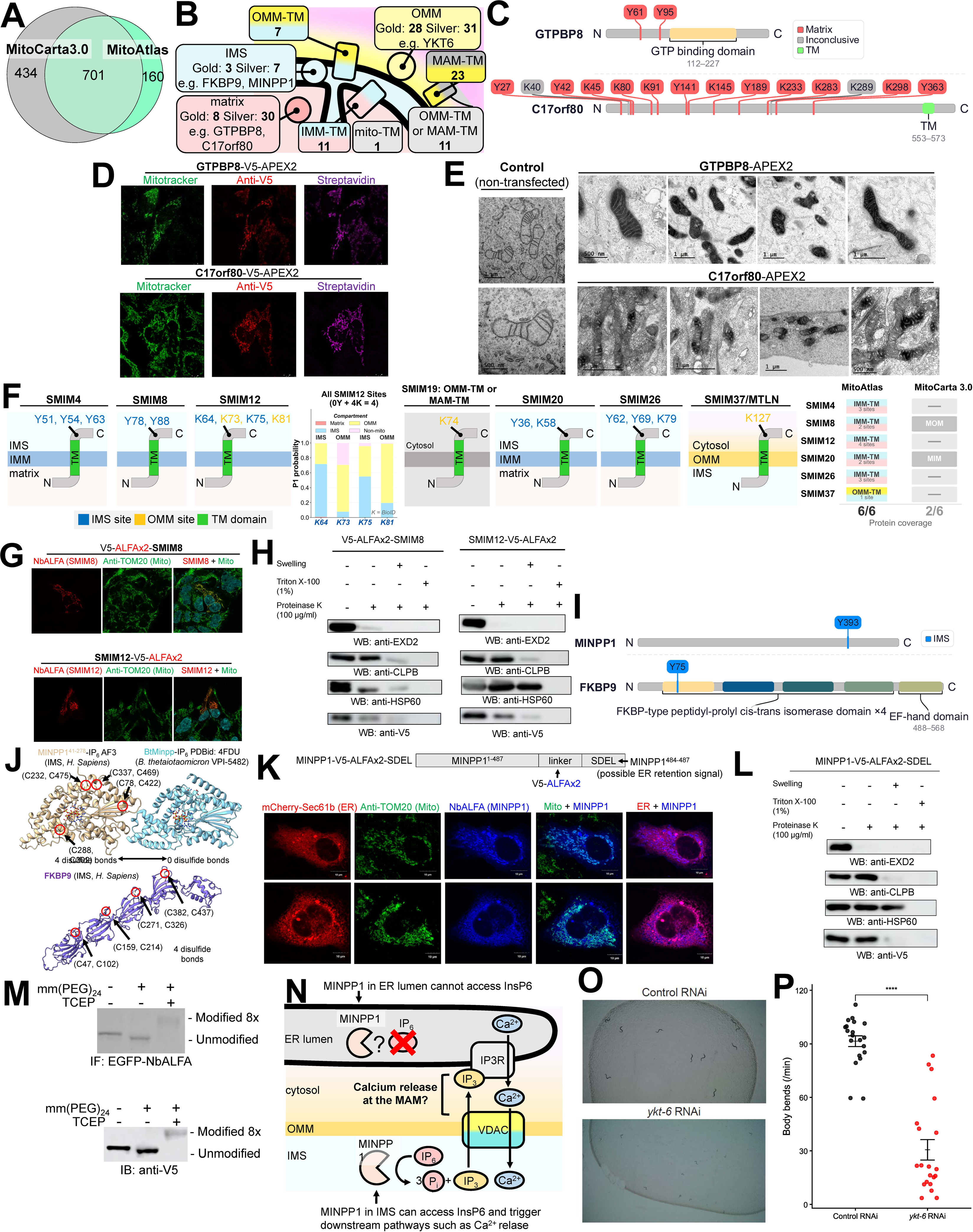
**Expanding the Mitochondrial Map: Characterization of 160 Mito-Orphan Proteins and MINPP1 as an IMS Protein (see also Figure S4 and Table S5).** (A) Venn diagram comparing protein coverage between MitoCarta3.0 and MitoAtlas. MitoAtlas identifies 160 mitochondrial proteins not annotated in MitoCarta3.0, termed “Mito-Orphans.” (B) Sub-mitochondrial distribution of the 160 mito-orphans across compartments: matrix (Gold and Silver), IMS (Gold and Silver), OMM-TM, IMM-TM, MAM-TM, mito-TM, and OMM/MAM-TM categories, with representative examples. (C) SR-PL site maps for two validated matrix mito-orphans, GTPBP8 and C17orf80, showing the distribution of labeled residues and their site-level classifications. Sites are colored by assigned compartment (matrix in red; inconclusive in gray). (D) Confocal immunofluorescence microscopy validating mitochondrial targeting of V5-APEX2-tagged GTPBP8 and C17orf80, co-stained with Mitotracker and streptavidin. (E) Transmission electron microscopy (TEM) of APEX2-mediated DAB polymerization for GTPBP8-APEX2 and C17orf80-APEX2, confirming matrix localization of the C-terminus. Control (non-transfected) cells are shown for comparison. (F) Proposed membrane topologies for SMIM8 and SMIM12 at the IMM, based on SR-PL site assignments. MitoCarta3.0 previously classified SMIM8 as “MOM” (OMM), while SMIM12 was not annotated. Both are reclassified as IMM-TM proteins with N-termini facing the matrix. For SMIM12, site-level P1 classification of four BioID-labeled lysine residues is shown: two sites are assigned to IMS and two to OMM. This mixed assignment pattern is consistent with an IMM-TM topology, as the short IMS-exposed loop of small single-pass TM proteins lies in close proximity to OMM-anchored BioID baits, falling within their ∼10 nm labeling radius. (G) Confocal microscopy validation of SMIM8 and SMIM12 localization using V5-ALFAx2-tagged constructs expressed in HEK293T-REx cells, detected with NbALFA-mScarlet and co-stained with anti-TOM20. (H) Proteinase K protection assay confirming IMM-TM topology of SMIM8 (left, V5-ALFA-SMIM8) and SMIM12 (right, SMIM12-V5-ALFAx2) using isolated mitochondria from HEK293T-REx cells, with the same assay layout as in (L). (I) SR-PL site maps for MINPP1 and FKBP9, two IMS mito-orphans. Each displays a single IMS-classified tyrosine site, with domain architecture shown (histidine phosphatase superfamily for MINPP1; FKBP-type PPIase and EF-hand domains for FKBP9). (J) AlphaFold2/3-predicted structures of MINPP1 and FKBP9, highlighting predicted disulfide bonds consistent with IMS residence where the MIA40/CHCHD4 disulfide relay system operates. Structural comparison with the bacterial homolog BtMinpp (PDB: 4FDU), which lacks disulfide bonds, is shown. (K) Construct design for MINPP1-V5-ALFAx2-SDEL (retaining the ER retention signal) and confocal microscopy in COS-7 cells showing dual localization of MINPP1 to both the ER (mCherry-Sec61b) and mitochondria (anti-TOM20), establishing MINPP1 as a dual-localizing ER lumen/IMS protein. (L) Proteinase K protection assay using isolated mitochondria from MINPP1-V5-ALFAx2-SDEL-expressing HEK293T-REx cells. Digitonin titration shows progressive loss of the MINPP1 signal, consistent with IMS localization. EXD2 (OMM) and HSP60 (matrix) serve as controls. (M) mmPEG24-maleimide modification assay confirming the presence of disulfide bonds in MINPP1. Treatment with the reducing agent TCEP prior to mmPEG24 labeling reveals a mobility shift, demonstrating that MINPP1 contains reducible disulfide bonds consistent with proper folding in the IMS environment. Detection by both immunofluorescence (IF: EGFP-NbALFA) and immunoblotting (IB: anti-V5). (N) Proposed model for MINPP1 function. In the ER lumen, MINPP1 cannot access its substrate InsP6. At ER–mitochondria contact sites, IMS-localized MINPP1 can hydrolyze InsP6 to generate IP3, which may promote calcium release through IP3R and subsequent mitochondrial calcium uptake via VDAC and MCU. (O) Representative images of *C. elegans* body morphology following RNAi knockdown of *ykt-6*, a mito-orphan identified as a novel OMM protein. (P) Quantification of body-bending frequency in *C. elegans* upon *ykt-6* RNAi knockdown, showing a significant reduction in motility compared to control RNAi (****p < 0.0001), confirming the functional importance of YKT6 at the OMM.

To validate these spatial assignments, we selected two matrix mito-orphans, GTPBP8 and C17orf80, which exhibited multiple matrix-specific SR-PL sites (**Figure 4C**). Confocal microscopy confirmed that APEX2 fusion constructs for both proteins targeted the mitochondria (**Figure 4D**), and DAB-enhanced electron microscopy established that the C-termini of both proteins reside in the matrix (**Figure 4E**). GTPBP8 has recently been identified as a matrix GTPase required for mitoribosomal biogenesis,^42^ while C17orf80 was shown to interact with the mitochondrial nucleoid,^43^ providing independent support for their matrix localization.

Among the newly identified mito-orphans, small integral membrane proteins (SMIMs) pose a particular challenge due to their extremely short domains. Using MitoAtlas, we resolved the membrane topologies of seven SMIMs: SMIM4, SMIM8, SMIM12, SMIM20, and SMIM26 as IMM-TM; SMIM19 as MAM-TM; and MTLN (SMIM37) as OMM-TM. MitoCarta3.0^35^ previously categorized SMIM8 as “MOM” (OMM), while SMIM12, SMIM19, and SMIM26 were absent entirely. For cases where site-level data showed ambiguous IMS/OMM patterns — as expected for small IMM-TM proteins within the ∼10 nm reach of OMM-anchored BioID baits — AlphaFold-Multimer predictions with known partners (SMIM8–AGK, SMIM12–PHB1) confirmed the IMM-TM assignment (**Figures 4F** **and S4A**). Confocal microscopy and proteinase K protection assays further validated the mitochondrial localization and IMM-TM topology of SMIM8 and SMIM12 (**Figures 4G and 4H**).

Since the IMS hosts a unique disulfide bond relay system mediated by MIA40/CHCHD4,^44^ we validated compartment-specific modifications in newly annotated IMS proteins. MINPP1 and FKBP9 were each identified with a single IMS-classified tyrosine site (**Figure 4I**), and AlphaFold2 predicted disulfide bonds in both^45^ (**Figure 4J**). We focused on MINPP1, which contains a putative C-terminal ER-retention signal (SDEL) yet showed dual localization to both the ER lumen and the IMS by confocal microscopy (**Figure 4K**). Proteinase K protection confirmed its IMS localization (**Figure 4L**), and mmPEG24-maleimide modification confirmed the presence of disulfide bonds (**Figure 4M**). Unlike the bacterial homolog BtMinpp, human MINPP1 contains four predicted disulfide bonds with a highly conserved catalytic site (**Figures S4B–S4D**). Given its inositol polyphosphate phosphatase activity,^46^ we propose that IMS-localized MINPP1 at ER–mitochondria contact sites can hydrolyze InsP6 to generate IP3 and promote mitochondrial calcium flux (**Figure 4N**).

To assess the functional relevance of mito-orphans, we examined C. elegans orthologs with mitochondria-associated phenotypes upon RNAi knockdown (**Figure S4E**). We selected *ykt-6*, the ortholog of YKT6 identified here as a novel OMM protein, for functional characterization. RNAi-mediated knockdown of *ykt-6* resulted in a significant reduction in body-bending frequency (**Figures 4O and 4P**), indicating impaired organismal motility consistent with a role in mitochondrial physiology.

### Spatial Distribution of Functional Protein Domains Across Mitochondrial Sub-compartments

SR-PL data enables both membrane topology determination and spatial mapping of functional protein domains and homologous superfamilies across mitochondrial compartments (**Figure 5A**). The mitochondrial matrix is enriched for GTP-binding domains, biotin/lipoyl attachment motifs, LYR domains,1,3 and the NAD(P)-binding domain superfamily; spatial profiling of all NAD(P)-binding members confirms their predominant matrix sequestration (**Figures 5B** **and S5A**), enabling refined topological modeling of the inner membrane protein NNT (**Figure S5B**). The IMS is characterized by AAA-type ATPases, CHCH domains, and Tim10-like superfamilies. Notably, the EF-hand domain pair superfamily also shows a strong IMS preference (**Figure 5E**), consistent with the role of calcium signaling in this compartment. SR-PL site mapping of the calcium-binding mitochondrial carriers SLC25A13 and SLC25A24 resolves the spatial orientation of their EF-hand domains relative to the IMS (**Figure 5F**). LETM1, which mediates mitochondrial Ca²⁺/H⁺ exchange, is a notable exception; its EF-hand domain faces the matrix rather than the IMS, alongside a LETM1-like ribosome-binding domain (**Figure 5G**). The OMM is predominantly associated with Bcl-2 homology (BH) domains, zinc finger motifs, and TPR domains. TPR-containing OMM proteins (FIS1, FKBP8, TOMM70, RMDN2, RMDN3) are structurally positioned with cytosolic exposure, likely facilitating organelle–cytosol interactions (**Figures 5H, S5D, and S5E**).

**Figure 5.**
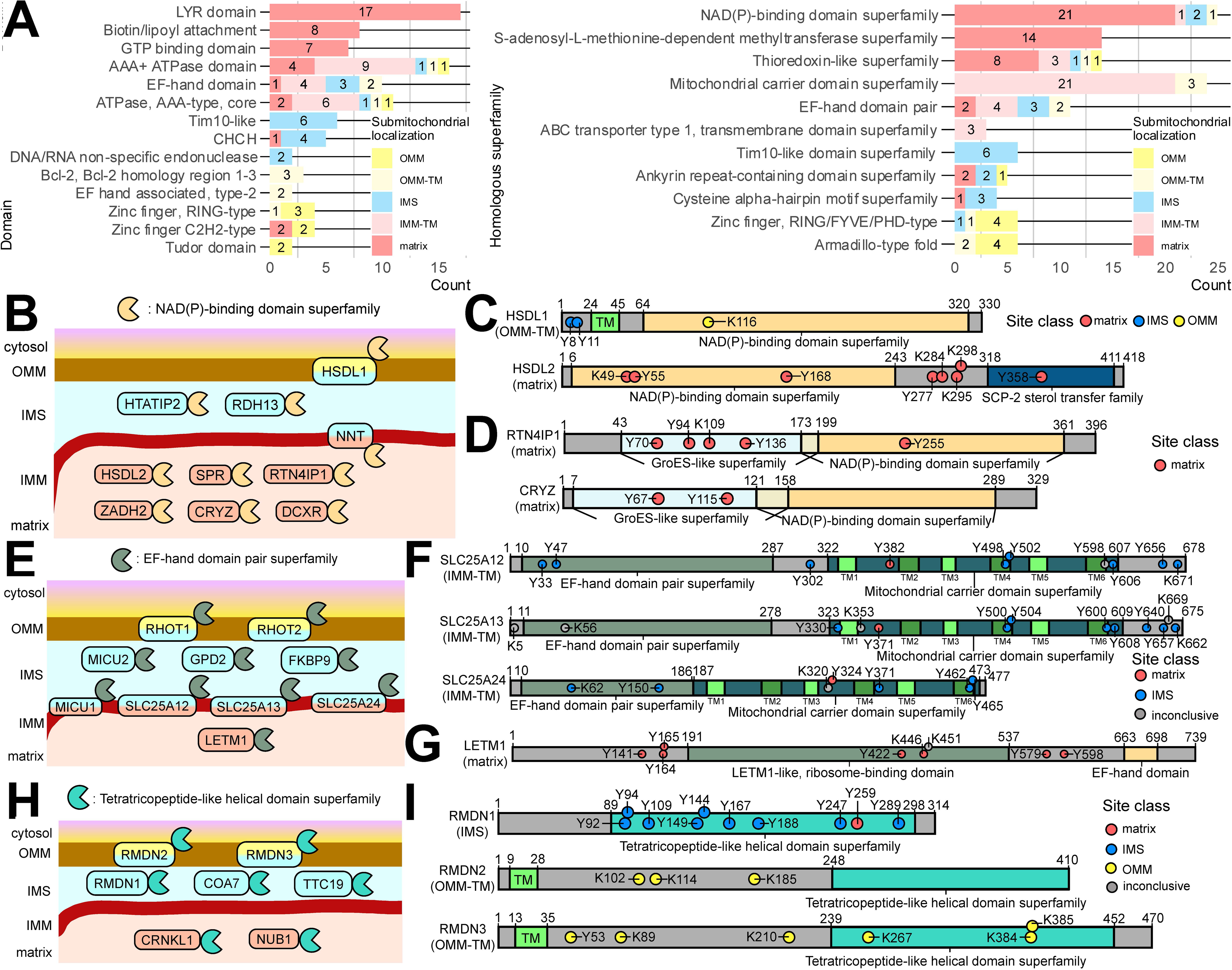
**Spatial Partitioning of Functional Domains and Homologous Superfamilies Across Mitochondrial Sub-compartments. (See also Figure S5 and Table S6)** (A) Comparative analysis of sub-mitochondrial enrichment for protein domains (left) and homologous superfamilies (right) based on InterPro^56^ classifications (B) Sub-mitochondrial distribution of proteins containing the NAD(P)H-binding domain superfamily, highlighting a predominant matrix localization. (C) Schematic representation of SR-PL sites and domain architectures for HSDL1 and HSDL2, illustrating the non-canonical OMM localization of HSDL1. (D) Schematic representation of SR-PL sites and domain architectures for RTN4IP1 and CRYZ. (E) Sub-mitochondrial distribution of the EF-hand domain pair superfamily, demonstrating a preference for the intermembrane space. (F) Schematic representation of SR-PL sites and domain architectures for the calcium-binding transporters SLC25A1, SLC25A13^65^, and SLC25A24^66^. (G) Schematic representation of SR-PL sites and domain architecture of LETM1. (H) Sub-mitochondrial distribution of the Tetratricopeptide-like (TPR) helical domain superfamily across mitochondrial compartments. (I) Labeled sites and domain architectures for RMDN1, RMDN2, and RMDN3, highlighting the atypical IMS localization of RMDN1 compared to its paralogs.

**Figure 6.**
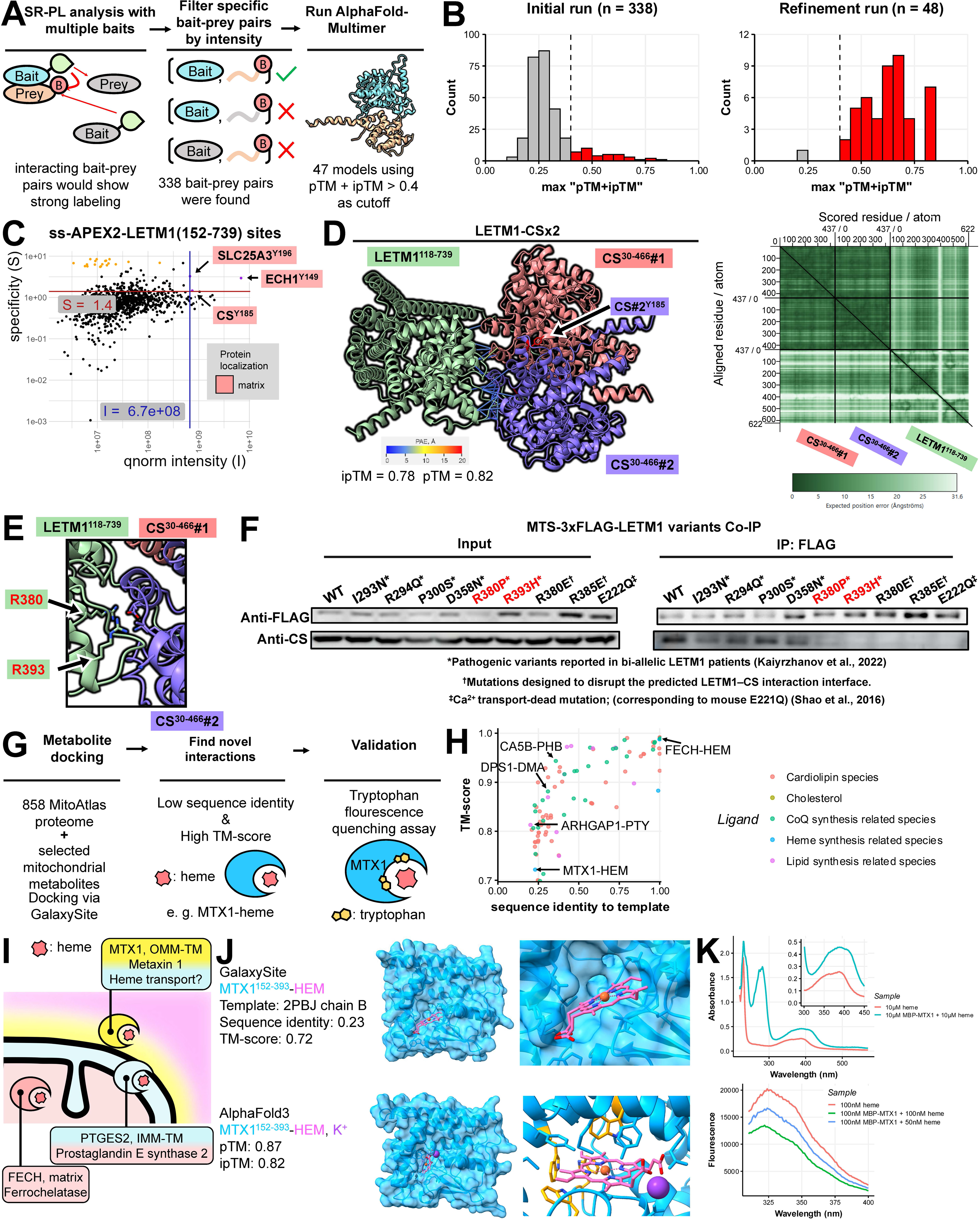
**Prediction of sub-mitochondrial protein-protein and protein-metabolite interactions (see also Figures S6 and S7 and Table S7).** (A) Workflow to find high pTM + ipTM scoring bait-prey models by filtering specific bait-prey sites. (B) Histogram of pTM + ipTM scores of AlphaFold models generated from bait-prey pairs with high specificity and intensity (left), and refined structures of score over 0.4 (right). (C) Quantile normalized intensity and specificity of ss-APEX2-LETM1 labeled sites. (D) AF3-Multimer prediction of trimer complex of one LETM1 and two CS with PAE values of residues near 4 angstroms between LETM1 and CS#1 or CS#2 highlighted (left). The LETM1 APEX2 specific site CS-Y185 is also depicted in red. PAE matrix is in the right. (E) Alternative view of AF3-Multimer prediction of LETM1-CSx2 trimer. Two residues at the LETM1-CS interfaces with known pathogenic consequences of mutation, R380 and R393, are highlighted. (F) Co-immunoprecipitation (CoIP) validation of the LETM1–CS interaction. MTS-3×FLAG-LETM1 (WT or indicated mutants) was transiently expressed in HEK293T-REx cells and immunoprecipitated with anti-FLAG beads; immunoblots were probed with anti-FLAG and anti-CS antibodies. Three classes of LETM1 variants were tested: (i) six pathogenic missense variants (I293N, R294Q, P300S, D358N, R380P, R393H) from bi-allelic patients with mitochondrial disease^7^; (ii) two structure-guided charge-reversal mutations (R380E, R385E) designed to disrupt the predicted LETM1–CS interaction interface; and (iii) E222Q, corresponding to mouse E221Q, which abolishes LETM1-mediated Ca2+/H+ antiport^50^. (G) Workflow used to find novel protein-metabolite interactions using MitoAtlas and GalaxySite. (H) Sequence identity and TM-score of predicted protein-metabolite interaction structures. (I) The sub-mitochondrial localizations of already known heme binding proteins FECH (matrix), PTGES2 (IMM-TM) and the newly discovered MTX1 (OMM). (J) Predicted complex structure of MTX1 and haem (HEM) modeled using GalaxySite docking with template restrictions (top) and AlphaFold3 (bottom). Tryptophan residues were colored orange in the AlphaFold3 model. (K) Absorbance spectrum of 10 μM MBP-MTX1 protein treated with heme compared to heme only (top) and Fluorescence spectrum of 100 nM MBP-MTX1 protein only, and 50 nM and 100 nM heme treated 100 nM MBP-MTX1 protein (bottom).

This high-resolution mapping also uncovers atypical spatial partitioning. HSDL1, a NAD(P)-binding protein, localizes to the OMM rather than the matrix (**Figures 5C** **and S5C**), and RMDN1 resides in the IMS unlike its OMM-localized TPR paralogs RMDN2 and RMDN3 (**Figure 5I**). Such unexpected localizations, combined with structural homology, enable functional inference: CRYZ shares a conserved fold with the matrix oxidoreductase RTN4IP1 (pruned RMSD 1.17 Å; **Figures 5D** **and S5C**), suggesting analogous activity in CoQ metabolism or redox homeostasis,^9,47,48^ and HSDL1 aligns tightly with HSDL2 (pruned RMSD 1.01 Å) despite their distinct compartments (**Figure S5C**). These examples demonstrate that MitoAtlas spatial data, combined with structural homology, can predict protein functions across sub-mitochondrial compartments.

### MitoAtlas-Guided Prediction of Sub-mitochondrial Protein–Protein and Protein–Metabolite Interactions

Proximity labeling captures not only protein localization but also the local protein neighborhood. By identifying proteins that share bait-specific labeled sites, MitoAtlas provides spatial constraints for predicting protein–protein interactions. We integrated these signatures with AlphaFold-Multimer to predict novel sub-mitochondrial interactions. Bait-prey pairs were filtered by site specificity and intensity and scored by AlphaFold-Multimer (**Figure 6A**); the resulting pTM + ipTM distribution identified a subset of high-confidence structural models (**Figure 6B**). For example, ss-APEX2-LETM1 yielded three specific labeled sites: CS-Y185, ECH1-Y149, and SLC25A3-Y196, all on proteins central to mitochondrial energy metabolism (**Figure 6C**). AF2-Multimer returned a high-scoring model for the LETM1–CS complex (0.8×ipTM + 0.2×pTM = 0.80), and AlphaFold3^49^ predicted a LETM1–CS×2 trimer with similar confidence (ipTM = 0.78, pTM = 0.82; **Figure 6D**). Two interface residues, R380 and R393, correspond to pathogenic variants (R380P, R393H) in LETM1-related mitochondrial disease (**Figure 6E**).^7^ Co-immunoprecipitation confirmed that these interface-proximal pathogenic variants, along with structure-guided charge-reversal mutations R380E and R385E, specifically abolished LETM1–CS binding, whereas pathogenic variants distal to the interface (I293N, R294Q, P300S, D358N) retained it (**Figure 6F**; **Figure S6A**). Notably, E222Q, which abolishes LETM1-mediated Ca^2+^/H^+^ antiport,^50^ also disrupted CS binding, suggesting that ion transport and CS interaction are structurally coupled. This interaction further supports LETM1’s reclassification as a peripheral matrix protein, since its binding partner CS is a matrix enzyme.

This approach generalizes beyond LETM1. For example, EXD2-APEX2 specific sites identified interactions with cytoskeletal proteins including CNN3 and ACTB (**Figures S6B and S6C**), consistent with EXD2’s OMM localization and cytosolic exposure. Network visualization of all EXD2 bait-prey connections (**Figure S6D**) and the complete bait-prey landscape across the 42-bait library (**Figure S6E**) illustrate the breadth of spatial interaction data captured by MitoAtlas.

MitoAtlas topology also independently validates external PPI predictions. Applying sub-mitochondrial topology as a spatial filter to 29,257 predicted mitochondrial PPIs,^51,52^ MitoAtlas identified 77 topologically incompatible pairs (7.0%), compared to 26 (1.7%) by MitoCarta3.0 (**Figure S6F**). Incompatible interactions exhibited significantly lower prediction confidence (median 0.831 vs. 0.946; Wilcoxon p = 6.27 × 10⁻⁷), providing reciprocal validation: MitoAtlas topology confirms computational predictions, while prediction scores validate MitoAtlas spatial assignments.

We next asked whether MitoAtlas could facilitate discovery of protein–metabolite interactions. Intersecting HProteome-BSite predictions^53^ with MitoAtlas-localized proteins, we screened for interactions involving mitochondria-specific metabolites (coenzyme Q, heme, cholesterol, lipids, cardiolipins), identifying 155 candidates with high structural similarity to known binding templates (TM-score ≥ 0.5; **Figure 6G, H**). Among these, GalaxySite docking and AlphaFold3 independently predicted a heme-binding site on MTX1 at the OMM (**Figure 6J**).^53–55^ Purified recombinant MBP–MTX1 exhibited tryptophan fluorescence quenching upon heme titration, confirming the interaction (**Figures S6G and 6K**).

### MitoAtlas.org: An Interactive Database for the Mitochondrial Research Community

MitoAtlas is accessible at mitoatlas.org (**Figure 7A**). For each of the 861 proteins, the platform provides a protein overview panel summarizing gene and protein names, synonyms, UniProt accession, database cross-references (MitoAtlas, UniProt/PubMed, MitoCarta), SR-PL labeled sites identified by APEX2 and BioID/BioID2, sub-mitochondrial localizations of both the protein and its individual sites, and transmembrane domain annotations. Five additional interactive modules enable in-depth exploration: a linear domain architecture map overlaying SR-PL labeled tyrosine and lysine sites onto InterPro domain annotations^56^ with compartment-specific color coding; an interactive membrane topology viewer that displays labeled sites across the lipid bilayer with logistic regression–derived topology scores; a multi-track sequence feature viewer integrating domain boundaries, labeled sites, AlphaFold2 pLDDT confidence scores, and biochemical properties in a single zoomable browser; a bait-resolved SR-PL intensity heatmap showing APEX2 and BioID signal profiles across all 42 proximity labeling baits with classification confidence for each site; and projection of labeled sites onto PDB and AlphaFold2 three-dimensional structures via the iCn3D viewer^57^, enabling spatial visualization of compartment assignments on experimentally determined and predicted protein structures. All data are downloadable in tabular format for computational analysis and integration with external datasets (tabular download function not implemented in current pre-submission stage).

**Figure 7.**
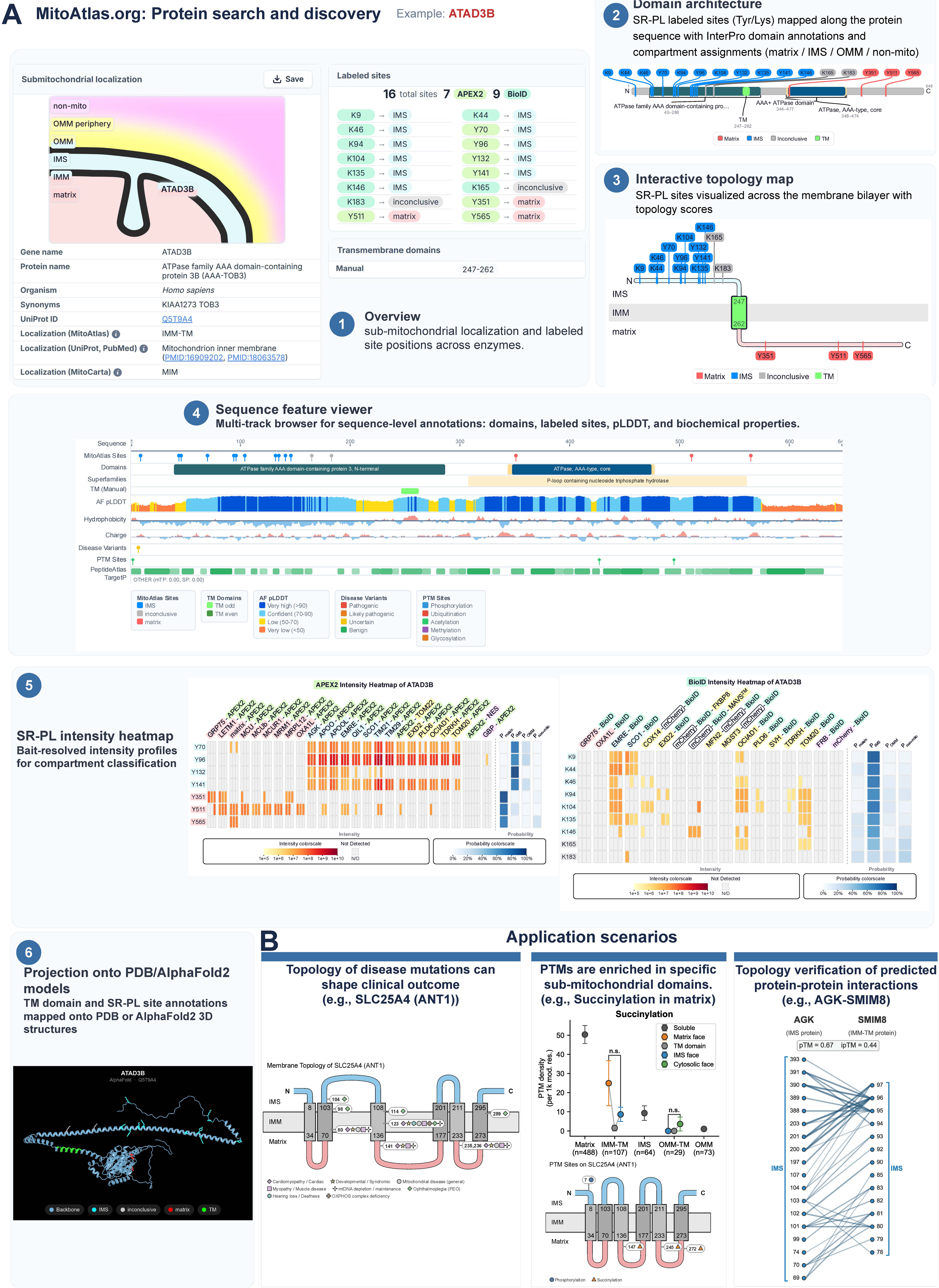
**MitoAtlas.org: an interactive web platform for the mitochondrial spatial proteome (see also Figure S7).** (A) Overview of MitoAtlas.org interface components, shown using ATAD3B as a representative example. The platform comprises six integrated modules: (i) Protein overview panel, displaying gene name, protein name, synonyms, UniProt ID, database cross-references (MitoAtlas, UniProt/PubMed sources, MitoCarta), SR-PL labeled sites identified by APEX2 and BioID, sub-mitochondrial localizations of the protein and individual sites, and transmembrane domain annotations; (ii) Linear domain architecture map, showing SR-PL labeled tyrosine and lysine sites along the full protein sequence with InterPro domain annotations^56^ and compartment assignments (matrix, IMS, OMM, or non-mitochondrial); (iii) Interactive membrane topology viewer, displaying SR-PL labeled sites across the membrane bilayer with color-coded compartment assignments and topology scores derived from logistic regression classification; (iv) Sequence feature viewer, a multi-track browser integrating domain boundaries, labeled sites, pLDDT confidence scores, and biochemical properties in a single linear view; (v) SR-PL intensity heatmap, showing bait-resolved APEX2 and BioID intensity profiles across all proximity labeling baits with logistic regression classification confidence for each site; and (vi) Projection onto PDB/AlphaFold2 3D structures, where TM domain and SR-PL site annotations are mapped onto experimentally determined or predicted structures via the interactive iCn3D viewer^57^. (B) Application scenarios demonstrating the utility of MitoAtlas for biological discovery (see Figure S7 for detailed examples). Three representative use cases illustrate how site-resolved spatial data enhances mitochondrial protein analysis: (i) Disease variant interpretation: membrane topology of SLC25A4 (ANT1) with ClinVar pathogenic^67^ variant mapping. Nine pathogenic variant positions are mapped onto the six-TM-helix topology and color/shape-coded by disease category, revealing a topology-dependent disease pattern: IMS-facing variants associate with progressive external ophthalmoplegia (adPEO), whereas matrix-facing variants cause cardiomyopathic mtDNA depletion syndrome. (ii) Post-translational modification mapping: PTM topology of SLC25A4 (ANT1) showing compartment-specific PTM distribution. A single phosphoserine (pos 7) localizes to the IMS-facing N-terminal segment, while three succinyllysine sites (pos 147, 245, 272) are exclusively on matrix-facing loops, consistent with IMS kinase accessibility and matrix succinyl-CoA–driven non-enzymatic acylation. (iii) Population variant density: gnomAD missense variant density^59^ across MitoAtlas-defined topological faces reveals strong purifying selection on TM domains of IMM proteins and reduced constraint on N-terminal mitochondrial targeting sequences.

Three application scenarios demonstrate how MitoAtlas site-resolved spatial data enhances biological discovery beyond compartment-level annotations (**Figure 7B**; see also **Figure S7**). First, mapping ClinVar^67^ pathogenic variants onto MitoAtlas membrane topology reveals topology-dependent disease patterns. For SLC25A4 (ANT1), an IMM carrier with six transmembrane helices, nine pathogenic variant positions segregate by sub-mitochondrial localization: variants within or near IMS-facing TM helices (positions 98, 104, 114, 289) are exclusively associated with autosomal dominant progressive external ophthalmoplegia (adPEO; MIM #609283), whereas variants in matrix-facing loops (positions 141, 235, 236) cause mtDNA depletion syndrome type 12 with cardiomyopathy (MIM #615418, #615438). Across all mitochondrial TM proteins, this approach assigned 3,139 disease variants in 88 genes to specific membrane faces: 556 (17.7%) to the matrix, 666 (21.2%) to the IMS, 502 (16.0%) within TM helices, and 1,415 (45.1%) to the cytosolic face of OMM proteins, a topological resolution unavailable from existing mitochondrial databases. Additional examples, including COX15, SLC25A22, and YME1L1, further illustrate topology-dependent disease segregation across diverse mitochondrial TM proteins (**Figure S7A**). Second, PTM sites show clear compartment specificity when mapped onto MitoAtlas topology. For ANT1, a single phosphoserine (position 7) localizes to the IMS-facing N-terminal segment, while three succinyllysine sites (positions 147, 245, 272) are exclusively found on matrix-facing loops, consistent with IMS kinase accessibility and non-enzymatic acylation driven by matrix succinyl-CoA concentration, respectively. Across all mitochondrial TM proteins, succinylation density is highest on the matrix face of IMM proteins and nearly absent from OMM-TM faces (**Figure 7B**). This compartment-dependent pattern extends to phosphorylation, acetylation, and ubiquitination across all mitochondrial TM proteins (**Figure S7B**). Third, analysis of gnomAD population variant density^59^ across MitoAtlas-defined topological faces reveals that TM domains of IMM proteins are strongly depleted in missense variants compared to soluble loops (43.1 vs. 45.0–51.8 variants per 100 aa; binomial p=8.0×10⁻ ³³), indicating strong purifying selection on membrane-embedded residues. N-terminal regions (positions 1–50), encompassing the mitochondrial targeting sequence, show consistently higher variant density than the remainder of the protein, consistent with reduced constraint on the cleaved MTS (**Figure S7C**).

In total, MitoAtlas contains 861 proteins, 13,440 SR-PL sites, 196 resolved topologies, 160 mito-orphans, 47 bait–prey interactions, and 155 protein–metabolite interactions, and will be updated as data from additional cell types become available.

## DISCUSSION

In this study, we present MitoAtlas, a spatial systems biology resource that resolves the sub-mitochondrial domain architecture of 861 proteins at high resolution. Rather than analyzing individual bait–prey relationships in isolation, MitoAtlas integrates proximity labeling data from 42 spatially defined APEX2 and BioID/BioID2 baits within a unified logistic regression framework, enabling systems-level classification that accounts for the full spatial context of each labeled site. This integrated approach identified 13,440 unique labeled sites whose MS intensity profiles encode precise spatial information. The logistic regression classifiers decode these profiles to assign sub-mitochondrial localizations at the level of individual tyrosine and lysine residues, a resolution not achievable by conventional fractionation-based approaches or binary bait-control comparisons.^2,12,16,35,36^ This site-level resolution enables MitoAtlas to provide not only protein localization but also membrane topology and domain architecture for all 196 transmembrane proteins in our dataset, information that is absent from existing mitochondrial databases. Moreover, MitoAtlas assigns spatial localizations to 160 proteins not previously annotated by MitoCarta3.0, substantially expanding the known mitochondrial proteome.

A key advance of MitoAtlas is the systematic determination of membrane topologies for mitochondrial transmembrane proteins. As of January 2026, UniProt contains curated topological annotations for only 52 IMM and 10 OMM proteins supported by specific experimental references.^29^ These annotations derive largely from protease-protection assays conducted on purified mitochondrial fractions under non-physiological conditions, which can yield conflicting results across laboratories. SR-PL overcomes these limitations by conducting proximity labeling in situ within intact living cells, and we demonstrate 97% concordance between our site assignments and AlphaFold2-predicted structures across multi-pass transmembrane carriers. Importantly, this concordance establishes MitoAtlas as an experimental validation platform for AI-predicted protein structures, a capability of growing significance as the field increasingly relies on computational structural models. MitoAtlas thereby provides a comprehensive and experimentally grounded topology resource that complements and extends the existing UniProt annotations.

Our topology analyses revealed several unexpected structural features. DNAJC11, previously classified as a peripheral OMM protein^60^, was reclassified as an OMM-TM protein with a predicted 17-stranded β-barrel architecture. This finding aligns with recent CRISPR screening data implicating DNAJC11 in the mitochondrial protein import pathway.^61^ While the original study attributed this phenotype to MICOS complex dysfunction^62^, the predicted pore diameter of DNAJC11 (14 Å) is comparable to that of TOMM40, the primary OMM import channel, raising the possibility that DNAJC11 may serve as an accessory import pore (**Figures S3E**). Similarly, our reclassification of LETM1 as a peripheral membrane protein rather than a single-pass IMM-TM protein informs its disputed role in Wolf–Hirschhorn Syndrome and its proposed function as a K+/H+ antiporter.^7^Notably, because K^+^/H^+^ antiport requires a transmembrane ion-conduction pathway, the reclassification of LETM1 as a peripheral protein raises the possibility that it participates in ion transport indirectly, perhaps as a regulatory subunit of a larger channel complex, rather than functioning as an autonomous antiporter. These examples illustrate how site-resolved proximity labeling can resolve longstanding topological ambiguities that have hindered functional characterization.

It is also noteworthy that our SR-PL findings uniquely demonstrate that BioID retains the BirA enzyme’s substrate preference^23^ (e.g., branched amino acids (Val, Leu, Ile) at the +1 position, **Figure S1F**), supporting its contact-dependent labeling mechanism.^20,25^ We hope that these insights will advance both the mechanistic understanding and rational engineering of proximity labeling enzymes.

Beyond topology determination, MitoAtlas enables systematic analysis of how sub-mitochondrial compartmentalization shapes both post-translational modification landscapes and evolutionary constraint. Mapping UniProt-annotated PTM sites onto MitoAtlas-defined topological faces reveals striking compartment specificity: phosphorylation is enriched on IMS-facing and cytosol-facing surfaces accessible to kinases, acetylation clusters on matrix-facing loops consistent with acetyl-CoA availability and SIRT3-regulated deacetylation, and ubiquitination is almost exclusively confined to the cytosol-facing domains of OMM proteins accessible to the ubiquitin-proteasome system (**Figure S7B**). These patterns demonstrate that the topology assignments are not merely structural descriptors but reflect the distinct biochemical environments of each mitochondrial compartment. Similarly, analysis of gnomAD population variant density^59^ across topological faces reveals that transmembrane domains of IMM proteins are under significantly stronger purifying selection than soluble loops (binomial p=2.0×10⁻ ³³), while N-terminal mitochondrial targeting sequences show elevated variant density consistent with reduced constraint on the cleaved MTS (**Figure S7C**). Together, these orthogonal validations, spanning disease genetics, post-translational biochemistry, and population genomics, demonstrate that site-resolved spatial data provides biological insights inaccessible from compartment-level annotations alone.

### Limitations of the Study

While MitoAtlas provides a comprehensive sub-mitochondrial spatial map, several limitations should be noted. First, all SR-PL data were generated in a single cell type (HEK293T-REx), which limits coverage of cell-type-specific proteins such as UCP1 in brown adipocytes or CYP11A1 in steroidogenic tissues; consequently, the 861 proteins resolved here are fewer than the collective inventories of databases that aggregate data across multiple cell types and organisms. However, this single-cell-type design provides an internally consistent spatial map in which all localizations, topologies, and expression levels are measured under identical conditions; information that cannot be extracted from aggregated databases. Second, the computationally predicted protein–protein and protein–metabolite interactions, although constrained by experimentally determined topologies, require systematic experimental validation. Third, proximity labeling coverage depends on bait accessibility, and proteins in poorly accessible sub-compartments may be underrepresented. We address this by providing probability-based confidence scores for all classifications, and the ∼1,700 inconclusive sites that did not meet the multi-bait consensus threshold are retained in the database for future investigation. To overcome these limitations, we are expanding MitoAtlas to include SR-PL datasets from additional human cell lines and mammalian tissues using transgenic mouse lines expressing mitochondria-targeted APEX2^9^, with the goal of building a tissue-resolved mitochondrial atlas.

## ACKNOWLEDGMENTS

MitoAtlas project was co-initiated by the laboratories of Prof. Hyun-Woo Rhee and Prof. Jong-Seo Kim. This work was supported by the National Research Foundation of Korea (RS-2025-15782968, RS-2023-KH136879, RS-2023-00260454 and RS-2023-00265581 to H.W.R.; RS-2024-00343424 and RS-2024-00440824 (the Bio&Medical Technology Development Program) to J.-S.K.). H.W.R. was supported by Samsung Science and Technology Foundation (SSTF-BA2201-08). J.-S.K. was supported by the Institute for Basic Science of the Ministry of Science and ICT of Korea (IBS-R008-D1). J.Y.M. and M.J. were supported by the Korea Brain Research Institute (KBRI) Basic Research Program, funded by the Ministry of Science and ICT (Information & Communication Technology) (23-BR-01-03). The EM data was acquired from the Brain Research Core Facilities of KBRI. Yun-Bin Lee helped cloning the novel mitochondrial protein APEX2-conjugated plasmids.

## AUTHOR CONTRIBUTIONS

H.W.R. and J.S.K., conceptualization; H.W.R., J.S.K., S.S., J.K., writing draft; S.Y.L. and C.K., cloning, stable cell generation and proteomics sample preparation; S.S., sample preparation, LC-MS analysis, and data analysis; J.K., data analysis, AF2 modeling, data curation and web application development; M.G.K., confocal microscope imaging; M.J. and J.Y.M., electron microscope imaging; H.L. and S.J.V.L., functional assay using *C. elegans*; J.S., metabolite docking; H.W.R. and J.S.K., supervision.

## DECLARATION OF GENERATIVE AI AN AI-ASSISTED TECHNOLOGIES IN THE WRITING PROCESS

During the preparation of this work the author(s) used Claude (Anthropic) in order to language editing and improving manuscript readability. After using this tool/service, the author(s) reviewed and edited the content as needed and take(s) full responsibility for the content of the publication.

## STAR★METHODS

### RESOURCE AVAILABILITY

#### Lead contact

Further information and requests for resources and reagents should be directed to and will be fulfilled by the lead contact, Hyun-Woo Rhee.

## Materials availability

All unique reagents generated in this study are available from the lead contact with a completed Materials Transfer Agreement.

## EXPERIMENTAL MODEL AND STUDY PARTICIPANT DETAILS

### Material and Method

#### Stable Cell Line Selection and Culturing

Flp-In T-REx 293 cells (Life Technologies) were transfected using Lipofectamine 2000 (Life Technologies), typically with 20 μL Lipofectamine 2000 and 4000 ng plasmid (9:1,pOG44:pCDNA5) per T25 flask. After 24 h, cells were cultured and selected with hygromycin B (100–200 μg/mL) for 2–3 weeks. To induce stable protein expression, the cells were treated with 100 ng/mL doxycycline. Cells were desthiobiotin-phenol-labeled or biotin-labeled depending on the bait enzyme used and were lysed 18–24 h after induction by doxycycline.

### Desthiobiotin-Phenol Labeling in Stably Expressed APEX Cell Lines

Stable-expression cells were incubated with 250 μM desthiobiotin-phenol at 37 °C under 5% CO2 for 30 min. Subsequently, 750 μL of 10 mM H_2_O_2_ (diluted from 30% H_2_O_2_, Sigma-Aldrich, H1009) was added to each flask, for a final concentration of 1 mM H_2_O_2_. The flasks were gently agitated for 1 min at room temperature. The reaction was then quenched by adding 7.5 mL of Dulbecco’s phosphate-buffered saline (DPBS) additionally containing 10 mM Trolox, 20 mM sodium azide, and 20 mM sodium ascorbate to each flask. The cells were then washed three times with cold quenching solution (DPBS containing 5 mM Trolox, 10 mM sodium azide, and 10 mM sodium ascorbate) and lysed with 1.5 mL RIPA lysis buffer (50 mM Tris, 150 mM NaCl, 0.1% SDS, 0.5% sodium deoxycholate, and 1% Triton X-100), 1× protease cocktail, 1 mM phenylmethylsulfonyl fluoride (PMSF), 10 mM sodium azide, 10 mM sodium ascorbate, and 5 mM Trolox for 10 min at 4 °C. Lysates were clarified by centrifugation at 15 000 ×*g* for 10 min at 4 °C.

### Biotin Labeling in Stably Expressed BioID and BioID2 Cell Lines

Stable-expression cell lines of promiscuous biotin ligases (BioID, BioID2) were incubated with 50 μM biotin (Alfa Aesar, A14207) at 37 °C for 16-18 hrs under 5% CO2 and lysed with RIPA buffer (ELPISBIO, EBA-1149) containing 1× protease cocktail (Invitrogen, 78438) for 30 min at 4°C. The cells were then washed three times with cold quenching solution (DPBS containing 5 mM Trolox, 10 mM sodium azide, and 10 mM sodium ascorbate) and lysed with 1.5 mL RIPA lysis buffer (50 mM Tris, 150 mM NaCl, 0.1% SDS, 0.5% sodium deoxycholate, and 1% Triton X-100), 1× protease cocktail, 1 mM phenylmethylsulfonyl fluoride (PMSF), 10 mM sodium azide, 10 mM sodium ascorbate, and 5 mM Trolox for 10 min at 4 °C. Lysates were clarified by centrifugation at 15 000 ×*g* for 10 min at 4 °C.

### Labeled Peptide Enrichment

For removal of unreacted free desthiobiotin-phenol or biotin-phenol, cell lysates were moved into an Amicon filter (Merck Millipore, 10 kDa cut-off) followed by centrifugation at 7500 ×g for 15 min at 4 °C. The volume was increased to 4 mL by adding phosphate-buffered saline (PBS) containing 1 mM PMSF and 1× protease cocktail, followed by centrifugation three more times. Finally, the cell lysates were transferred to an Eppendorf tube and mixed with 300 μL of streptavidin beads (Pierce). The sample was rotated for 1 h at room temperature and washed twice with PBS. After removing the PBS, 100 μL of denaturing solution (6 M urea, 2 M thiourea, and 10 mM 4-(2-hydroxyethyl)-1-piperazineethanesulfonic acid (HEPES)) was added and reduced using 20 μL of 100 mM dithiothreitol (DTT) in 50 mM ammonium bicarbonate (ABC) buffer for 60 min at 56 °C using a ThermoMixer (Eppendorf). Protein alkylation was performed by adding 35 μL of 300 mM iodoacetamide in 50 mM ABC buffer with shaking in the dark for 30 min. Trypsin gold (Promega) was then added to the solution, which was then incubated at 37 °C with shaking overnight. Formic acid was then added to terminate the trypsin reaction, and the beads were washed with PBS four times. After adding 250 μL of 95% formamide, and 10 mM ethylene-diamine-tetraacetic acid (EDTA) (pH 8.2), the elute was obtained by boiling at 95 °C for 10 min. The eluted peptide samples were then desalted using Varian Bond ELUT (Agilent, 12109301) and a home-made column.

### Liquid chromatography tandem mass spectrometry (LC-MS/MS) analysis of enriched peptide samples

Analytical capillary columns (100 cm × 75 µm i.d.) and trap columns (2 cm × 150 µm i.d.) were packed in-house with 3 µm Jupiter C18 particles (Phenomenex, Torrance). The long analytical column was placed in a column heater (Analytical Sales and Services) regulated to a temperature of 45°C. The nanoAcquity UPLC system (Waters, Milford) was operated at a flow rate of 300 nL/min over 2 h with a linear gradient ranging from 95% solvent A (cH2O with 0.1% formic acid) to 40% of solvent B (acetonitrile with 0.1% formic acid). The enriched samples were analyzed on an Orbitrap Fusion Lumos mass spectrometer (Thermo Fisher Scientific) equipped with an in-house customized nanoelectrospray ion source. Precursor ions were acquired at 120 k resolving power (m/z 300–1500), and the isolation of precursor ions for MS/MS analysis was performed with a 1.4 Th. Higher-energy collisional dissociation (HCD) with 30% collision energy was used for sequencing with auto gain control (AGC) target of 1 × 10^9^. The resolving power to acquire the MS2 spectra was set to 30 k with 200 ms maximum injection time.

### MS data processing and protein identification

All MS/MS data were screened using MaxQuant (v. 1.6.2.3) with the Andromeda search engine at a 10 ppm precursor ion mass tolerance against the SwissProt Homo sapiens proteome database (20,199 entries, UniProt; http://www.uniprot.org). Label free quantification (LFQ) and Match Between Runs were used, with the following search parameters for BioID experiments: semi-tryptic digestion, fixed carbamidomethylation of cysteine, dynamic oxidation of methionine, protein N-terminal acetylation, and biotinylation for lysine residues. Following parameters were used for APEX2 experiment: tryptic digestion, fixed carbamidomethylation on cysteine, dynamic oxidation of methionine, and protein N-terminal acetylation with DBP (+331.1896) for tyrosine residues. A false discovery rate (FDR) < 1% was obtained for the uniquely labeled peptides and unique labeled proteins. LFQ intensity values were log-transformed for further analysis.

### Sub-mitochondrial localization classification by logistic regression

Sub-mitochondrial localization was assigned through a two-step logistic regression (LR) pipeline implemented in Python (scikit-learn v1.2). In Step 1 (Site Classification, P1), separate one-vs-rest (OVR) LR models were trained for APEX2 and BioID datasets. The APEX2 site LR model was trained on 660 manually curated sites (Matrix: 135, IMS: 149, OMM: 107, Cytosol: 269) and applied to predict the localization of all 6,046 APEX2-labeled tyrosine sites. Input features consisted of log10-transformed MS intensities across all 25 APEX2 baits, calibrated by iBAQ protein abundance. The BioID site LR model was trained on 576 manually curated sites (Matrix: 85, IMS: 80, OMM: 216, Cytosol: 195) and applied to predict the localization of all 7,394 BioID-labeled lysine sites, using log10-transformed MS intensities across all 17 BioID baits as features. Training sites were curated from proteins with well-established sub-mitochondrial localizations in the literature, with transmembrane proteins manually annotated at the individual residue level based on known topology. For both models, 10-fold cross-validation accuracy exceeded 90%, and probability thresholds were optimized to maintain an overall false discovery rate (FDR) below 10%.

In Step 2 (Protein Classification, P2), site-level P1 outputs were aggregated into protein-level features. For each protein, the logit-transformed P1 probabilities were summed across all labeled sites, yielding an 8-dimensional feature vector (4 APEX2 compartment scores + 4 BioID compartment scores: Matrix, IMS, OMM, Non-mitochondrial). A second OVR LR model was trained on curated reference protein localizations and applied to classify the 3,825 non-TM proteins. The remaining 233 TM proteins were instead routed to a topology scoring pipeline that integrates site-level P1 classifications with transmembrane domain predictions from DeepTMHMM, TMbed, and AlphaFold2 structural models. In total, 4,058 proteins were classified across both arms of the pipeline. Non-TM proteins were assigned to tiered confidence categories: Gold (≥2 concordant sites above the P1 threshold) or Silver (1 site above the P1 threshold) for Matrix and IMS assignments. For OMM, the thresholds were set at ≥3 sites (Gold) and ≥2 sites (Silver) to account for the higher background labeling in the outer membrane compartment. Complete training sets and classification outputs are provided in Tables S1–S2.

### Comparison of binary enrichment analysis and MitoAtlas classification

To evaluate the added value of integrating multiple proximity labeling datasets over a conventional single-experiment analysis, we compared binary enrichment analysis with the MitoAtlas logistic regression (LR) classifier for discriminating outer mitochondrial membrane (OMM) from non-mitochondrial (cytosolic) biotinylation sites. For binary enrichment analysis, site-level intensities were extracted for the OMM-targeted bait condition and the cytosolic control condition (APEX2: TOM20-APEX2, n = 3, versus APEX2-NES, n = 6; BioID: TOM20-BioID, n = 3, versus mCherry-BioID, n = 3). Raw intensities were log2-transformed and median-centered across all channels to a common global median. Missing values were imputed using a Perseus-style missing-not-at-random (MNAR) strategy, in which each missing value was drawn from a down-shifted normal distribution (mean = column mean – 1.8 SD, width = 0.3 SD) ^63^. For each biotinylation site, the log2 fold change was calculated as the difference in mean log2 intensity between bait and control replicates, and statistical significance was assessed by a two-sided Welch’s t-test. For MitoAtlas classification, the pre-computed LR posterior probabilities across compartments were used directly. Receiver operating characteristic (ROC) analysis was performed on the subset of sites with known compartment assignments (OMM as positive class; non-mitochondrial as negative class). For the binary approach, log2 fold change served as the classification score; for MitoAtlas, p(OMM) was used. The classification threshold was swept across 500 evenly spaced values spanning the full range of each score, and the true positive rate and false positive rate were computed at each threshold to generate the ROC curve. The area under the ROC curve (AUC) was calculated by the trapezoidal rule. For APEX2, the analysis included 107 OMM and 269 non-mitochondrial training sites; for BioID, 216 OMM and 195 non-mitochondrial training sites. All analyses were performed in R (v4.5) using ggplot2 for visualization.

### Transmission Electron Microscopy

HEK293T cells were grown in 35-mm glass grid-bottomed culture dishes to 50%−60% confluence. The cells were transfected with APEX2-conjugated plasmid with lipofectamine reagent. The next day, the cells were fixed with 2.5% glutaraldehyde and 2% paraformaldehyde in 0.1 M cacodylate solution (pH 7.0). After washing, DAB staining was performed for 5−45 min until a light brown stain was visible under an inverted light microscope. DAB-stained cells were post-fixed with 2% potassium ferrocyanide-reduced osmium tetroxide for 1 h at 4 °C. The fixed cells were dehydrated using an ethanol series (50%, 60%, 70%, 80%, 90%, and 100%) for 10 min at each concentration, then infiltrated with an embedding medium. The identified samples were cut horizontally to the plane of the block into 60 nm sections (UC7; Leica Microsystems, Germany), mounted on copper slot grids with a specimen support film, then, double stained with UranyLess (Electron Microscopy Sciences, #22409) for 2 min and 3% lead citrate (Electron Microscopy Sciences, #22410) for 1 min. The sections were then analyzed using a transmission electron microscope at 120 kV (Tecnai G2, ThermoFisher, USA).

### AlphaFold2

AlphaFold v2.3.0 was downloaded from GitHub and installed as a dockerized application under Linux (Ubuntu 22, AMD Ryzen 9 5950X, 128 GB RAM, Nvidia RTX 4090 with 24GB RAM). The full databases were stored on a 4TB NVMe SSD using the flag "db_preset=full_dbs", and 25 multimer models were generated using the flag "num_multimer_predictions_per_model=5".

### Data Visualization

Protein structure visualization and PAE visualization were performed using UCSF ChimeraX (https://www.rbvi.ucsf.edu/chimerax).

### Web Application

The mitoatlas.org web application is based on the JavaScript frameworks React (https://react.dev/) and ExpressJS (https://expressjs.com/). The experimental data are stored and accessed using a mySQL relational database (https://www.mysql.com/). The iCn3D viewer was embedded using the npm package (https://www.npmjs.com/package/icn3d) to visualize the labeled sites on the AlphaFold2 structure. D3.js (https://d3js.org/) was used to generate heatmaps of the MS data. PubMed queries were performed using the Entrez Programming Utilities API (https://www.ncbi.nlm.nih.gov/books/NBK25500/).

### TM domain annotation sources

UniProt TM domain annotation were downloaded from the website using the UniProt ID-mapping service (https://www.uniprot.org/id-mapping). TMbed transmembrane domain annotations were downloaded from the pre-calculated human proteome data (https://rostlab.org/public/tmbed/predictions/human_210422_tmbed.tar.gz). DeepTMHMM TM domain annotations were obtained by running the BioLib application (https://dtu.biolib.com/DeepTMHMM/) on a local machine (Ubuntu 22, AMD Ryzen 9 5950X, 128 GB RAM, NVidia RTX 4090 with 24 GB RAM) as per the instructions on the website, using the UniProt primary protein sequence as input. AF2-TMDET TM domain boundaries were obtained by running TMDET 4.1 via Docker (brgenzim/tmdet:4.1.1) on AlphaFold2-predicted structures (CIF format) for each protein.^30^

### Membrane topology scoring and assignment

For each TM protein classified in Step 2, membrane topology was assigned using a topology scoring algorithm that integrates site-level sub-mitochondrial localizations (from Step 1) with TM domain boundary predictions from four independent sources: UniProt (curated annotations), DeepTMHMM (deep learning), TMbed (protein language model embeddings), and AF2-TMDET (AlphaFold2 structure-based detection). For each source, every SR-PL labeled site was classified relative to the predicted TM segments as residing in the N-terminal loop, within a TM segment, in an inter-TM loop, or in the C-terminal loop. Sites in loop regions contributed a full count to the corresponding membrane face (lower or upper), while sites within TM segments contributed fractional counts to both faces based on their relative position within the segment (logit-transformed). IMM and OMM hypotheses were then scored: the IMM score rewards matrix sites on one face and IMS sites on the other; the OMM score rewards IMS sites on one face and OMM or cytosolic sites on the other. The source with the highest adjusted score was selected as the adopted TM model, with adjustments including a structural confidence bonus for AF2-TMDET predictions, penalties for short TM segments and TM count outliers, and a boundary-agreement bonus when independent methods (AF2-TMDET and TMbed) converged on the same TM architecture. Ties were resolved by consensus TM count among tied sources, with manual review for unresolved cases. N- and C-terminus positions were then assigned based on the winning orientation (score1 vs. score2) and the determined membrane identity.

For proteins where the topology scoring algorithm could not confidently distinguish between IMM and OMM assignment, typically due to ambiguous site localizations (e.g., IMS-only evidence, which is consistent with both membranes), AlphaFold-Multimer structure predictions with known interacting partners were used as additional evidence to resolve the membrane assignment. For example, the predicted SMIM8–AGK and SMIM12–PHB1 complexes confirmed IMM-TM assignments, while the SMIM20–COX14–MTCO1, and HIGD2A–MTCO3 complexes provided independent structural evidence for IMM localization. In cases where the algorithm returned ambiguous results for proteins with well-established sub-mitochondrial localization (e.g., OMM-resident TOM complex subunits and VDACs), the known sub-mitochondrial context was used to assign membrane identity.

### Co-immunoprecipitation (CoIP)

HEK293T-REx cells were transiently transfected with MTS-3×FLAG-LETM1 (WT or mutant) constructs using polyethyleneimine (PEI) and harvested 24 h post-transfection. Cells were lysed in PBS (pH 7.4) containing 1% NP-40 and 1× protease inhibitor cocktail for 10 min at 4 °C. Lysates were cleared by centrifugation at 15,000 ×g for 5 min and incubated with anti-FLAG affinity gel (ChemScene) for 2 h at 4 °C with rotation. Beads were washed twice with PBS containing 0.1% NP-40 and eluted by boiling in 2× SDS-PAGE loading buffer at 95 °C for 5 min. Input and elution fractions were analyzed by western blot using anti-FLAG and anti-CS antibodies.

1. *C. elegans* strains and RNAi clones
2. *C. elegans* strain N2 (wild-type) was maintained under standard laboratory culture conditions on nematode growth medium (NGM) seeded with *E. coli* OP50 or HT115, which expressed double-stranded RNA for RNAi experiments. The RNAi clones used in this study were L4440 (control) and *ykt-6*.

Measurement of *C. elegans* swimming rates

The *C. elegans* swimming rate (body bends per min) was measured as described previously.^58^ Wild-type worms were treated with control RNAi or *ykt-6* RNAi for two generations before performing the assay. Fifteen pre-fertile adult worms were transferred to 24-well plates with 1 mL of M9 buffer per well. After stabilization for 1 min, body bends were recorded for 30 s using a digital microscope (DIMIS-M, Siwon Optical Technology, Anyang, South Korea); this number was then converted to the number of bends per min. *P* values were calculated using the unpaired Student’s *t*-test (two-tailed), comparing the swimming rates of the control and RNAi-treated worms.

### Measurement of C. elegans developmental rates

Images of control RNAi- and *ykt-6* RNAi-treated worms cultured for two generations were obtained using a digital microscope (DIMIS-M, Siwon Optical Technology, Anyang, South Korea). Adult *C. elegans* were washed with M9 buffer and the remaining eggs were incubated for 2 hrs at 20°C. Hatched L1 larvae were transferred to new NGM plates seeded with OP50. Images were obtained at 20 °C, 44 h after transfer.

### Ligand-Binding Site Similarity Search and Docking Analysis

A similarity-based search for mitochondrial protein targets that may interact with key metabolites like heme was done using HProteome-BSite^53^. HProteome-BSite is a database which contains ligand binding sites and putative ligands for human proteins. With protein annotations from MitoAtlas, there were 263 mitochondrial proteins with HEM as their putative binding partner, and 209 with PTY. Findings included novel OMM protein like ARHGAP1. Complex structure of PTY and ARHGAP1, one of the novel OMM protein, was modeled using GalaxyDock3^54^, and CSAlign^55^. Some novel interactions were found for known mitochondrial proteins such as MTX1. Results for MTX1 suggested interaction with HEM, and its complex structure was modeled using docking algorithm from latest version of GalaxySite^53^.

## SUPPLEMENTAL INFORMATION

Figure S1. Validation of APEX2 and BioID/BioID2 stably expressing cells, related to Figure 1. Figure S2. Site and Protein Classification Model Details, related to Figure 2.

Figure S3. Validation and Structural Refinement of Mitochondrial Membrane Topologies, related to Figure 3.

Figure S4. Structural Validation and Functional Analysis of Mito-Orphan Proteins, related to Figure 4.

Figure S5. Spatial Enrichment and Membrane Topology of NAD(P)-binding and Tetratricopeptide Helical Domain Superfamilies, related to Figure 5.

Figure S6. CoIP optimization, specific bait–prey site analysis, and topology validation of predicted interactome, related to Figure 6.

Figure S7. Application Scenarios: Enhanced Biological Discovery with MitoAtlas Site-Resolved Spatial Data, related to Figure 7.

## SUPPLEMENTAL VIDEOS AND SPREADSHEETS

Table S1. LC-MS/MS detection of biotin or DBP labeled SR-PL sites, related to Figure 1. Table S2. Classification of SR-PL sites into sub-mitochondrial spaces, related to Figure 2. Table S3. Classification of proteins into sub-mitochondrial spaces, related to Figures 2 and 3. Table S4. Membrane topology information from SR-PL, related to Figure 3.

Table S5. Further study on mito-orphans, related to Figure 4.

Table S6. Interpro domain annotations of MitoAtlas proteins, related to Figure 5.

Table S7. Protein-protein interactions and protein-metabolite interactions of MitoAtlas proteins, related to Figure 6.

**Figure S1.**
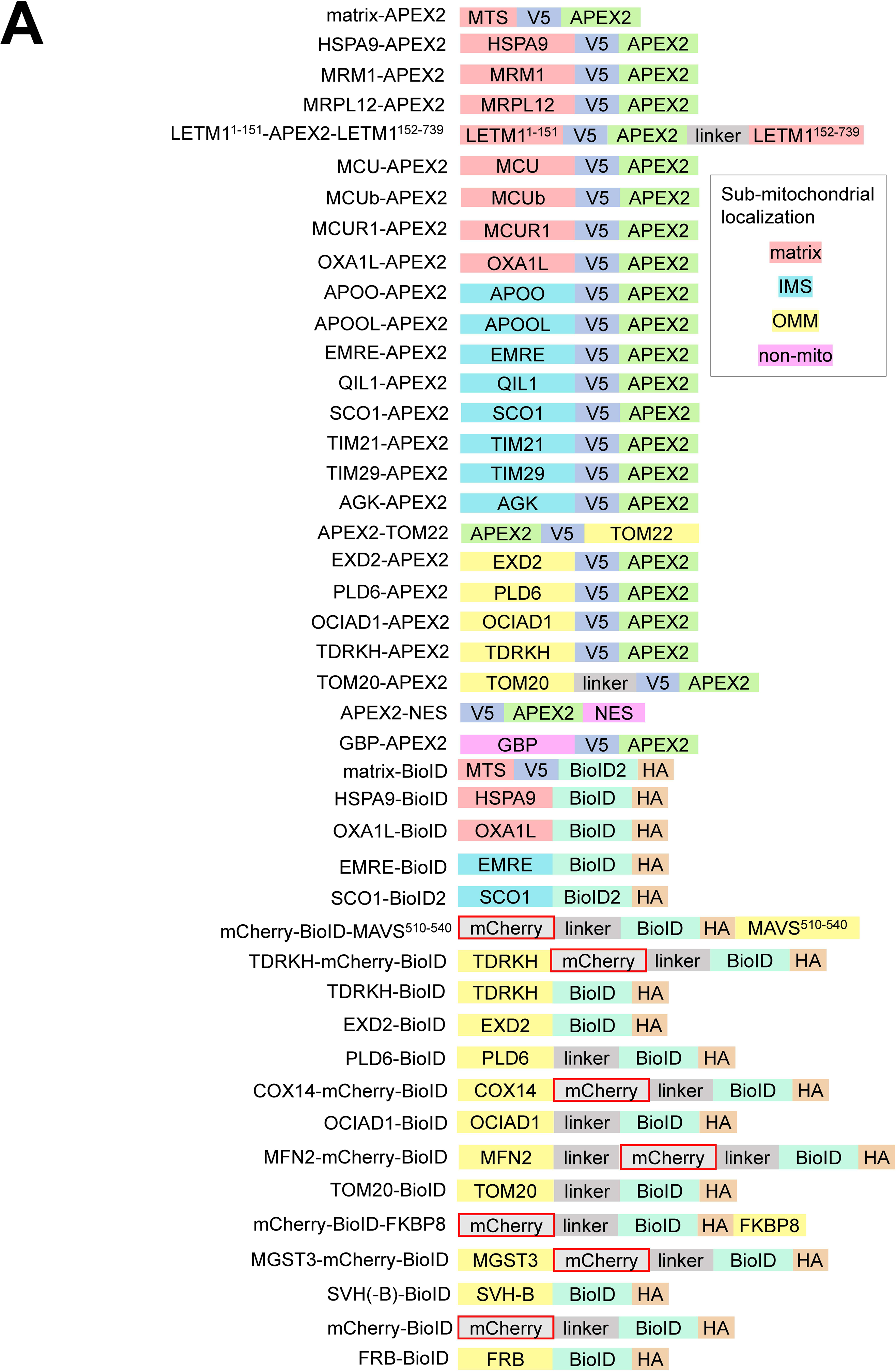
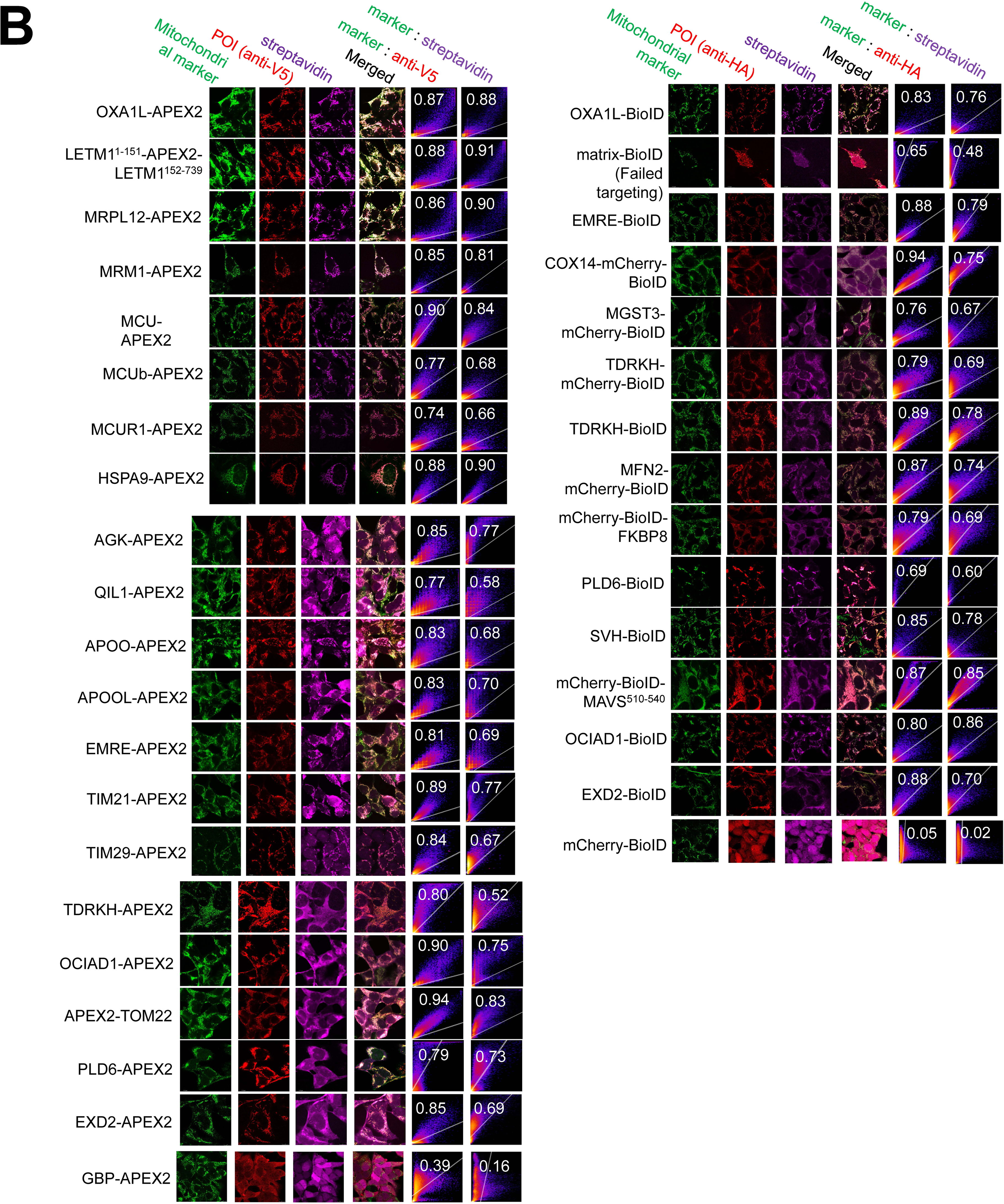
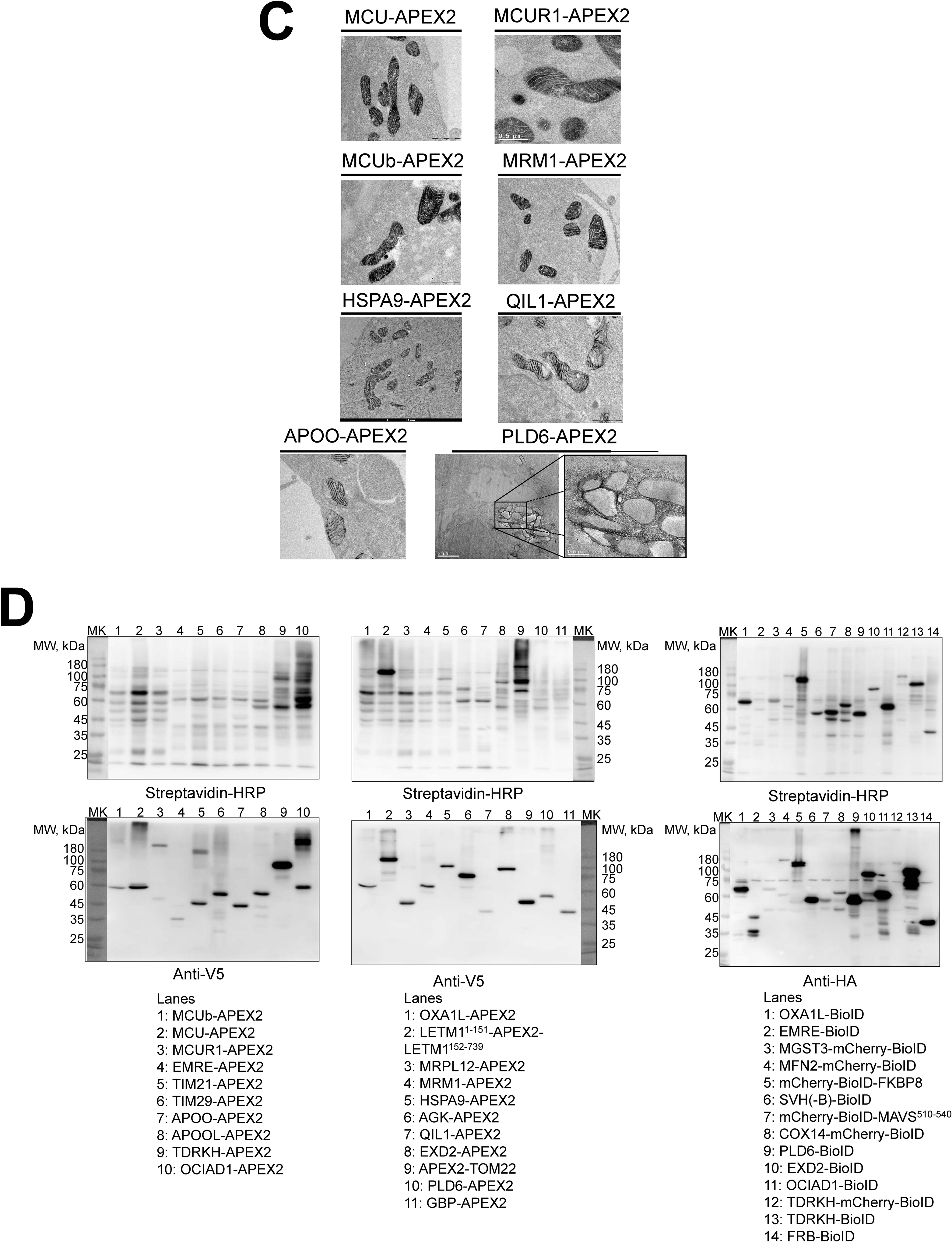
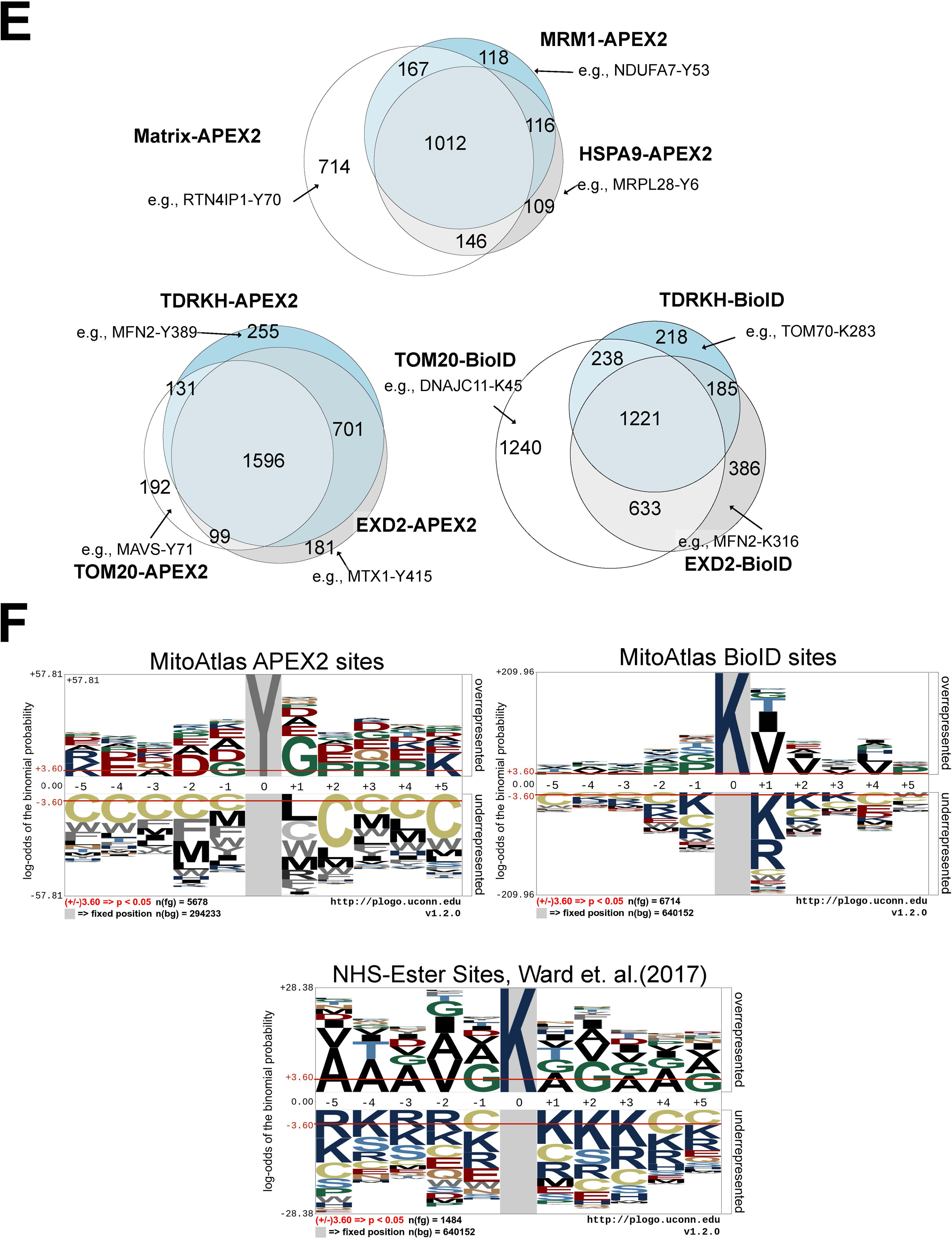
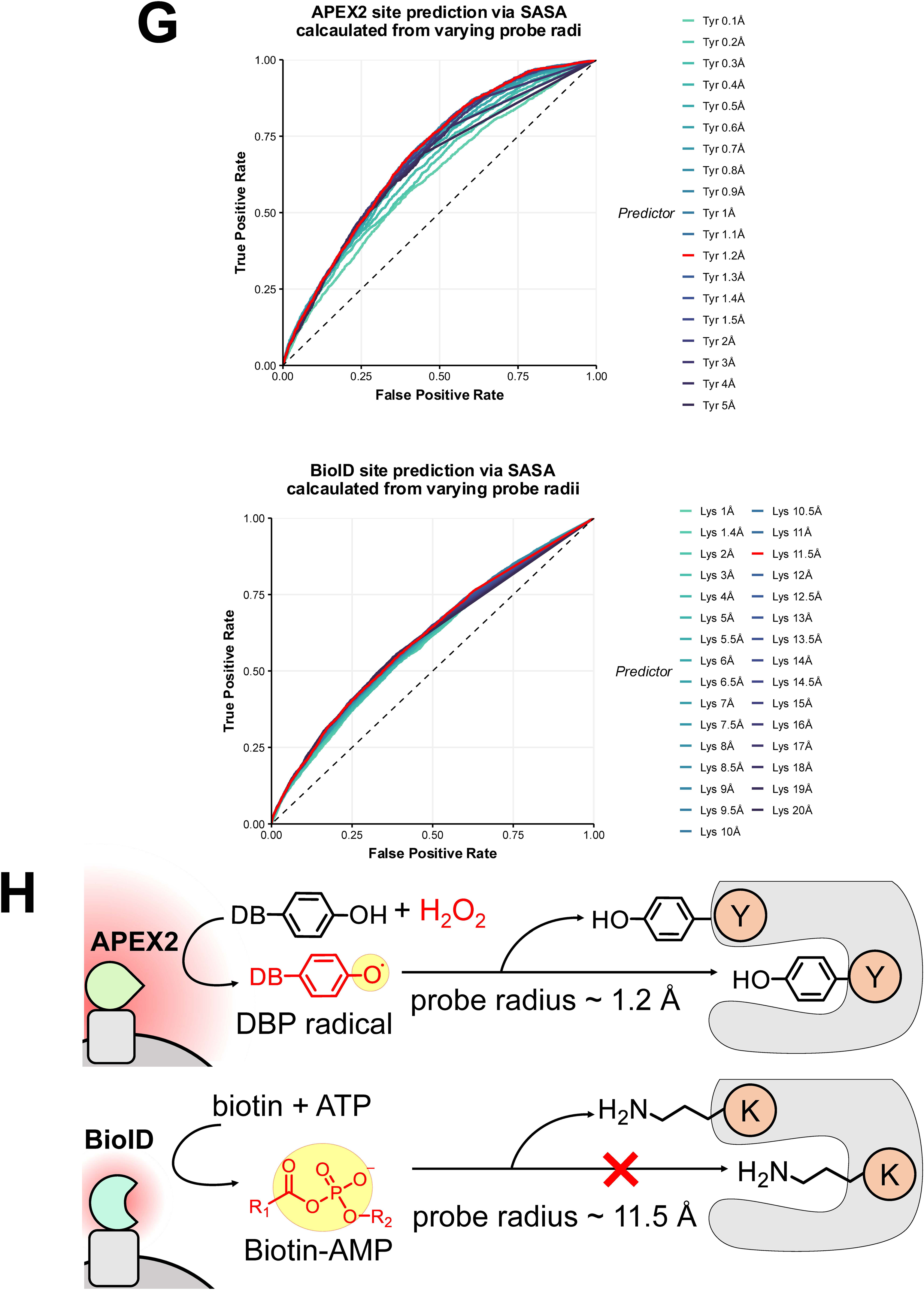
Validation of APEX2 and BioID/BioID2 stably expressing cells (related to. Figure 1). (A) Architecture of all APEX2- and BioID/BioID2-fused bait constructs used in this study. APEX2 baits are tagged with V5; BioID/BioID2 baits are tagged with HA. Inclusion of mCherry as a linker in specific constructs was utilized to enhance protein solubility or adjust bait size. Bait proteins are color-coded by their targeted sub-mitochondrial compartment: matrix (red), IMS (blue), OMM (yellow), and non-mitochondrial (purple). (B) Confocal immunofluorescence microscopy of all HEK293T-REx stable cell lines. APEX2 baits (left, anti-V5) and BioID/BioID2 baits (right, anti-HA) were co-stained with a mitochondrial marker and streptavidin to confirm sub-organellar targeting and biotinylation activity. Pearson’s correlation coefficients for colocalization with the mitochondrial marker are shown. Merged images display the overlay of the POI signal and the mitochondrial marker. (C) Transmission electron microscopy (TEM) images of APEX2-mediated 3,3’-diaminobenzidine (DAB) polymerization for representative matrix- and IMS-targeted baits, providing ultrastructural validation of bait positioning. (D) Western blot analysis using Streptavidin-HRP (top rows) and anti-V5 or anti-HA antibodies (bottom rows) to confirm expression levels and total protein biotinylation across the stable cell lines. (E) Venn diagrams illustrating the overlap and unique coverage of modified peptides identified by multiple baits within the same sub-compartment. Top: three matrix APEX2 baits (Matrix-APEX2, MRM1-APEX2, HSPA9-APEX2) show extensive non-overlapping peptide sets, demonstrating the value of multiplexed baits for expanding proteome depth. Bottom left: three OMM-targeted APEX2 baits (TDRKH-APEX2, TOM20-APEX2, EXD2-APEX2). Bottom right: three OMM-targeted BioID baits (TDRKH-BioID, TOM20-BioID, EXD2-BioID). Comparison of the bottom panels reveals that BioID generates a higher proportion of non-overlapping peptides relative to APEX2, consistent with its shorter labeling radius capturing more spatially restricted protein neighborhoods. (F) Sequence logos highlighting the local amino acid preferences surrounding APEX2-labeled tyrosine residues (left) and BioID/BioID2-labeled lysine residues (right), compared with previously reported N-hydroxysuccinimide (NHS)-ester-modified lysine sites (bottom). APEX2-modified sites show enrichment of glycine at ±1 positions, while BioID sites favor branched amino acids (Val, Leu, Ile) at the +1 position. Conversely, radical-quenching amino acids (Cys, Trp, Met, Arg, Phe, Leu, Tyr, His) are negatively enriched around APEX2 sites, with cysteine showing the strongest depletion, consistent with its favorable one-electron reduction potential for phenoxyl radical scavenging. (G) Receiver Operating Characteristic (ROC) curves evaluating the correlation between solvent-accessible surface area (SASA) and peptide detection using AlphaFold2 structures. Top: APEX2 site prediction at varying probe radii, showing an optimal probe radius near that of water (∼1.2 Å), suggesting APEX2 can probe deep into small protein clefts. Bottom: BioID site prediction, exhibiting a much larger optimal radius (∼11.5 Å), consistent with the bulkiness of the biotin-AMP molecule restricting labeling to more exposed surface residues. (H) Mechanistic model illustrating the differences in reactive probe size between APEX2 and BioID. The small desthiobiotin-phenol (DBP) radical generated by APEX2 (∼1.2 Å probe radius) can access sterically restricted tyrosine residues, whereas the larger biotin-AMP intermediate produced by BioID (∼11.5 Å probe radius) is limited to more solvent-exposed lysine residues.

**Figure S2.**
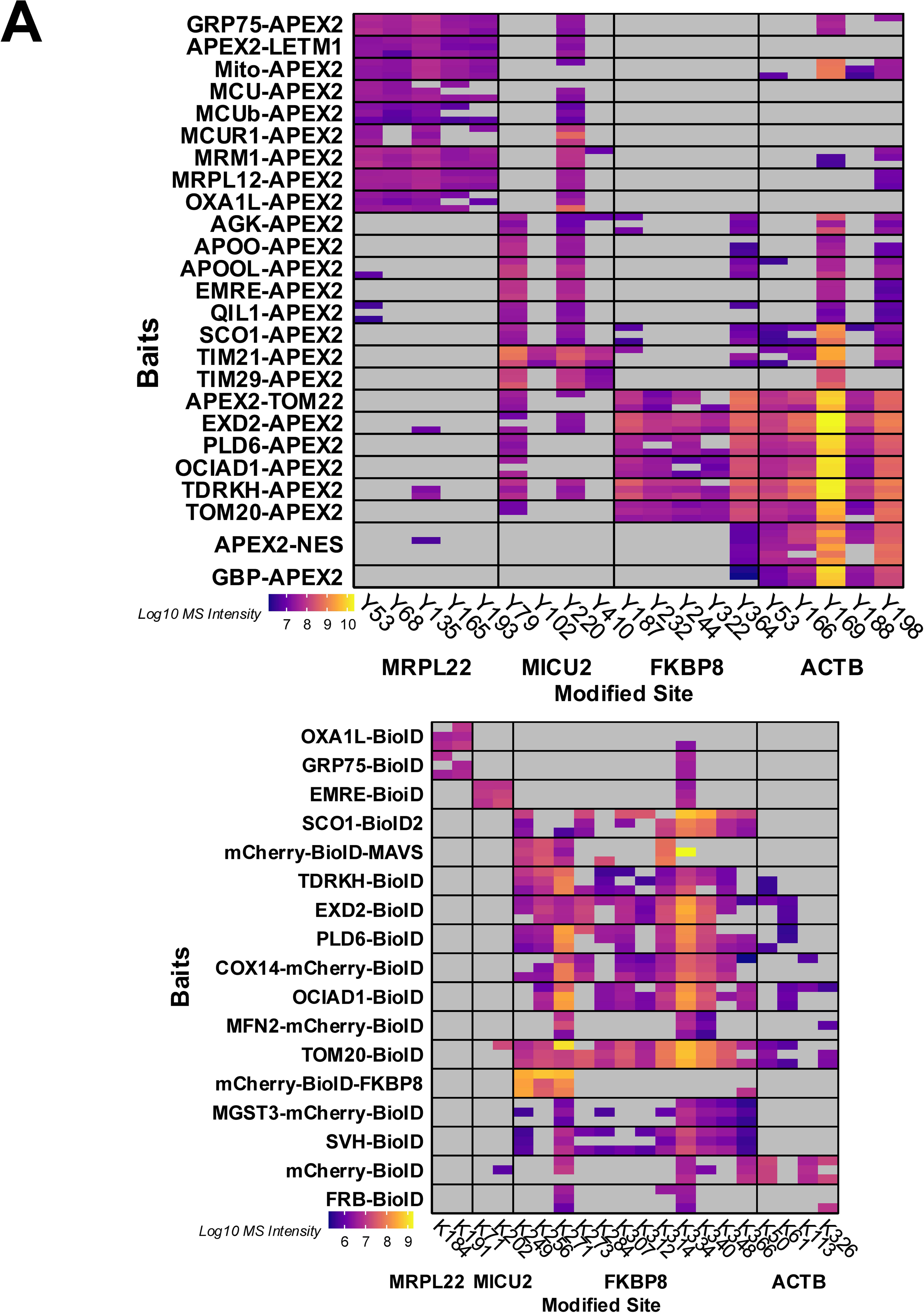
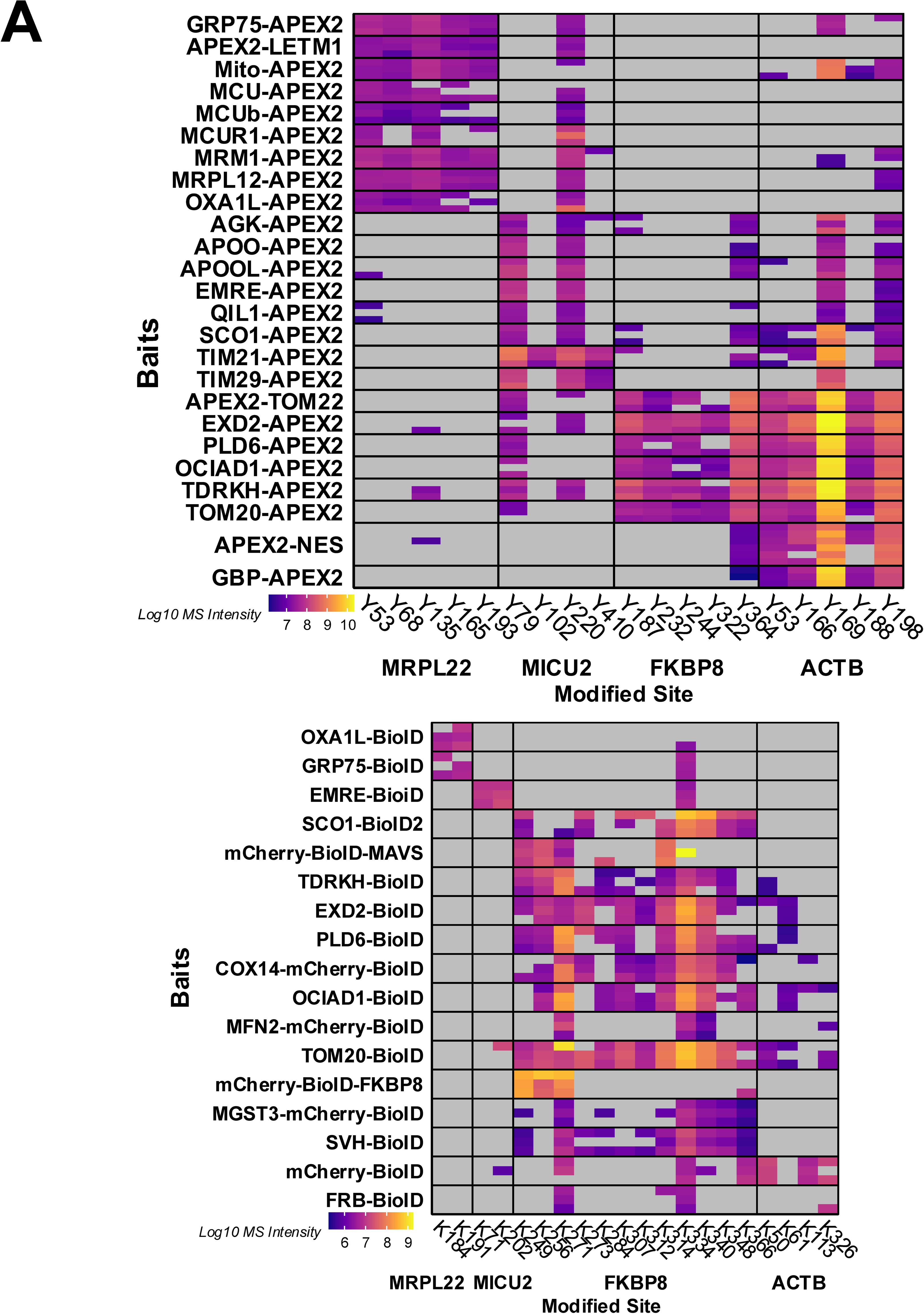
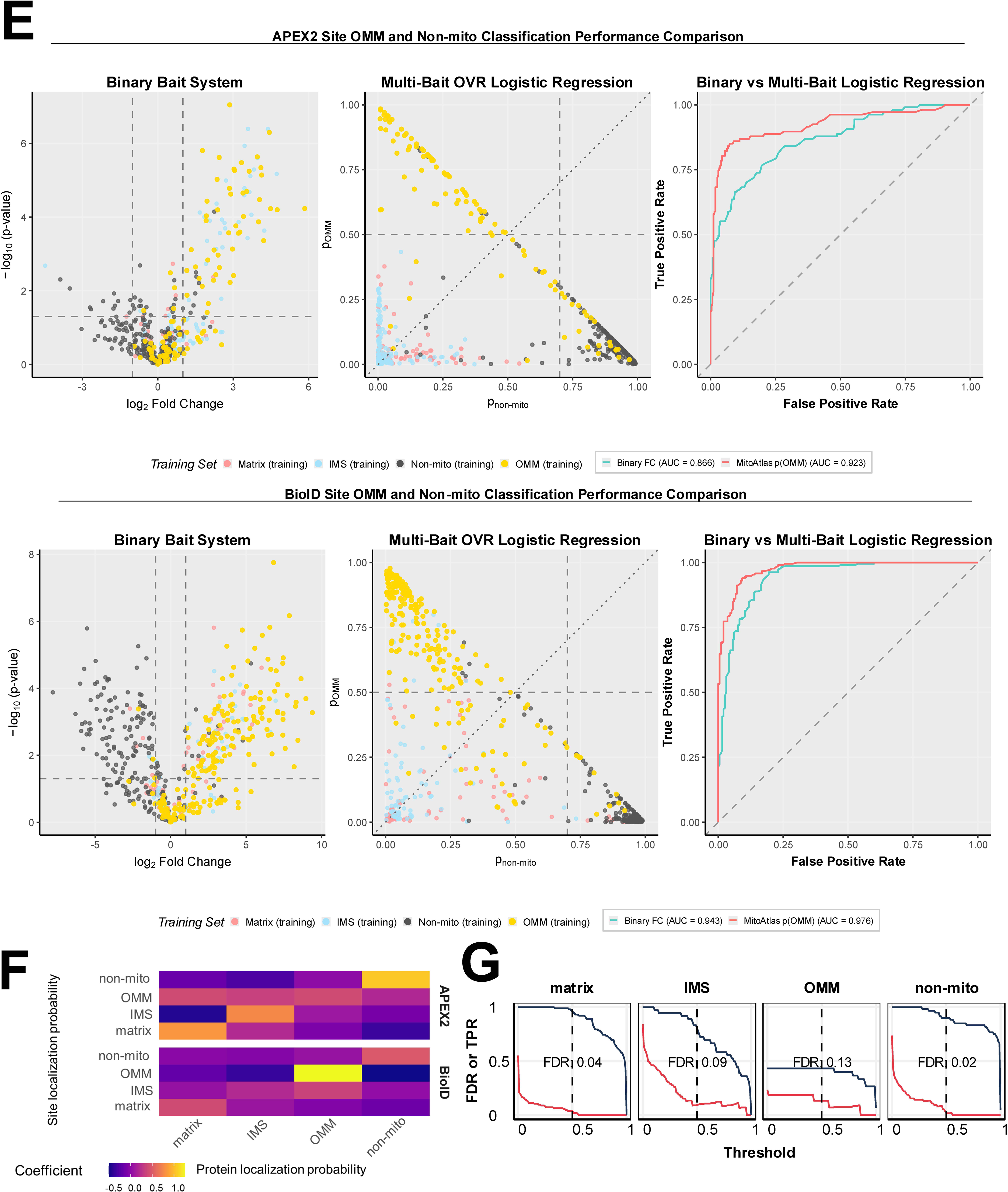
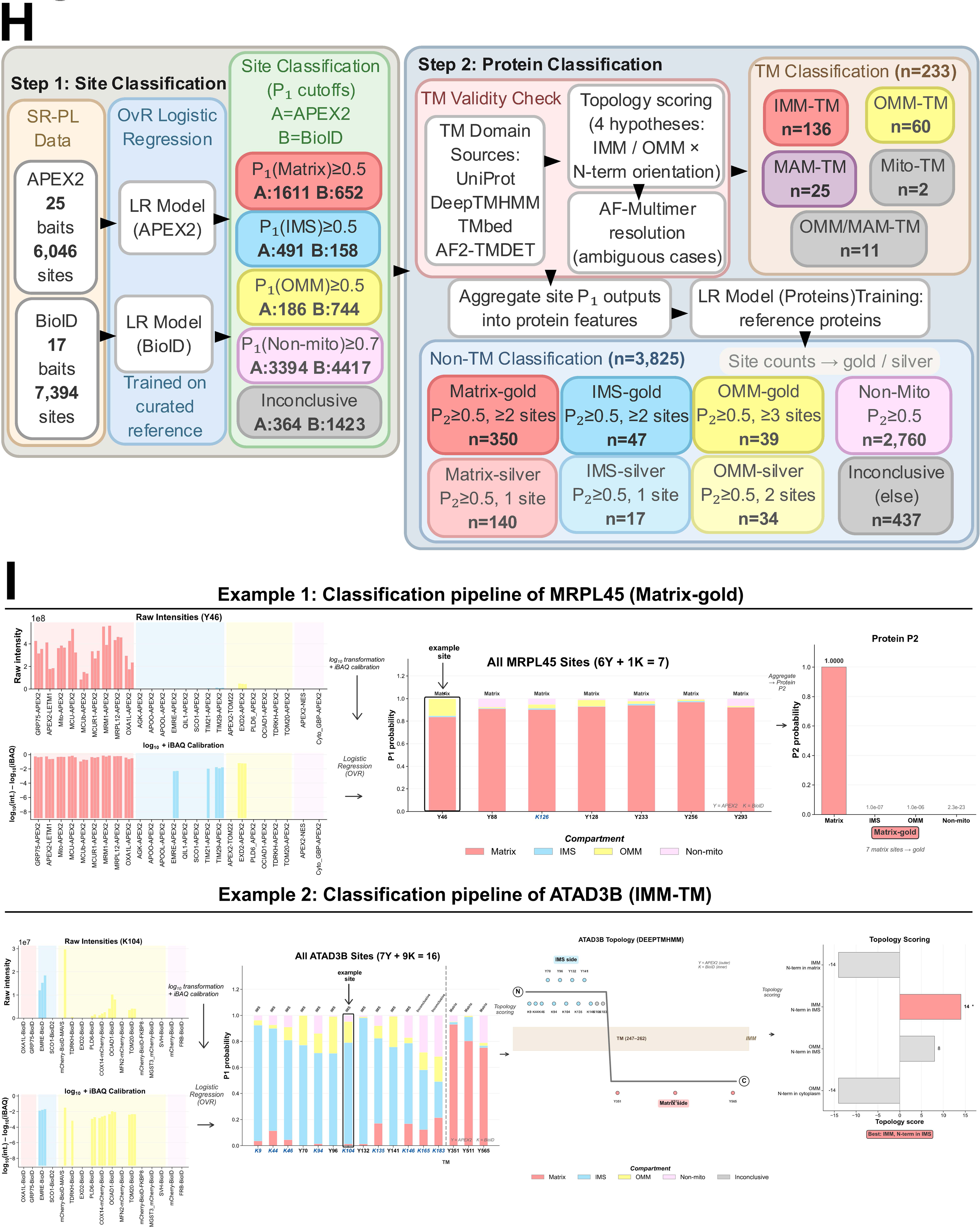
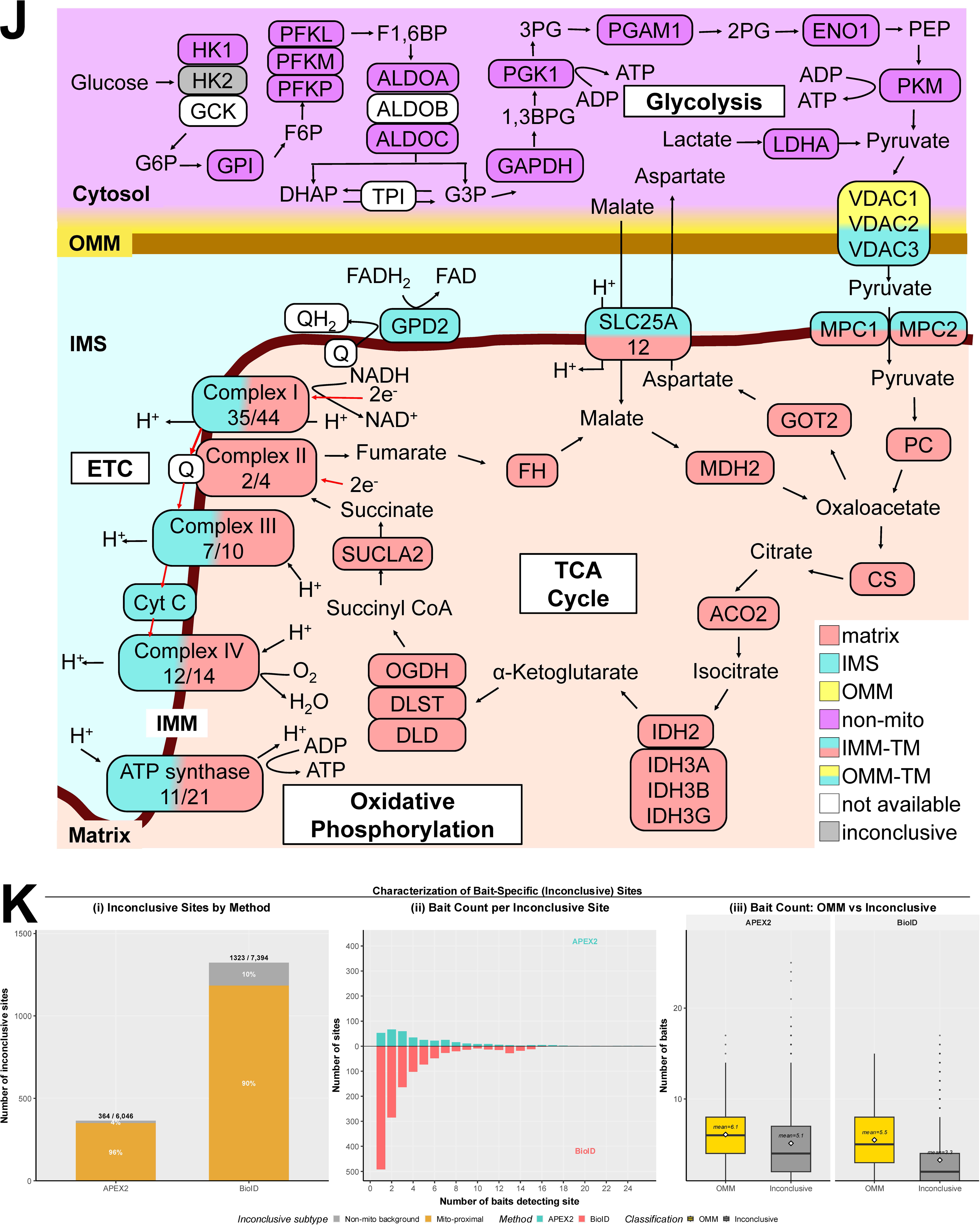
Site and Protein Classification Model Details (Related to. Figure 2). (A) Comparative MS intensity heatmaps of representative peptides containing DBP-modified tyrosine (top, APEX2 baits) and biotin-modified lysine (bottom, BioID baits) residues. Reference proteins include MRPL22 (Matrix), MICU2 (IMS), FKBP8 (OMM), and ACTB (Non-mitochondrial), demonstrating the compartment-specific enrichment patterns used as training features for the logistic regression model. (B) Comparison of 10-fold cross-validation accuracy for six supervised learning algorithms (CART, KNN, LDA, LR, NB, and SVM) trained separately on APEX2 (left) and BioID (right) datasets. Logistic Regression (LR) was selected for its high accuracy, interpretability, and tunable probability thresholds. (C) True Positive Rate (TPR) and False Discovery Rate (FDR) curves across a range of P1 probability thresholds for APEX2 (top) and BioID (bottom), evaluated separately for Matrix, IMS, OMM, and Non-mitochondrial compartments. Optimal thresholds were selected to maintain an overall FDR below 10%. (D) Visualization of the trained site-level logistic regression coefficient weights for each APEX2 bait (left) and BioID/BioID2 bait (right). Rows represent baits; columns represent the four sub-mitochondrial compartment predictions. Positive weights (yellow) indicate baits that serve as strong predictors for a specific localization. (E) Comparison of binary enrichment analysis and MitoAtlas multi-bait classification. Top: APEX2. Left: Volcano plot of log2 fold change (TOM20-APEX2 vs. APEX2-NES) versus –log10(p-value) for training-set biotinylation sites. OMM training sites (gold) cluster at positive fold changes, while non-mitochondrial training sites (dark gray) cluster near zero or negative values. Center: MitoAtlas classification probabilities for the same sites, plotted as p(non-mito) versus p(OMM), showing clear separation. Right: ROC curves comparing binary fold change (teal; AUC = 0.866) and MitoAtlas p(OMM) (red; AUC = 0.923) for discriminating 107 OMM from 269 non-mitochondrial training sites. Bottom: BioID. Left: Volcano plot of log2 fold change (TOM20-BioID vs. mCherry-BioID) versus –log10(p-value). Center: MitoAtlas probability scatter. Right: ROC curves comparing binary fold change (teal; AUC = 0.943) and MitoAtlas p(OMM) (red; AUC = 0.976) for discriminating 216 OMM from 195 non-mitochondrial training sites. Both comparisons demonstrate that the multi-bait MitoAtlas classifier consistently outperforms conventional binary enrichment analysis, particularly for the APEX2 dataset where the dual-bait approach shows substantial overlap between OMM and non-mitochondrial sites. (F) Visualization of the trained protein-level (P2) logistic regression coefficients used to aggregate site-level P1 outputs into protein-level localization probabilities. The heatmap shows the relative contribution of APEX2 and BioID site classifications for each compartment assignment. (G) Protein-level classification performance. True Positive Rate (TPR) and False Discovery Rate (FDR) curves across a range of P2 probability thresholds, evaluated separately for Matrix, IMS, OMM, and Non-mitochondrial compartments, demonstrating the accuracy of the second-step protein-level classifier. (H) Schematic overview of the two-step classification pipeline. Step 1 (Site Classification): raw MS intensities from 25 APEX2 and 17 BioID baits are log-transformed and iBAQ-calibrated, then classified by one-vs-rest (OvR) logistic regression into four sub-mitochondrial compartments (Matrix, IMS, OMM, and Non-mitochondrial), yielding a site-level probability (P_1_) for each of the 13,440 labeled sites. Step 2 (Protein Classification): proteins are first screened for transmembrane domains using four independent predictors (UniProt TM, TMbed, DeepTMHMM, AF2-TMDET). The 233 TM proteins are routed to a topology scoring pipeline that integrates site-level classifications with predicted TM domain boundaries to assign membrane orientation (IMM-TM, OMM-TM, MAM-TM). The remaining 3,825 non-TM proteins are classified by aggregating site-level P1 outputs into an 8-dimensional protein-level feature vector and applying a second OvR LR model, with results assigned to tiered confidence categories (Gold/Silver) based on P_2_ probability and the number of supporting sites. (I) Worked examples of the classification pipeline. Left: non-TM protein MRPL45 (Matrix-gold). Raw MS intensities across all baits for a representative tyrosine site (Y46), followed by log-transformation and iBAQ calibration. OvR logistic regression assigns P_1_ probabilities to all seven MRPL45 sites (6 Tyr + 1 Lys), with all sites classified as Matrix (one site, K126, shown in yellow, falls below the confidence threshold). Aggregation of site-level P_1_ values into a protein-level P_2_ probability yields a high-confidence Matrix-gold assignment. Right: TM protein ATAD3B (IMM-TM). Raw MS intensities and log-transformed/iBAQ-calibrated values for a representative lysine site (K104) across all BioID baits. P_1_ site-level classification of all 16 ATAD3B sites (7 Tyr + 9 Lys) reveals a mixture of Matrix and IMS assignments, characteristic of an IMM-spanning protein. Topology scoring integrates site localizations with DeepTMHMM transmembrane domain predictions and AlphaFold structural models to determine the membrane orientation, assigning ATAD3B as IMM-TM with its N-terminus facing the matrix. (J) Metabolic pathway map including glycolysis, the TCA cycle, and the electron transport chain (OXPHOS), with enzymes and complexes color-coded by their MitoAtlas sub-mitochondrial localization assignments. Numbers beneath OXPHOS complexes indicate the fraction of subunits detected by MitoAtlas. (K) Characterization of inconclusive sites. Sites that failed to meet the multi-bait consensus threshold for any compartment (p(matrix) < 0.5, p(IMS) < 0.5, p(OMM) < 0.5, and p(non-mito) < 0.7) were designated inconclusive. (i) Stacked bar chart showing the number of inconclusive sites by method: APEX2 (364 of 6,046 sites, 6.0%) and BioID (1,323 of 7,394 sites, 17.9%), split by subtype. Mito-proximal inconclusive sites (gold) were detected by at least one mitochondrial bait but lacked multi-bait consensus; non-mito background sites (gray) were detected exclusively by cytosolic control baits. For APEX2, 96% of inconclusive sites (350 of 364) are mito-proximal; for BioID, 90% (1,185 of 1,323), indicating that the vast majority represent biologically relevant mitochondria-proximal labeling events rather than cytosolic background. (ii) Mirrored histogram of the number of baits detecting each inconclusive site (APEX2 upward, BioID downward). For BioID, 37% of inconclusive sites (492 of 1,323) were detected by only a single bait, consistent with contact-dependent bait–prey interactions rather than compartment-resident labeling. (iii) Box plots comparing the number of detecting baits for confidently classified OMM sites versus inconclusive sites, faceted by method. OMM-classified sites are detected by significantly more baits (BioID: median 5 vs. 2; APEX2: median 6 vs. 4), confirming that multi-bait consensus underlies confident spatial assignment.

**Figure S3.**
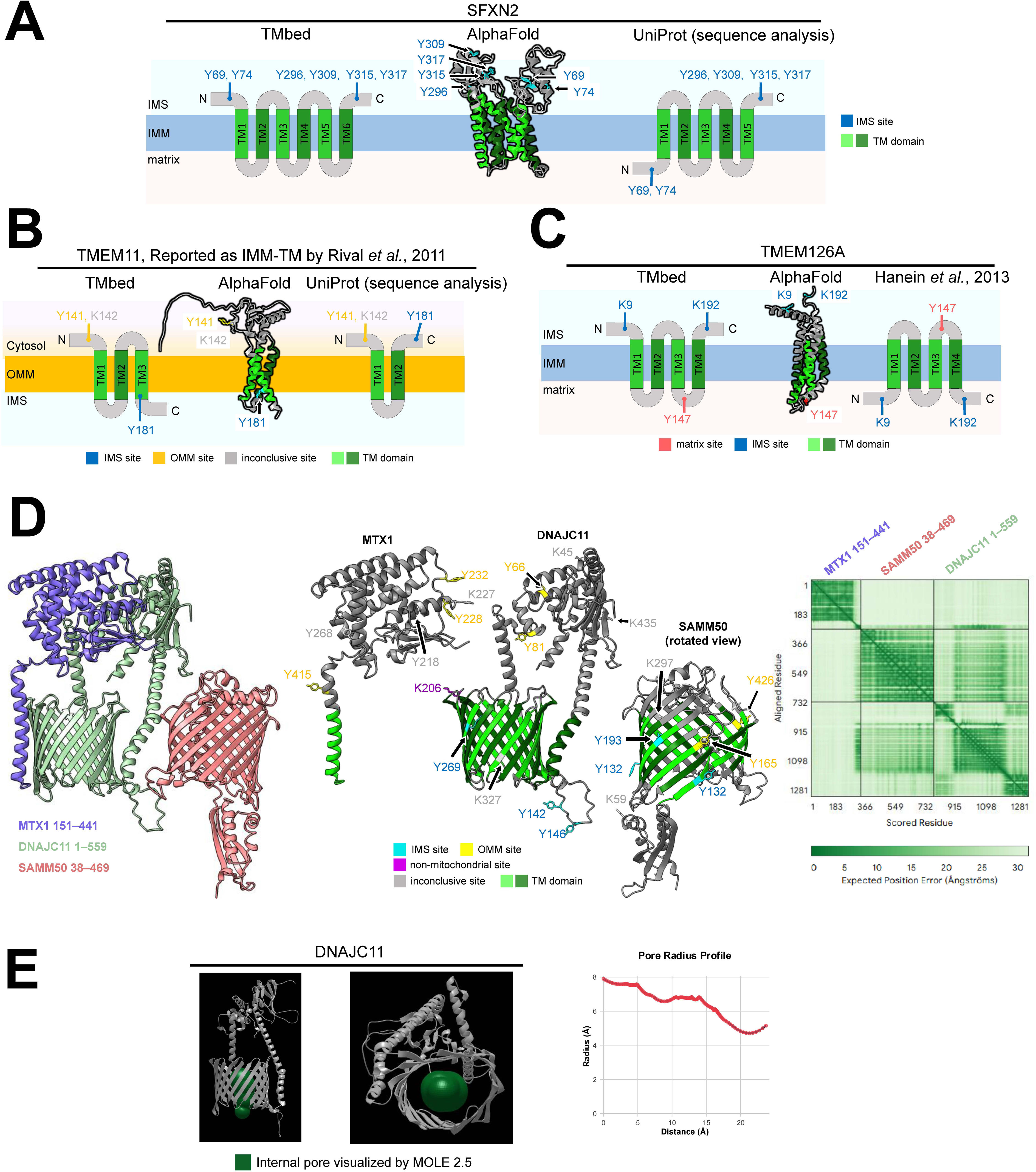
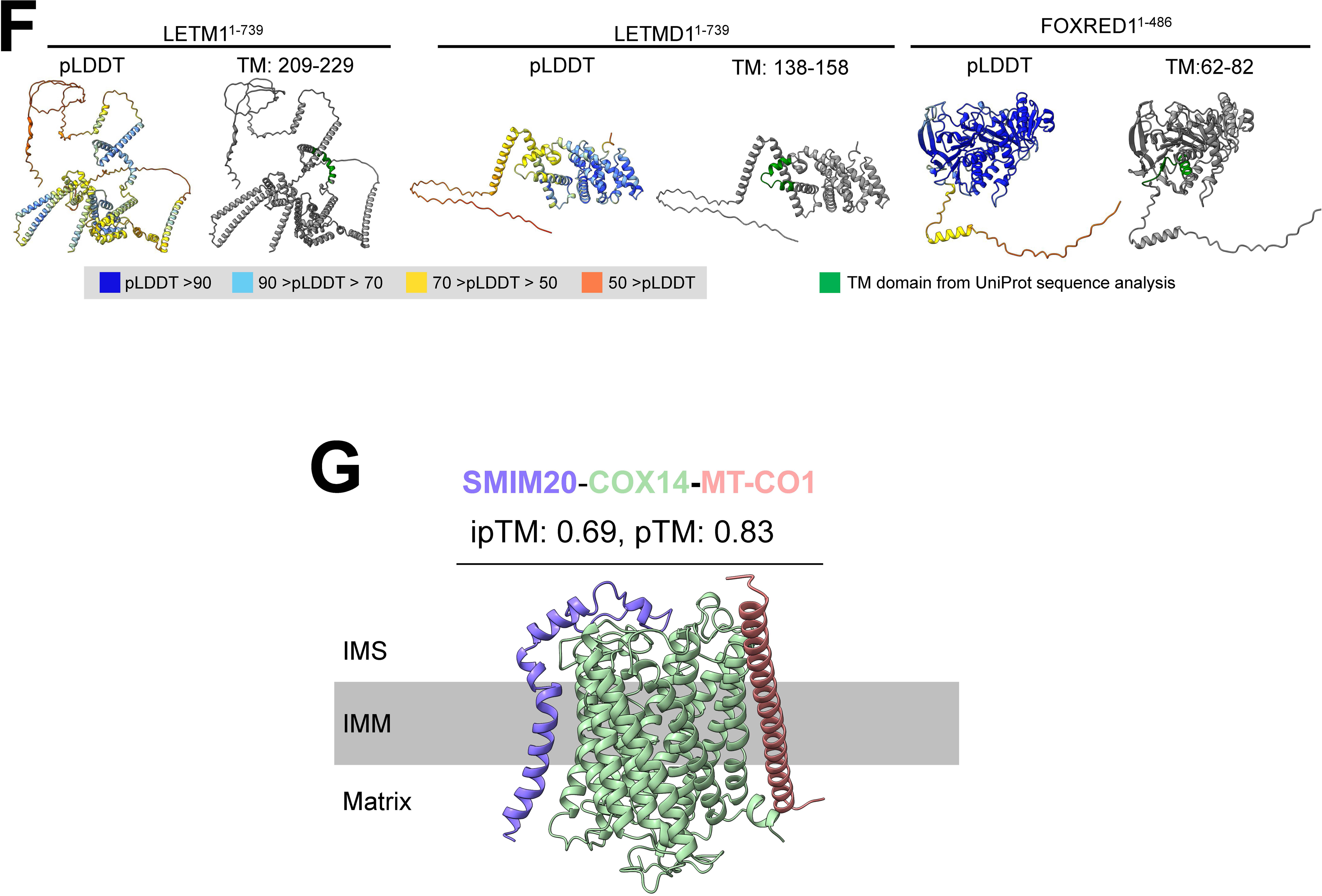
Validation and Structural Refinement of Mitochondrial Membrane Topologies (related to. Figure 3). (A) Revised 6-TM architecture of SFXN2. Comparison of the transmembrane (TM) domain count and topological orientation of the IMM protein SFXN2. The MitoAtlas data, integrated with the TMbed model (left) and AlphaFold2 (AF2) structure (middle), supports a 6-TM model, contradicting the 5-TM model currently annotated in UniProt (right). (B) Reclassification and topology of TMEM11. Comparison of TM domain predictions for TMEM11. MitoAtlas and AF2 evidence (left, middle) demonstrate that TMEM11 is an OMM protein with a 3-TM architecture, challenging the UniProt annotation of a 2-TM IMM protein (right). (C) Orientation of the TMEM126A N-terminus. Topological analysis of the IMM protein TMEM126A. While UniProt (right) suggests a matrix-facing N-terminus based on historical protease protection assays, the integration of MitoAtlas SR-PL data with TMbed and AF2 (left, middle) indicates an intermembrane space (IMS) orientation. (D) Structural modeling of the MTX1-DNAJC11-SAMM50 complex. AlphaFold3 (AF3) predicted model of the OMM complex (left), with SR-PL labeling sites projected onto the structure (middle). The Predicted Alignment Error (PAE) matrix (right) indicates the relative orientations of the subunits. (E) Pore analysis of the DNAJC11 β-barrel. Visualization of the internal pore of DNAJC11 (green) generated via MOLE 2.5 software from side (left) and bottom (middle) perspectives. The corresponding pore radius profile is shown on the right. (F) Reclassification of UniProt-predicted TM proteins to the matrix. AF2 structures of proteins with legacy TM annotations in UniProt that were reclassified as matrix-resident proteins in MitoAtlas. Structures are colored by pLDDT confidence scores, with UniProt-predicted TM domains highlighted in green to illustrate their lack of characteristic transmembrane structural features. (G) AlphaFold3-Multimer predicted structure of the SMIM20–COX14–MT-CO1 complex, rendered in ChimeraX (ipTM = 0.69, pTM = 0.83).

**Figure S4.**
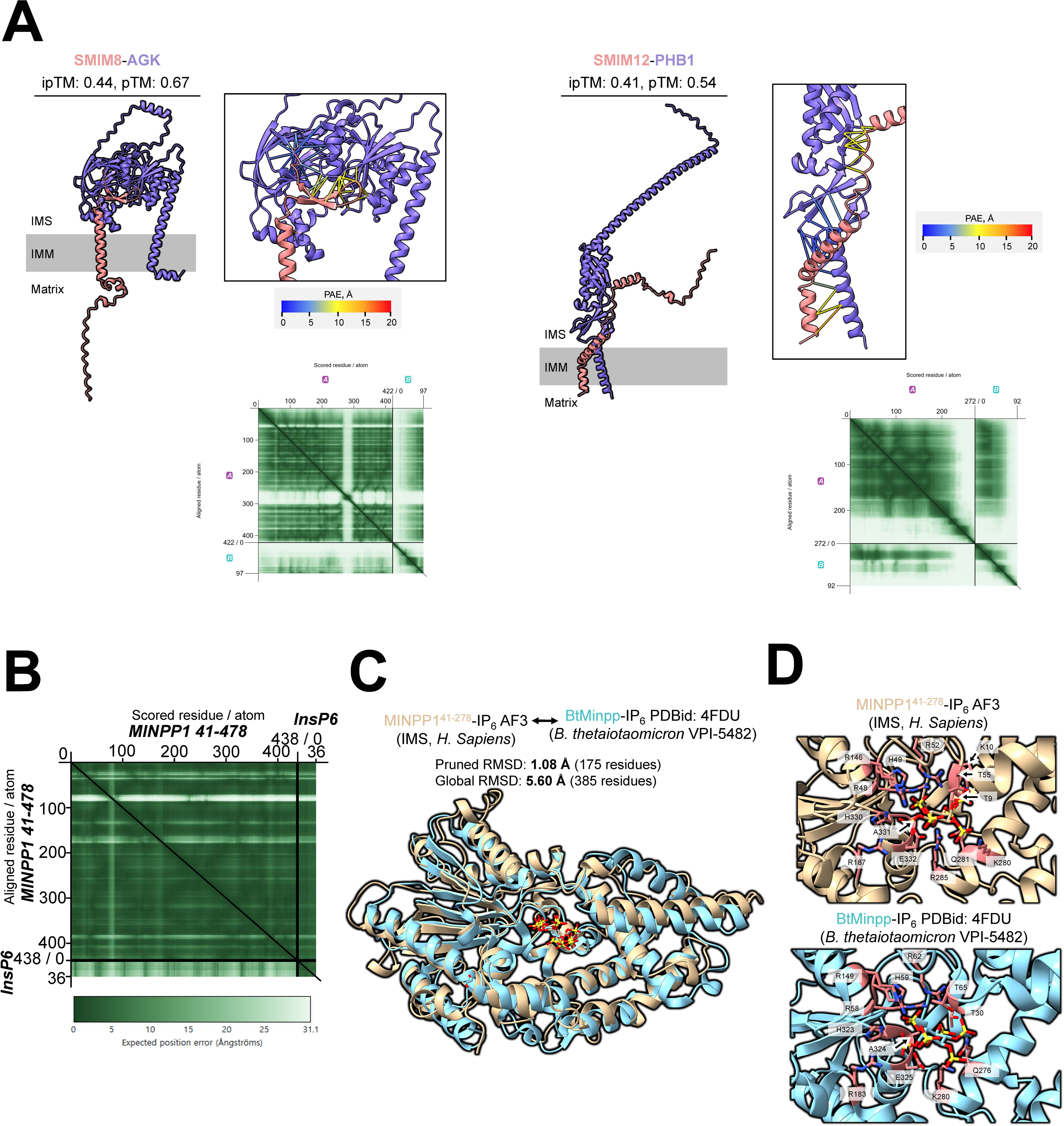
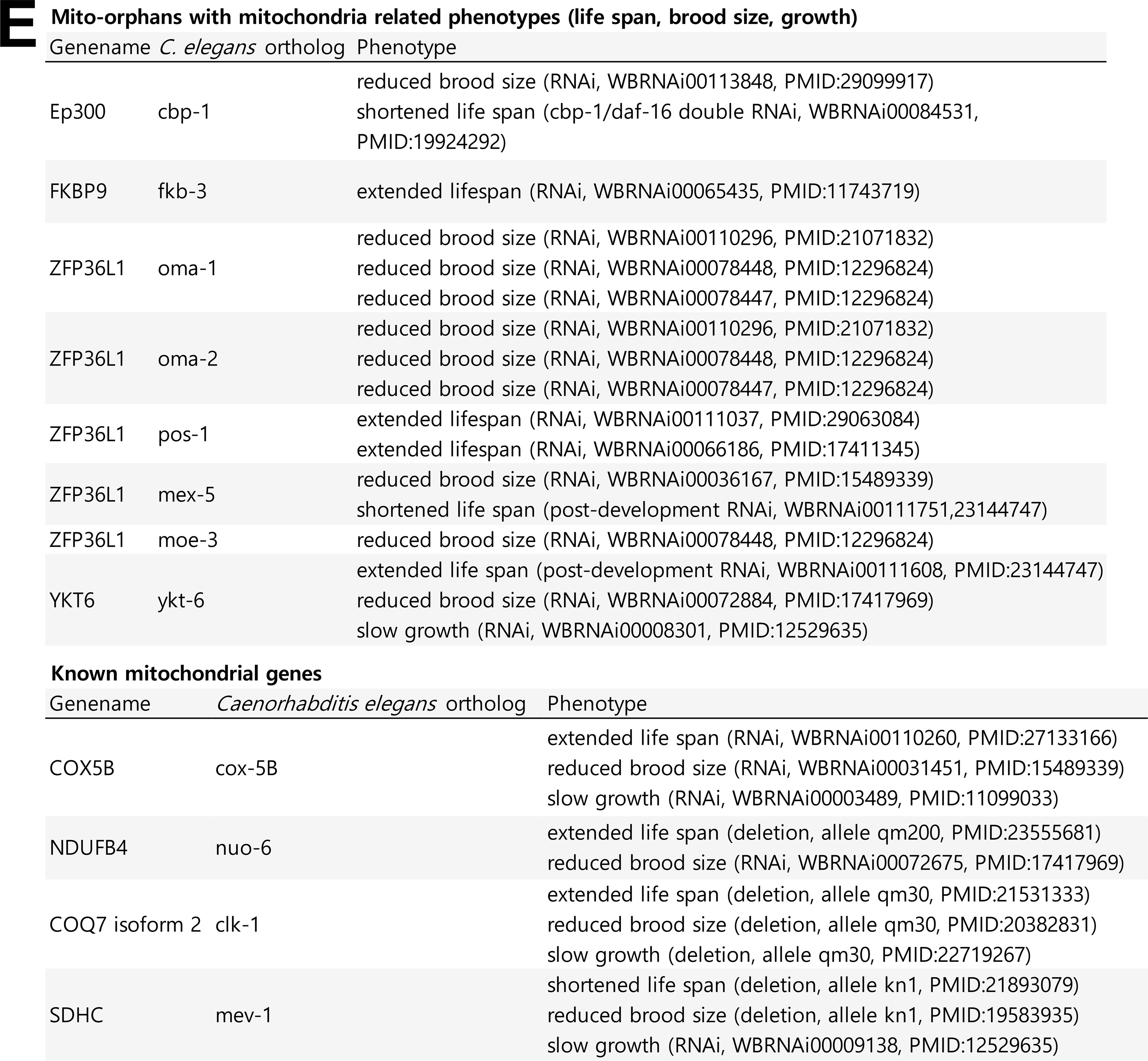
Structural Validation and Functional Analysis of Mito-Orphan Proteins (related to. Figure 4). (A) AlphaFold3-Multimer predicted structures for SMIM8–AGK and SMIM12–PHB1 complexes, supporting the IMM-TM localization of SMIM8 and SMIM12. Predicted Aligned Error (PAE) matrices and interface confidence scores (ipTM, pTM) are shown for each complex. (B) Structural alignment of MINPP1 (residues 41–478) with InsP6 substrate, showing the predicted binding site. The distance matrix (bottom) and PAE plot indicate high confidence in the core catalytic domain structure. (C) Structural superposition of MINPP1 AlphaFold3 model (human, IMS) with the bacterial homolog BtMinpp (B. thetaiotaomicron, PDB: 4FDU), showing a pruned RMSD of 1.08 Å over 175 aligned residues, highlighting the conserved histidine phosphatase fold despite divergent disulfide bond content. (D) Active site comparison between human MINPP1 (top) and bacterial BtMinpp (bottom), with key catalytic residues labeled. The conserved catalytic architecture supports enzymatic activity of MINPP1 in the IMS. (E) Table of mito-orphan orthologs in *C. elegans* showing mitochondria-related phenotypes upon RNAi knockdown (top), compared with known mitochondrial genes exhibiting similar phenotypes (bottom). Phenotypes include reduced brood size, altered lifespan, and slow growth, supporting the functional relevance of the newly identified mito-orphans.

**Figure S5.**
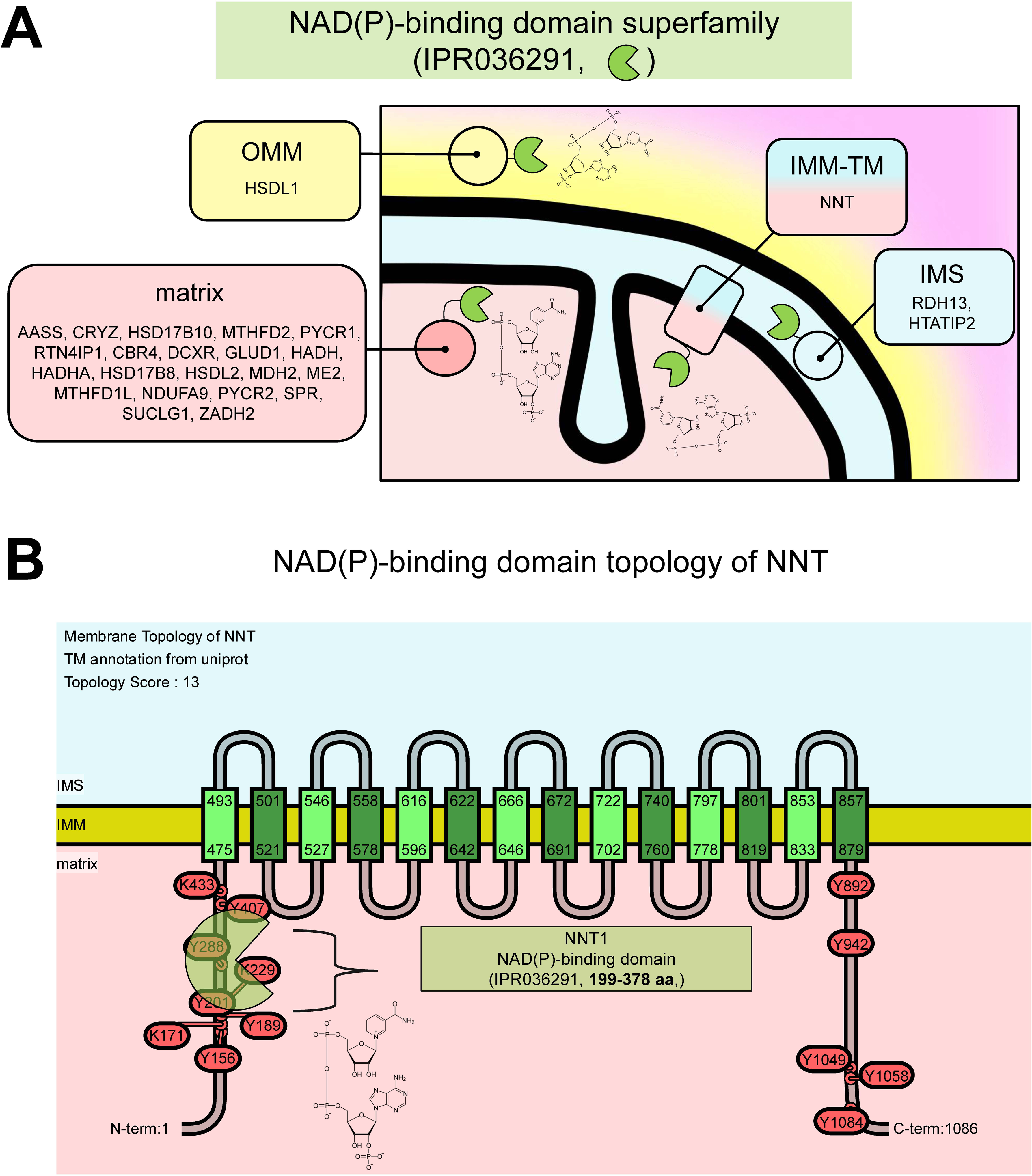
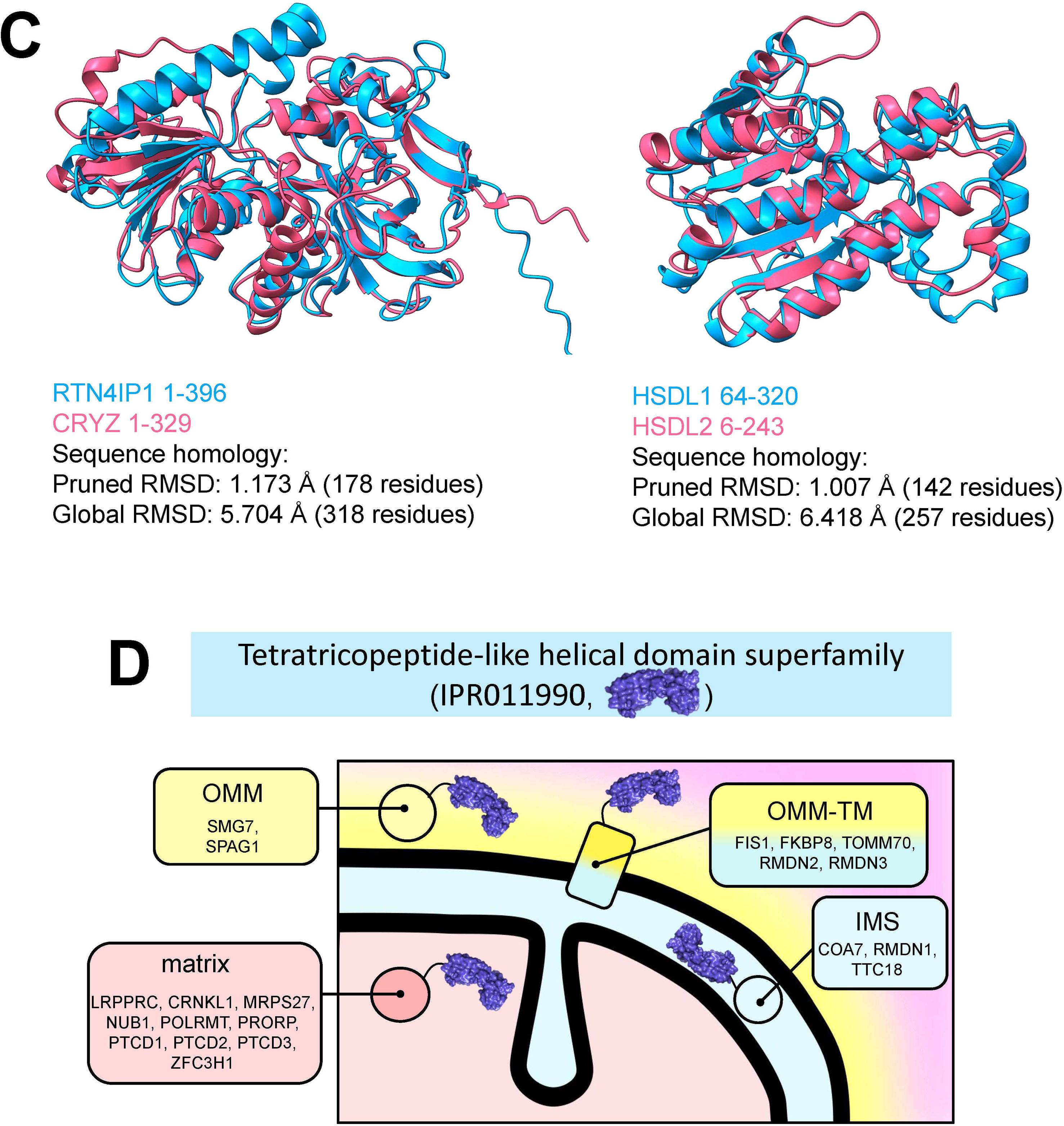
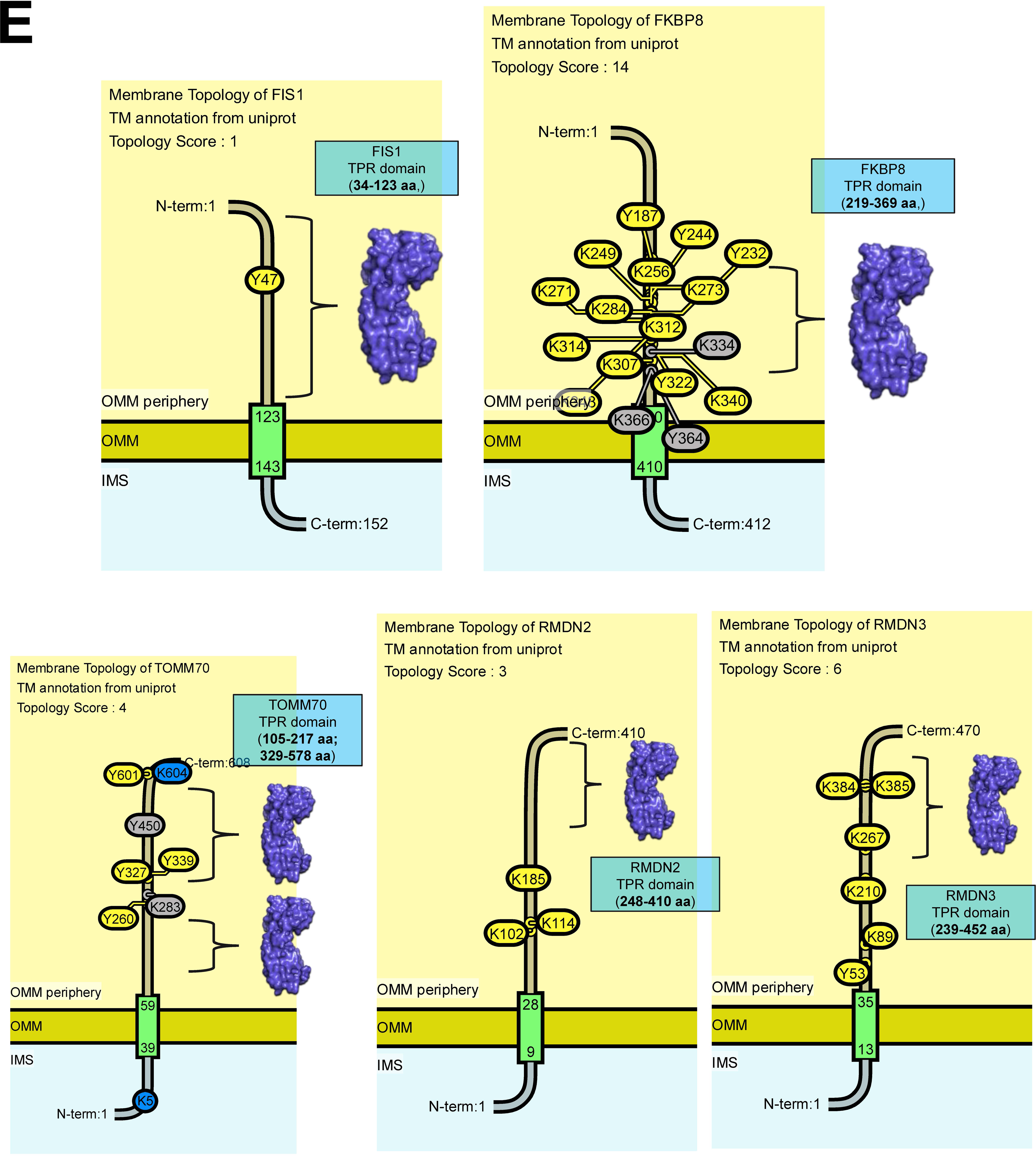
Spatial Enrichment and Membrane Topology of NAD(P)-binding and Tetratricopeptide Helical Domain Superfamilies. (related to. Figure 5) (A) Sub-mitochondrial localization profiles for all members of the NAD(P)-binding superfamily, indicating a predominant matrix enrichment. (B) Refined topological model of the inner mitochondrial membrane (IMM) protein Nicotinamide Nucleotide Transhydrogenase (NNT), illustrating the orientation of its NAD(P)-binding domains relative to the matrix and IMS. (C) AlphaFold2-based structural alignments of representative protein pairs RTN4IP1-CRYZ and HSDL1-HSDL2 using ChimeraX matchmaker, demonstrating conserved folding patterns despite divergent primary sequences. Sequence homology was calculated using LALIGN. (D) Sub-mitochondrial distribution of the Tetratricopeptide-like (TPR) helical domain superfamily. (E) Membrane topology models for OMM-resident proteins FIS1, FKBP8, TOMM70, RMDN2, and RMDN3, depicting the cytosolic exposure of their respective tetratricopeptide repeat domains.

**Figure S6.**
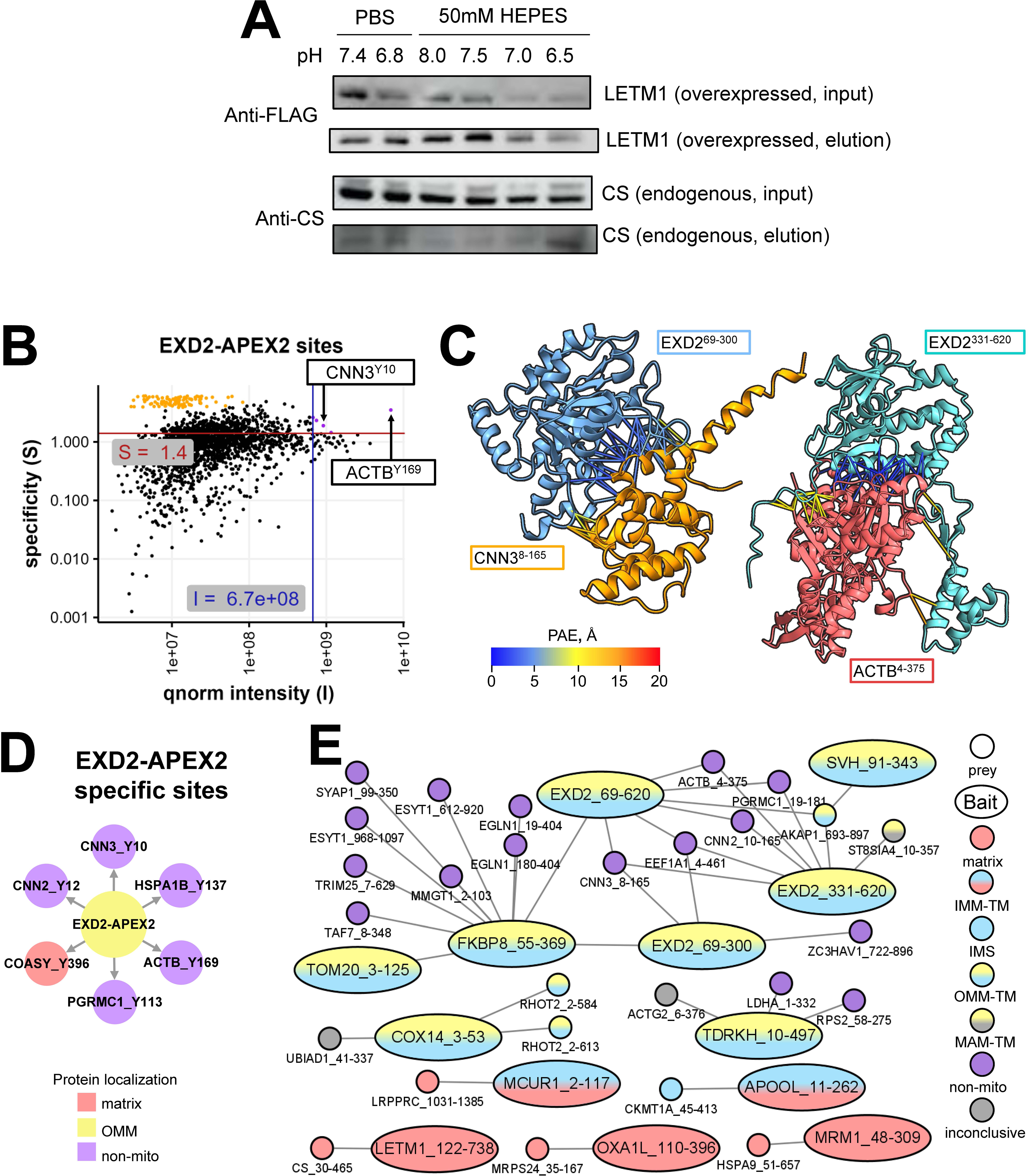
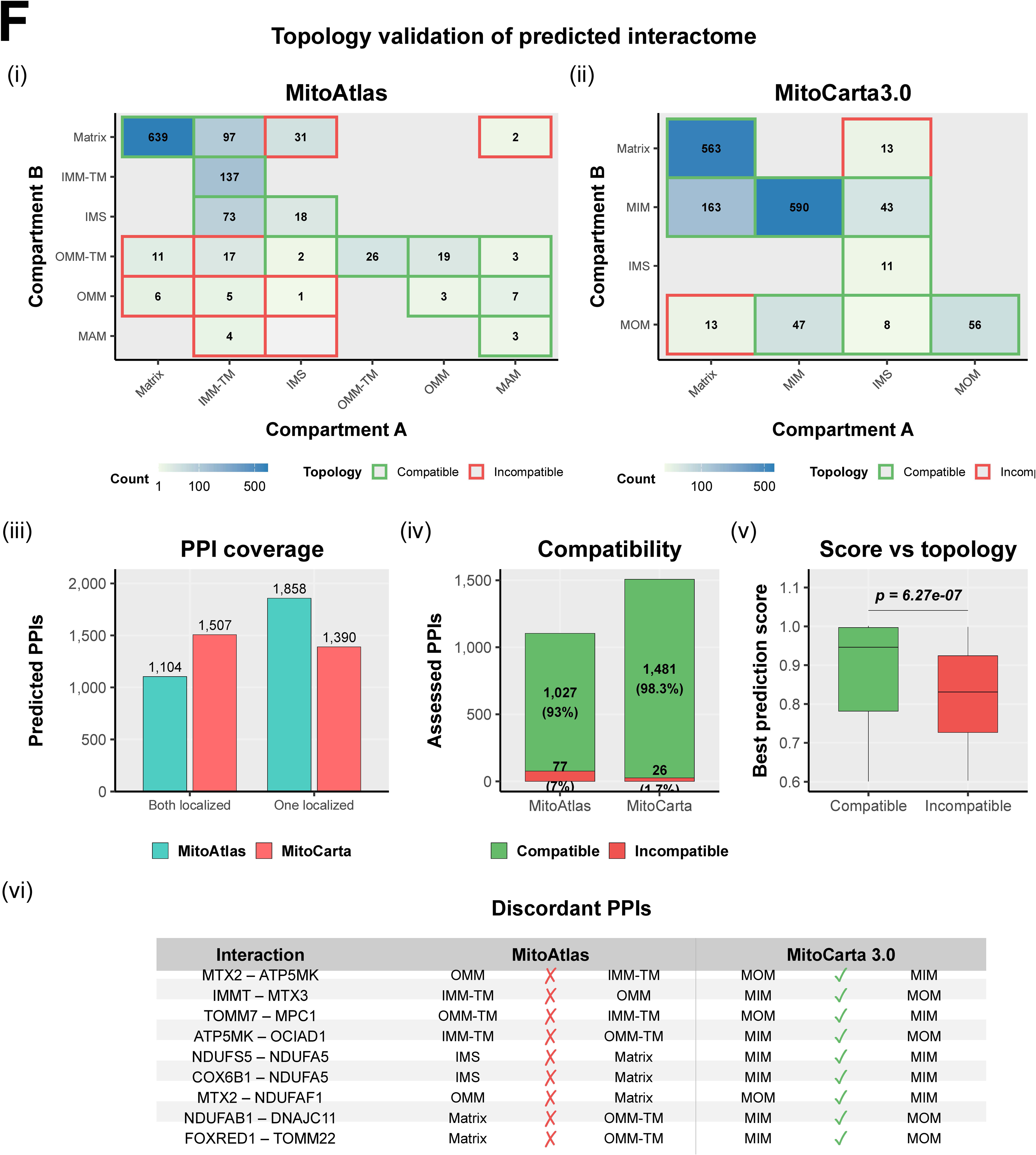
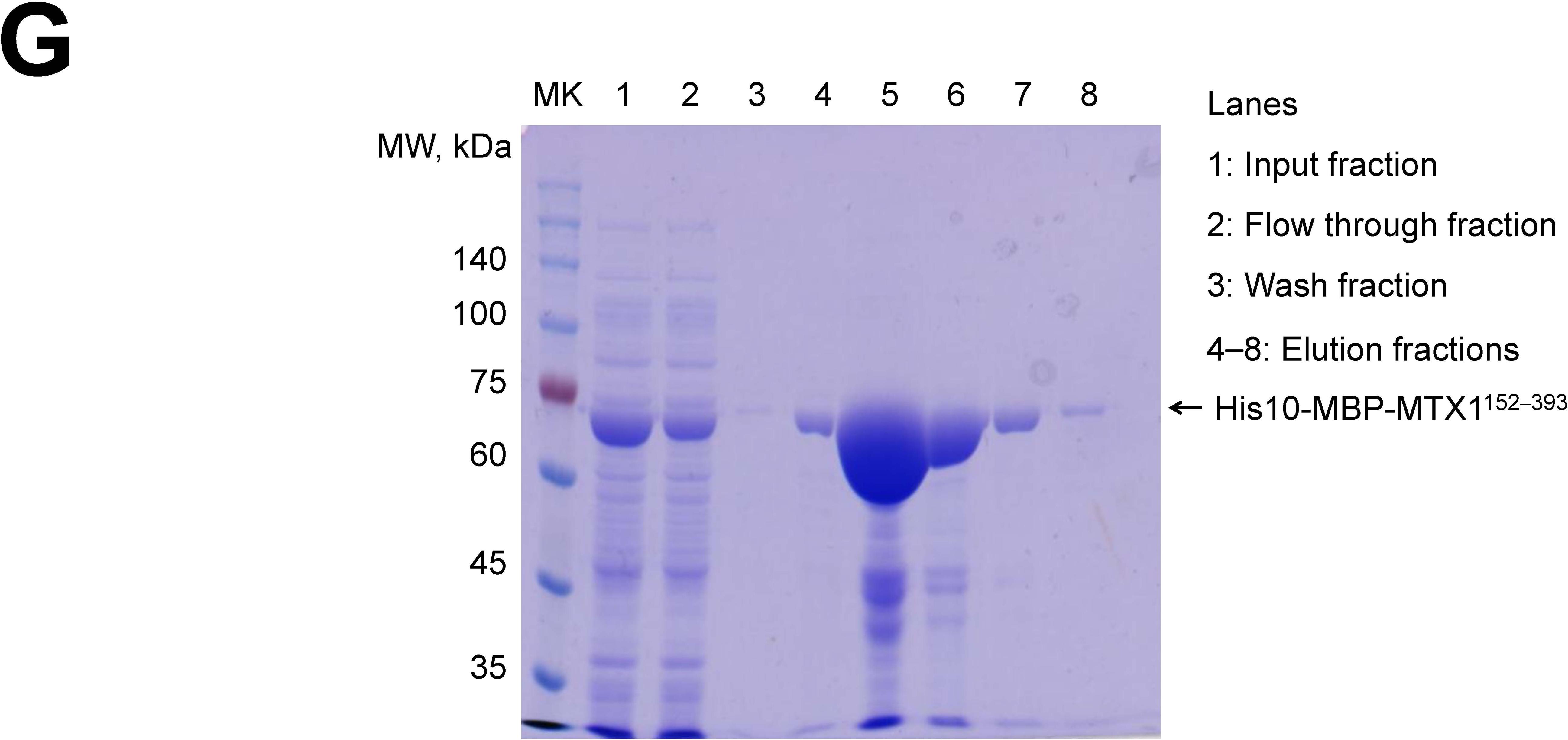
CoIP optimization, specific bait–prey site analysis, and topology validation of predicted interactome (related to. Figure 6). (A) Optimization of co-immunoprecipitation (CoIP) lysis conditions for the LETM1–CS interaction. Six buffer conditions containing 1% NP-40 were tested: PBS at pH 7.4 and 6.8 (left), and 50 mM HEPES at pH 8.0, 7.5, 7.0, and 6.5 (right). Four western blots are shown (top to bottom): anti-FLAG input (overexpressed MTS-3×FLAG-LETM1), anti-FLAG elution, anti-CS input (endogenous), and anti-CS elution. PBS pH 7.4 with 1% NP-40 was selected as the optimal condition for the main figure CoIP experiment (Figure 6F). (B) Quantile normalized intensity and specificity of EXD2-APEX2 labeled sites. (C) AF2-Multimer prediction of the EXD2 N-terminus-domain (NTD) and CNN3 complex and EXD2 C-terminus-domain (CTD) and ACTB complex. (D) Network graph representation of all EXD2-APEX2 specific bait–prey sites identified by the intensity and specificity filtering pipeline. (E) Network graph representation of all MitoAtlas baits connected to their specific bait–prey sites, showing the complete landscape of high-confidence proximity interactions across the 42-bait library. (F) Validation of computationally predicted mitochondrial protein–protein interactions using MitoAtlas sub-mitochondrial topology. (i) Coverage: grouped bar chart showing the number of predicted PPIs (from 29,257 total^27^) assessable by MitoAtlas (1,104) versus MitoCarta3.0 (1,507). (ii) Compartment co-occurrence heatmap of MitoAtlas-assessable interactions across sub-mitochondrial compartment pairs (Matrix, IMM-TM, IMS, OMM-TM, OMM, MAM). Green borders indicate topologically compatible pairs; red borders indicate incompatible pairs in non-adjacent compartments. (iii) Compartment co-occurrence heatmap using MitoCarta3.0’s coarser four-compartment scheme (Matrix, MIM, IMS, MOM). (iv) Topology compatibility rate: stacked bar chart comparing the fraction of assessed PPIs that are topologically compatible versus incompatible for MitoAtlas (7.0% incompatible, 77 pairs) versus MitoCarta3.0 (1.7% incompatible, 26 pairs). MitoAtlas’s finer granularity, particularly the distinction between OMM-TM and peripheral OMM proteins, enables more stringent spatial filtering. (v) Box plots comparing computational prediction confidence scores (best of AFprob, CFprob, AFMprob) between topologically compatible and incompatible interactions. Incompatible interactions have significantly lower prediction scores (median 0.831 vs. 0.946; Wilcoxon p = 6.27 × 10⁻⁷), providing reciprocal validation of both MitoAtlas topology and the computational predictions. (vi) Representative table of PPIs that pass MitoCarta3.0’s coarser topology check but are flagged as incompatible by MitoAtlas, demonstrating the added value of domain-level sub-mitochondrial resolution for interactome filtering. (G) Coomassie staining result of immobilized metal affinity chromatography of recombinant protein His10-MBP-TEVs-MTX1^152–393^ expressed in E. coli.

**Figure S7.**
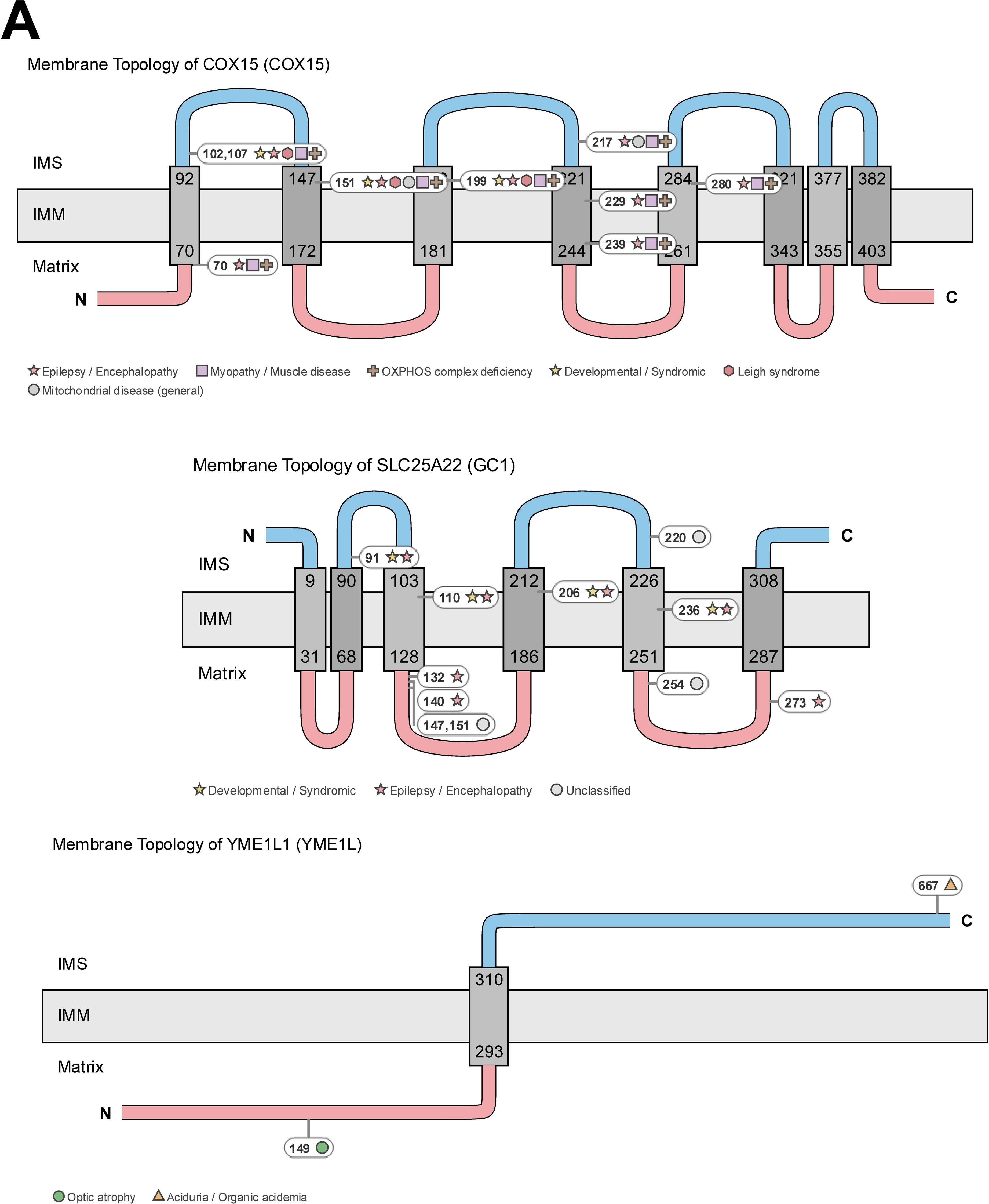
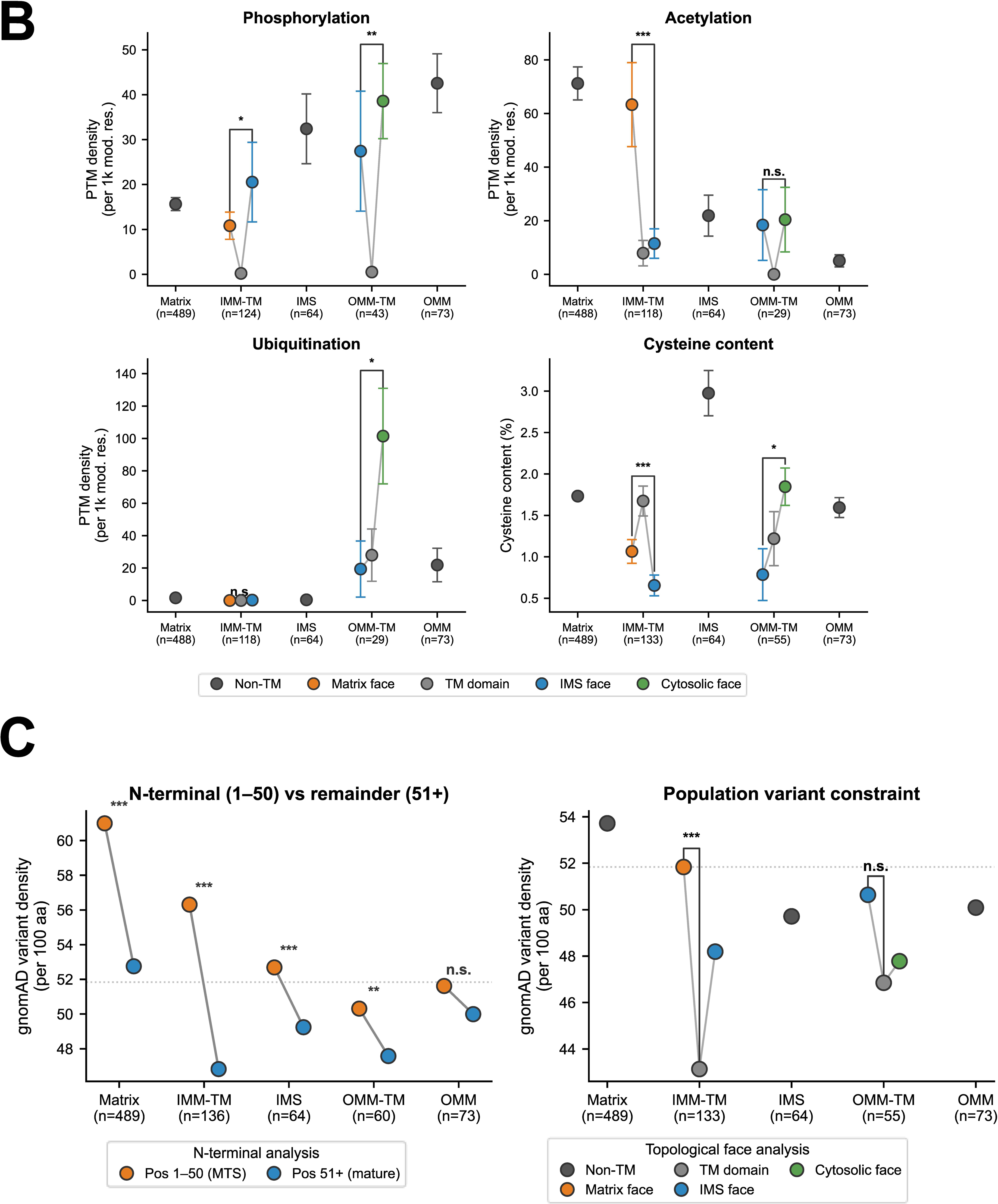
Application Scenarios: Enhanced Biological Discovery with MitoAtlas Site-Resolved Spatial Data (related to. Figure 7). (A) Disease variant interpretation: additional examples of topology-dependent disease patterns. Top: COX15 (UniProt Q7KZN9), a heme A farnesyltransferase with eight TM helices (N-terminus in matrix). Nine ClinVar pathogenic variants are mapped. All are associated with fatal infantile cardioencephalomyopathy due to cytochrome c oxidase deficiency type 2 (MIM #615119); positions 102 and 199 are additionally linked to Leigh syndrome. Variants are distributed across TM helices (pos 70, 151, 199, 229, 239, 280) and IMS-facing loops (pos 102, 107, 217) with no clear topology-dependent segregation, consistent with a uniformly essential role of COX15 in heme A biosynthesis. Middle: SLC25A22 (GC1; UniProt Q9H936), an IMM carrier with six TM helices (N-terminus in IMS). Eleven variants are mapped, all associated with early-onset epileptic encephalopathy (DEE type 3, MIM #609304; early myoclonic encephalopathy). Variants span IMS (pos 91, 220), TM helices (pos 110, 206, 236), and matrix-facing loops (pos 132, 140, 147, 151, 254, 273). Matrix-facing variants (6/11) tend to present as early myoclonic encephalopathy, while IMS/TM variants are more frequently annotated as DEE type 3. Bottom: YME1L1 (i-AAA protease; UniProt Q96TA2), a single-pass IMM protein with a large IMS-facing AAA+ protease domain and matrix-facing N-terminal domain. Two pathogenic variants show clear topology-dependent disease segregation: position 149 (matrix-facing) causes optic atrophy 11 (MIM #617302), while position 667 (IMS-facing catalytic domain) causes 3-methylglutaconic aciduria (MIM #617248). (B) PTM density and cysteine content across mitochondrial compartments and membrane faces. PTM site density and amino acid composition were analyzed across five compartments: matrix (non-TM, n=489), IMM-TM (n=134), IMS (non-TM, n=64), OMM-TM (n=57), and OMM (non-TM, n=73). For TM proteins, residues and PTM sites were assigned to membrane faces based on MitoAtlas topology. PTM annotations from UniProt. Dots, mean; error bars, SEM. Gray lines connect three faces within each TM group. Brackets, one-way ANOVA (*p<0.05, **p<0.01, ***p<0.001). First panel, phosphorylation (sites/1,000 S/T/Y): TM domains show near-zero phosphorylation in both IMM (0.2/1k) and OMM (0.5/1k). Within IMM-TM, IMS-facing loops of IMM-TM proteins have the highest density (20.5/1k vs. matrix 10.8/1k, TM 0.2/1k; p=0.031). OMM-TM cytosol-facing surfaces are most modified (38.6/1k; p=0.004) across all faces. Second panel, acetylation (N6-acetyllysine/1,000 K): IMM-TM matrix-facing loops are enriched (63.3/1k vs. TM 7.9/1k, IMS 11.5/1k; ANOVA ***), consistent with matrix acetyl-CoA availability and SIRT3 deacetylation. OMM-TM shows minimal face-dependent variation (n.s.). Third panel, ubiquitination (sites/1,000 K): absent from IMM-TM faces (0-0.2/1k; n.s.). OMM-TM cytosol-facing domains show elevated ubiquitination (101.5/1k vs. TM 28.0/1k, IMS 19.4/1k; p=0.049), reflecting cytosolic ubiquitin-proteasome access. Fourth panel, cysteine content (%): IMM-TM faces differ significantly (ANOVA p=1.4×10⁻⁵); matrix faces 1.07%, TM 1.67%, IMS faces 0.65% (matrix vs. IMS Welch p=0.032). OMM-TM faces also differ (p=0.040); cytosol-facing 1.85%, TM 1.22%, IMS 0.79%. Soluble IMS proteins are cysteine-enriched (2.98% vs. matrix 1.73%, OMM 1.59%), consistent with the Mia40/CHCHD4 disulfide relay pathway. (C) gnomAD population variant density across mitochondrial compartments. Missense variant density was analyzed across 822 MitoAtlas proteins in five compartments: matrix (non-TM, n=489), IMM-TM (n=122–134), IMS (non-TM, n=64), OMM-TM (n=55–57), and OMM (non-TM, n=73); ranges reflect per-face protein counts. Variant positions from gnomAD via EBI Proteins API. Overall density: 51.8 variants/100 aa (182,774 unique missense positions across 352,895 residues). Dashed line, overall mean. Brackets, binomial test (TM vs. non-TM; ***p<0.001). Left, N-terminal (pos 1–50) vs. remainder: N-terminal residues, encompassing the mitochondrial targeting sequence in most matrix and IMM proteins, show higher variant density across all compartments (matrix 61.0 vs. 52.8/100aa, ***; IMM-TM 56.3 vs. 46.8, ***; IMS 52.7 vs. 49.3, ***; OMM-TM 50.3 vs. 47.6, **; OMM 51.6 vs. 50.0, n.s.), indicating reduced constraint on the cleaved MTS. Right, density per topological face: IMM-TM transmembrane domains are depleted in variants (43.1/100aa) compared to matrix-facing (51.8) and IMS-facing (45.0) loops (binomial p=8.0×10⁻³³, ***). OMM-TM shows a similar but non-significant trend (TM 46.9 vs. non-TM 47.8–50.6; p=df0.20). Soluble matrix proteins show the highest overall density (53.7/100aa).

